# *Salmonella* Typhimurium outer membrane protein A (OmpA) renders protection against nitrosative stress by promoting SCV stability in murine macrophages

**DOI:** 10.1101/2021.02.12.430987

**Authors:** Atish Roy Chowdhury, Shivjee Sah, Umesh Varshney, Dipshikha Chakravortty

## Abstract

Porins are highly conserved bacterial outer membrane proteins involved in the selective transport of charged molecules across the membrane. Despite their significant contributions to the pathogenesis of Gram-negative bacteria, their precise role in salmonellosis remains elusive. In this study, we investigated the role of porins (OmpA, OmpC, OmpD, and OmpF) in *Salmonella* Typhimurium (STM) pathogenesis. OmpA played a multifaceted role in STM pathogenesis, and a strain deleted for *ompA* (STM *ΔompA*) showed enhanced proneness to phagocytosis and compromised proliferation in macrophages. However, in the epithelial cells, despite being invasion deficient, it was hyper-proliferative. The poor colocalization of STM *ΔompA* with LAMP-1 confirmed impaired stability of SCV membrane around the intracellular bacteria, resulting in its (STM *ΔompA*) release into the cytosol of macrophages where it is assaulted with reactive nitrogen intermediates (RNI). The cytosolic localization of STM *ΔompA* was responsible for the downregulation of SPI-2 encoded virulence factor SpiC, which is required to suppress the activity of iNOS. The reduced recruitment of nitrotyrosine on STM in the macrophage cytosol upon ectopically expressing Listeriolysin O (LLO) explicitly supported the pro-bacterial role of OmpA against the host nitrosative stress. Further, we show that the generation of time-dependent redox burst could be responsible for the enhanced sensitivity of STM *ΔompA* towards nitrosative stress. The absence of OmpA in STM *ΔompA* resulted in the loss of integrity and enhanced porosity of the bacterial outer membrane, which was attributed to the upregulated expression of *ompC*, *ompD,* and *ompF*. We showed the involvement of OmpF in the entry of excess nitrite in STM *ΔompA,* thus increasing the susceptibility of the bacteria towards *in vitro* and *in vivo* nitrosative stress. In conclusion, we illustrated a mechanism of strategic utilization of OmpA compared to other porins by wildtype *Salmonella* for combating the nitrosative stress in macrophages.

## Introduction

The pathogenicity of many Gram-negative bacteria of the family *Enterobacteriaceae* is regulated by porins, commonly known as outer membrane proteins (or Omp). Porins are outer membrane-bound β barrel proteins with 8 to 24 antiparallel β strands connected by loops and are well known for their role in the selective diffusion of ions and solutes across the outer membrane of bacteria [1]. Despite their roles in maintaining outer membrane stability, their pathogenic functions have also been well documented. OmpA, one of the most abundant porins of the bacterial outer membrane, is extensively utilized by *Klebsiella pneumoniae* to prevent the IL-8 dependent pro-inflammatory response in the airway epithelial A549 cells [2]. The deletion of *ompC* and *ompF* from pathogenic *E. coli* not only impaired its invasion in bEnd.3 cells but also reduced its virulence in a mouse model [3]. OprF, an OmpA ortholog in the outer membrane of *Pseudomonas sp.,* has been reported to function as a sensor of quorum signaling and to induce virulence [4]. *E. coli* OmpW has been reported to play a significant role against phagocytosis and complement activation [5, 6].

The outer membrane of *Salmonella* Typhimurium, a member of the *Enterobacteriaceae* family that causes typhoid fever-like symptoms in mice and self-limiting gastroenteritis in humans, is densely populated with many porins such as OmpA, OmpC, OmpD, and OmpF. Unlike OmpA, which has a significant role in the tight attachment of the outer membrane to the underlying peptidoglycan layer with its periplasmic tail [7], the other porins facilitate transportation of charged ions [8]. The connection between the outer membrane porins of *Salmonella* Typhimurium and its pathogenesis remains elusive due to the lack of detailed studies. Earlier, Heijden *et al*. proposed a mechanism of OmpA and OmpC dependent regulation of outer membrane permeability in *Salmonella* in response to H_2_O_2_ and antibiotic stresses [9].

In the current study, we have investigated the individual roles of OmpA, OmpC, OmpD, and OmpF in the pathogenesis of *Salmonella* Typhimurium with a profound focus on OmpA. We found a strong dependence of wild-type *Salmonella* on OmpA for its survival in macrophages and the mouse model. Our study illustrates a mechanism of strategic utilization of OmpA by intracellular *Salmonella* in combatting the nitrosative stress of macrophages by enhancing the outer membrane stability of the bacteria. To the best of our knowledge, this is the first study to demonstrate the impact of outer membrane porins in maintaining the stability of *Salmonella-*containing vacuole (SCV) in macrophages and epithelial cells.

## Results

### OmpA promotes the evasion of phagocytosis and intracellular survival of *Salmonella* in macrophages

The transcriptomic analyses of intracellular *Salmonella* Typhimurium infecting J774-A.1 and HeLa cells have demonstrated almost 2 to 2.5 fold induction in the expression level of *ompA* during early (4h), middle (8h), and late (12h) stages of infection in comparison with other major membrane porins namely *ompC*, *ompD*, and *ompF* [10, 11]. To validate this observation, we isolated total RNA from *Salmonella* Typhimurium strain 14028S grown in nutritionally enriched Luria-Bertani broth, low magnesium minimal acidic F media (pH= 5.4) mimicking the nutrient-deprived acidic environment of *Salmonella* containing vacuole (SCV) [12], and RAW264.7 cells at different time points (3, 6, 9, 12 h post-inoculation/ infection) and investigated the transcript levels of *ompA* (**Figure S1A**), *ompC* (**Figure S1B**), *ompD* (**Figure S1C**), and *ompF* (**Figure S1D**) by real-time PCR. Unlike the steady decrease in the expression levels of *ompC*, *ompD*, and *ompF*, whose expression steadily decreased in wild type bacteria growing in LB broth, F media, and macrophages at all the time points except for *ompF* in macrophages at 9^th^ h post-infection, there was a significant increase in the transcript level of *ompA* during the late phase of infection (9h and 12 h post-infection) in macrophages (**Figure S1A, S1D**). These observations suggest a preference of *ompA* over the other membrane-bound larger porins to enable *Salmonella* to thrive inside macrophages. The observation is also consistent with the microarray data published by Eriksson *et al*. and Hautefort *et al.* [10, 11].

To investigate the importance of *ompA* in the intracellular survival of *Salmonella*, we knocked out *ompA* from the genome of *Salmonella* Typhimurium using a protocol demonstrated earlier by Datsenko and Wanner (Data not shown) [13]. The complementation of *ompA* (1.1 kb) in the knockout bacteria was done using the pQE60 plasmid. A ∼1.42-fold upregulation of *ompA* was found by real-time PCR analysis compared to the wild type *S.* Typhimurium (Data not shown). Unpublished data from our lab suggested that knocking out *ompA* from *Salmonella* did not alter the *in vitro* growth of the bacteria in LB broth. To determine the role of *S.* Typhimurium *ompA* in phagocytosis, we checked phagocytosis of the wild-type and knockout strains in macrophages (**Figure 1A**). We found that the OmpA of STM (WT) helps evade phagocytosis by RAW264.7 and activated U937 cells (**Figure 1A**). The percent phagocytosis of STM *ΔompA* in RAW 264.7 and U937 cells is more than STM (WT).

**Figure 1.**
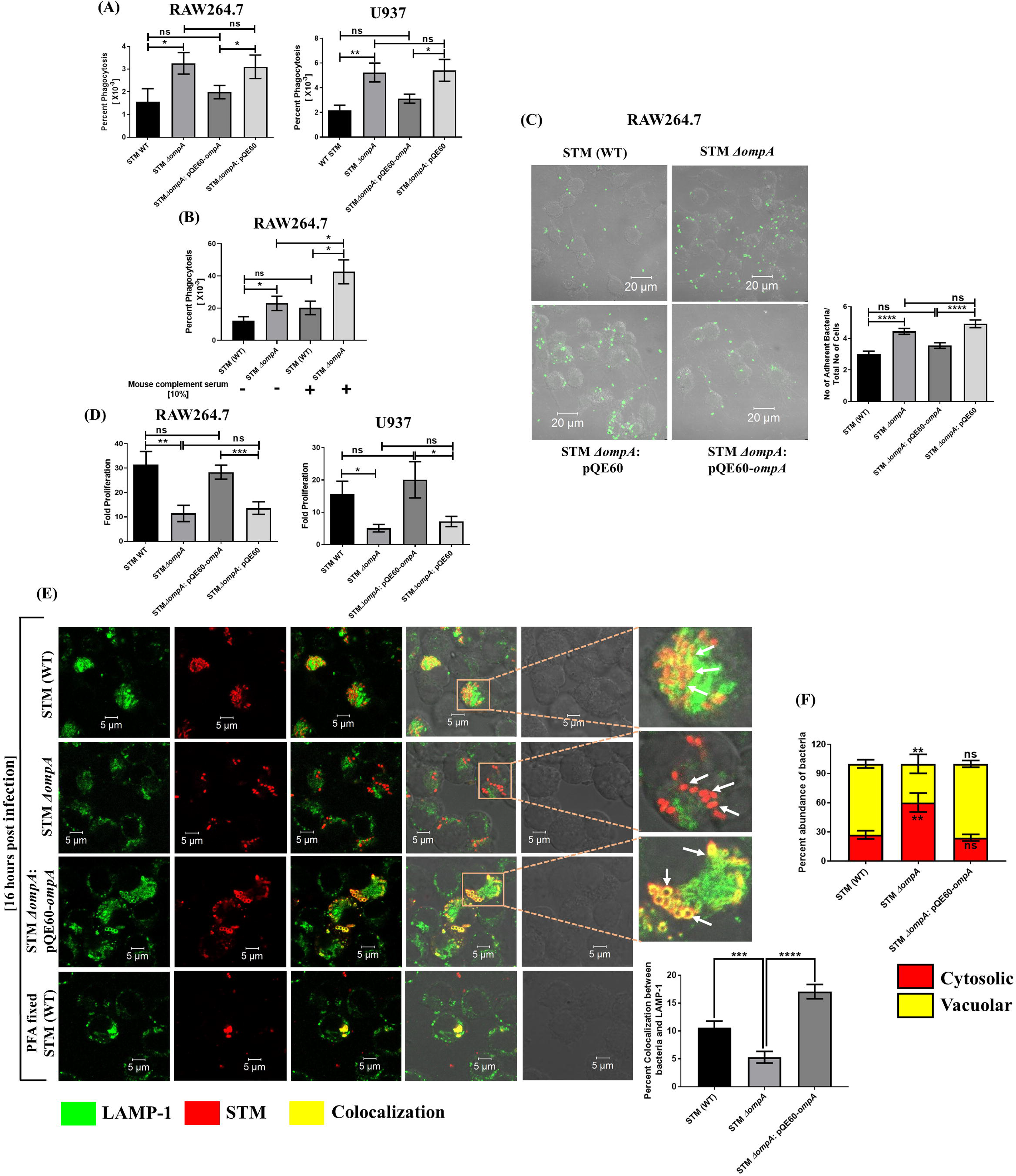
OmpA promotes the evasion of phagocytosis and intracellular survival of *Salmonella* in macrophages. (A) RAW 264.7 cells and PMA activated U937 cells were infected with STM wild type (WT), *ΔompA*, *ΔompA*: pQE60-*ompA*, & *ΔompA*: pQE60 respectively at MOI 10. 2 hours post-infection, the cells were lysed and plated. The CFU at 2 hours was normalized with pre-inoculum to calculate the percent phagocytosis. Data are represented as mean ± SEM (n=3, N=3 for RAW264.7 cells and n=3, N=2 for activated U937 cells). (B) RAW264.7 cells were infected with either 10% mouse complement sera treated or untreated STM (WT) and *ΔompA* respectively at MOI of 50. 2 hours post-infection, the cells were lysed and plated. The CFU at 2 hours was normalized with pre-inoculum to calculate the percent phagocytosis. Data are represented as mean ± SEM (n=3, N=2). (C) RAW 264.7 cells were infected with STM (WT), *ΔompA, ΔompA*: pQE60-*ompA*, & *ΔompA*: pQE60 respectively at MOI 50. Cells were fixed at 15 minutes post-infection with PFA. Externally attached bacteria were probed with anti-*Salmonella* antibody without saponin. 20 microscopic fields were analyzed to calculate the no. of adherent bacteria/ total no. of cells per field. Scale bar = 20μm. Data are represented as mean ± SEM (n=20, N=3). (D) RAW264.7 cells and PMA activated U937 cells were infected with STM- (WT), *ΔompA*, *ΔompA*: pQE60-*ompA*, & *ΔompA*: pQE60 respectively at MOI 10. 16- and 2-hours post-infection, the cells were lysed and plated. The CFU at 16 hours was normalized with the CFU at 2 hours to calculate the fold proliferation of bacteria in macrophages. Data are represented as mean ± SEM (n=3, N=3 for RAW 264.7 cells and n=3, N=2 for activated U937 cells). (E) RAW264.7 cells were infected with STM- (WT): RFP, *ΔompA*: RFP, *ΔompA*: pQE60-*ompA* at MOI of 20. PFA-fixed wild-type bacteria were used for infection at MOI 25. Cells were fixed at 16 hours post-infection & LAMP-1 was labeled with anti-mouse LAMP-1 antibody. To stain the complemented strain and PFA fixed dead bacteria anti-*Salmonella* antibody was used. The quantification of LAMP-1 recruitment on bacteria in RAW 264.7 cells has been represented in a graph. Percent colocalization was determined after analyzing 50 different microscopic stacks from three independent experiments. Scale bar = 5μm. Data are represented as mean ± SEM. (n=50, N=3). (F) Chloroquine resistance assay of RAW 264.7 cells infected with STM- (WT), *ΔompA*, *ΔompA*: pQE60-*ompA* strains, respectively. Data are represented as mean ± SEM (n=3, N=2). ***(P)* *< 0.05, *(P)* **< 0.005, *(P)* ***< 0.0005, *(P)* ****< 0.0001, ns= non-significant, (Student’s *t-*test- unpaired).**

On the contrary, the complemented strain was less prone to phagocytosis by both the macrophages. To mimic the physiological condition, RAW264.7 cells were further infected with STM (WT), and *ΔompA* coated with 10% mouse complement sera (**Figure 1B**). A marked increase in the phagocytosis of complement coated STM *ΔompA* in comparison with complement uncoated STM *ΔompA* and coated STM (WT) confirms the role of *Salmonella* OmpA against complement recruitment and phagocytosis by macrophages (**Figure 1B**). Adhesion of bacteria onto the host cell surface occurs before it enters the host cell, either by phagocytosis or by invasion [14]. Since we found enhanced phagocytosis of STM *ΔompA* in macrophages compared to wild-type bacteria, we decided to evaluate the bacterial attachment on the macrophage surface by *in vitro* adhesion assay (**Figure 1C**). The adhesion of STM *ΔompA* on phagocytic RAW 264.7 cells is more than STM (WT) (**Figure 1C**). The enhanced adhesion of STM *ΔompA* was abrogated upon complementation with *ompA*. We then performed the intracellular survival assay of STM (WT) and STM *ΔompA* strains in RAW264.7, activated U937 cells, respectively (**Figure 1D**). We observed that the intracellular growth of STM *ΔompA* (fold proliferation in RAW264.7 cells- 11.45± 3.349, U937 cells- 5.075± 1.157) is significantly attenuated in RAW 264.7 (by 2.75 folds) and activated U937 cells (by 3.08 folds) compared to the wild type parent (fold proliferation in RAW264.7 cells- 31.5± 5.347, U937 cells- 15.65± 3.981) (**Figure 1D**). When the cell lines were infected with the complemented strain, there was a recovery of the intracellular proliferation of bacteria (**Figure 1D**). Hence, we conclude that *Salmonella* utilizes OmpA as a double-edged sword to protect the bacteria from phagocytosis and then helps it to survive within macrophages.

Once ingested, *Salmonella* starts invading M cells of the Peyer’s patches in the small intestine with the help of the SPI-1 encoded type 3 secretion system. Hence, we checked the role of *Salmonella* OmpA in bacterial invasion of non-phagocytic epithelial cells (**Figure 2A**). Compared to the wild-type bacteria, STM *ΔompA* exhibited significant attenuation in the invasion of the human colorectal adenocarcinoma cell line- Caco-2 and human cervical cancer cell line-HeLa (**Figure 2A**), and the invasiveness was rescued upon complementation (**Figure 2A**). To further validate this observation, we carried out an *in vitro* adhesion assay using HeLa cells as hosts (**Figure 2B**). STM *ΔompA* showed reduced attachment on the surface of HeLa cells compared to the wild-type bacteria (**Figure 2B**). The inefficiency of the knockout strain to attach to the epithelial cell surface was rescued in the complemented strain (**Figure 2B**). This observation is consistent with the result obtained from the invasion assay and showed utilization of OmpA by *Salmonella* as an important adhesion and invasion tool. We further verified the role of OmpA in maintaining the intracellular life of bacteria in epithelial cells (**Figure 2C**). Surprisingly, compared to wild-type bacteria, STM *ΔompA* exhibited hyperproliferation in Caco-2 and HeLa cells (**Figure 2C**). The intracellular proliferation of complement strain was comparable to the wild-type bacteria in both the cell lines. Since we found opposite outcomes in the intracellular survival of the *ompA* knockout strain of *Salmonella* in two different cell types (attenuation in macrophages and hyper-proliferation in the epithelial cells), we decided to monitor the intracellular niche of the bacteria in both cell types. After entering the host cells, *Salmonella* resides inside a modified phagosomal compartment called SCV, which is acidic. The intracellular life and proliferation of *Salmonella* depend upon the stability and integrity of SCV. *Salmonella* recruits a plethora of host proteins to maintain the sustainability of SCV. During the early stage of infection in macrophages and epithelial cells, SCV is characterized by the presence of the markers of early endocytic pathway such as EEA1, Rab5, Rab4, Rab11 and transferrin receptors, etc. These proteins are replaced by late endosome markers such as LAMP-1, Rab7, vATPase, etc., within 15 to 45 min post-infection [15, 16]. Since we found an attenuation in the intracellular proliferation of STM *ΔompA* in macrophages and hyperproliferation in epithelial cells, we decided to check the intracellular niche of the bacteria using LAMP-1 as a marker of SCV in RAW264.7 cells (**Figure 1E**) and Caco-2 cells (**Figure 2D**) 16 h post-infection. It was observed that the percent colocalization of LAMP-1 with STM *ΔompA* is less compared to STM (WT) in RAW264.7 cells (**Figure 1E**). The colocalization with LAMP-1 increased when the macrophage cells were infected with complement strain. These observations corroborated not only the gradual loss of the SCV membrane from its surrounding but also its enhanced cytosolic localization in macrophages (**Figure 1E**). The PFA-fixed dead bacteria lacking the ability to quit vacuole was used as a positive control in this study (**Figure 1E**). The environment within the SCV is acidic (pH= 5.4) in comparison with the cytosol (pH= 7.4) of the macrophages [17], [18]. This acidic environment of the SCV is sensed by the wild-type *Salmonella* Typhimurium with the help of *envZ/ ompR* and *phoP/ phoQ* two-component systems, which activates the expression of SPI-2 genes [17, 19, 20]. The SPI-2 codes for a multiprotein needle-like complex called type III secretion system (T3SS) and several effector proteins (called translocon), which are primarily accumulated on the surface of the matured SCV and secreted into the cytosol of the host cells during the late stage of infection to enhance the severity of the infection (**Figure S2A**). The assembly of the SPI-2 encoded effector proteins, namely SseB, SseC, SseD, on the surface of the SCV facilitates the formation of a functionally active T3SS needle complex, which in turn helps in the systemic colonization of the bacteria (**Figure S2A**) [21, 22]. Since the cytosolic population of STM *ΔompA* lacks the vacuolar membrane, we anticipated an interruption in the accumulation of these virulent proteins on the bacterial surface and their secretion into the host cytosol. In line with our previous observations, we found a marked reduction in the accumulation and secretion (the area of the infected macrophage demarcated with a dotted line for the wild-type *Salmonella*) of SseC (**Figure S2B and S2C**) and SseD (**Figure S2B and S2D**) on or from the surface of STM *ΔompA* into the host cytosol. In continuation of this observation, we found a dampened expression of *sseC* (**Figure S2E**) and *sseD* (**Figure S2F**) genes in intracellularly growing STM *ΔompA* compared to the wild-type bacteria. We further wanted to test the impact of the cytosolic localization of STM *ΔompA* on the expression profile of several other important SPI-2 virulent genes such as *ssaV* (**Figure S2G**) and *sifA* (**Figure S2H**). The attenuated expression of *ssaV* and *sifA* suggested that unlike the acidic pH of SCV, the cytosolic pH of the macrophage does not favor the expression of SPI-2 virulent genes, which could be a reason behind the compromised growth of *ompA* deficient bacteria in macrophage. To verify whether this is a cell type-specific phenomenon or not, we further investigated the intracellular niche of wild type, knockout, and complement strains of *Salmonella* in epithelial Caco-2 cells. The quantitative percent colocalization between LAMP-1 and all three bacterial strains (**Figure 2D**) demonstrated that STM *ΔompA* comes into the cytoplasm of Caco-2 cells after being released from the SCV during the late phase of infection. In contrast, wild-type and the complemented strains abstained themselves from doing so (**Figure 2D**). These findings were further supported by chloroquine resistance assay of STM-(WT), *ΔompA*, *ΔompA*: pQE60-*ΔompA* in RAW264.7 (**Figure 1F**) and Caco-2 cells (**Figure 2E**). After being protonated inside SCV because of the acidic pH, chloroquine cannot quit the vacuole and kills the vacuolar population of *Salmonella*, allowing the growth of the cytosolic population. During the late phase of infection (16 h post-infection) in RAW264.7 cells (**Figure 1F**) and Caco-2 cells (**Figure 2E**), the cytosolic abundance of STM *ΔompA* was found more comparable to the wild type and complement strain. Taken together, these results authenticate that, irrespective of the cell type, the intravacuolar life of the intracellular *Salmonella* strongly depends upon OmpA.

**Figure 2.**
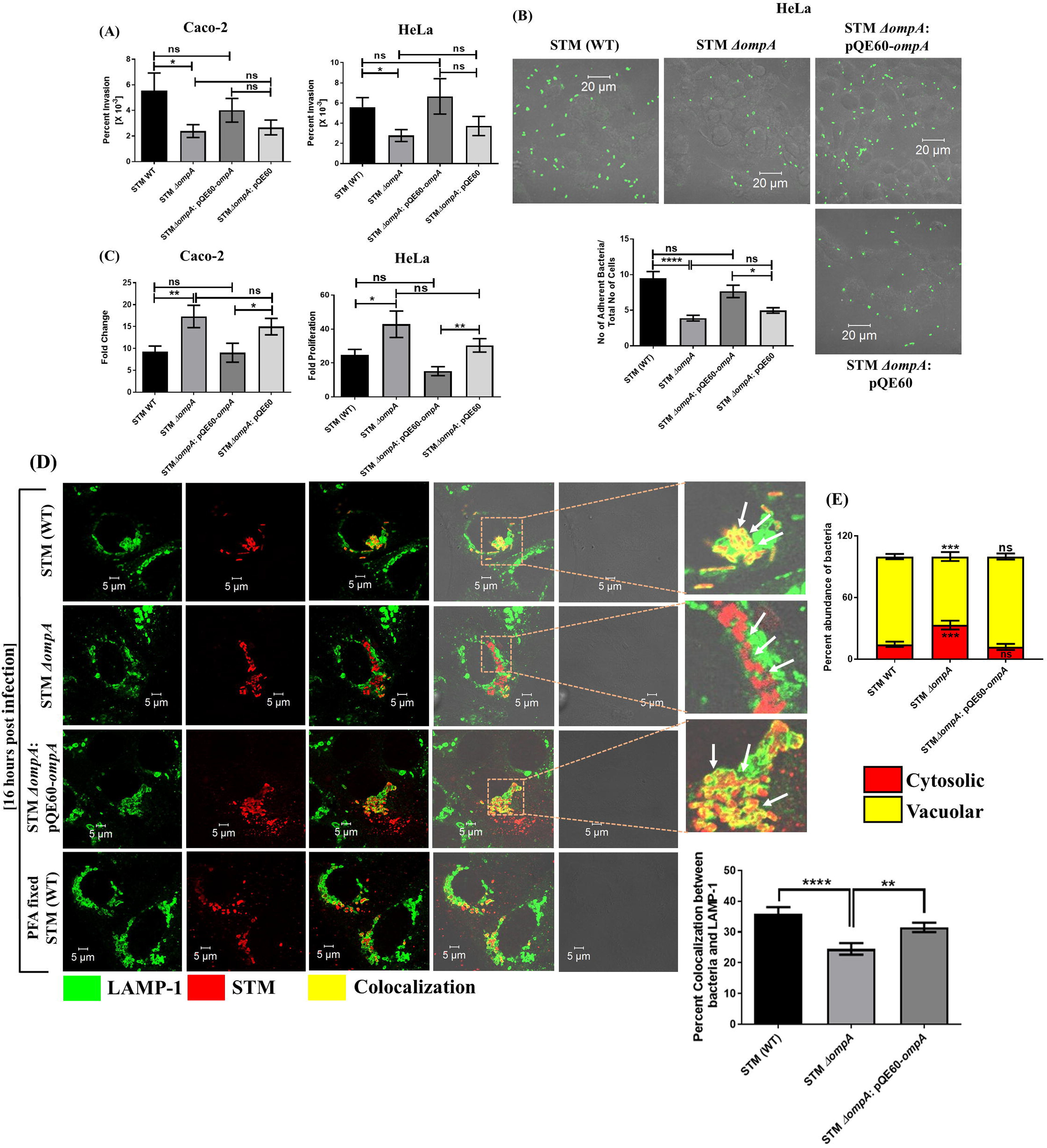
OmpA dependent invasion and intracellular proliferation of *Salmonella* inside the epithelial cell. (A) Caco-2 cells and HeLa cells were infected with log phase culture of STM (WT), *ΔompA*, *ΔompA*: pQE60-*ompA*, & *ΔompA*: pQE60 respectively at MOI 10. 2 hours post-infection, the cells were lysed and plated. The CFU at 2 hours was normalized with pre-inoculum to calculate the percent invasion in epithelial cells. Data are represented as mean ± SEM (n=3, N=3). (B) HeLa cells were infected with the log phase culture of STM (WT), *ΔompA*, *ΔompA*: pQE60- *ompA*, & *ΔompA*: pQE60, respectively at MOI of 50. Cells were fixed at 25 minutes post- infection with PFA. Externally attached bacteria were probed with anti-*Salmonella* antibody without saponin. 20 microscopic fields were analyzed to calculate the no. of adherent bacteria/ total no. of cells per field. Data are represented as mean ± SEM (n=20, N=3). Scale bar = 20μm. (C) Caco-2 and HeLa cells were infected with log phase culture of STM- (WT), *ΔompA*, *ΔompA*: pQE60-*ompA*, & *ΔompA*: pQE60 respectively at MOI 10. 16- and 2-hours post-infection, the cells were lysed and plated. The CFU at 16 hours was normalized with the CFU at 2 hours to calculate the fold proliferation of bacteria in epithelial cells. Data are represented as mean ± SEM (n=3, N=3). (D) Caco-2 cells were infected with STM (WT): RFP, *ΔompA*: RFP, and *ΔompA*: pQE60-*ompA* at MOI 20. PFA-fixed wild-type bacteria were used for infection at MOI 25. Cells were fixed at 16 hours post-infection & LAMP-1 was labeled with anti-human LAMP-1 antibody. The complemented strain and PFA fixed dead bacteria were tracked with an anti-*Salmonella* antibody. The quantification of LAMP-1 recruitment on bacteria in Caco-2 cells has been represented in the form of a graph. Percent colocalization was determined after analyzing 50 different microscopic stacks from three independent experiments. Scale bar = 5μm. Data are represented as mean ± SEM (n=50, N=3). (E) Chloroquine resistance assay of Caco-2 cells infected with STM- (WT), *ΔompA*, *ΔompA*: pQE60-*ompA* strains. Data are represented as mean ± SEM (n=3, N=2). ***(P)* *< 0.05, *(P)* **< 0.005, *(P)* ***< 0.0005, *(P)* ****< 0.0001, ns= non-significant, (Student’s *t-*test- unpaired).**

### OmpA dependent protection of *Salmonella* against nitrosative stress inside RAW264.7 cells

The intracellular population of *Salmonella* is heavily encountered by NADPH phagocytic oxidase-dependent oxidative burst and iNOS dependent nitrosative burst during early and late stages of infection in macrophages, respectively [23, 24]. The SCV membrane protects the vacuolar niche of wild-type *Salmonella* from the potential threats present in the cytoplasm of macrophages in the form of Reactive Oxygen Species (ROS), Reactive Nitrogen Intermediates (RNI), antimicrobial peptides (AMPs), etc. [25, 26]. Our study confirmed the proneness of STM *ΔompA* release from SCV into the cytoplasm of macrophages and epithelial cells during the late stage of infection. Hence, we assumed the increasing possibility of the mutant bacteria being targeted with ROS and RNI in the cytoplasm of the macrophages, which could be a probable reason for attenuation of intracellular proliferation. Our hypothesis was verified by measuring nitrosative (**Figure 3A and 3B**) and oxidative (**Figure S3A – S3C**) stress response of macrophages infected with wild type, knockout, and complemented strains. We quantified the extracellular (**Figure 3A**) [NO] response produced by infected macrophages by Griess assay. It was found that during the late stage of infection (16 h post-infection), the accumulation of nitrite in the culture supernatant of RAW264.7 cells infected with STM *ΔompA* was significantly higher compared to the wild type. This heightened [NO] response was revoked when the RAW264.7 cells were infected with complement strain (**Figure 3A**). This result was further validated by quantifying intracellular [NO] response using DAF2-DA (**Figure 3B**). Only 2.22 % of wild-type bacteria-infected macrophages produced [NO] (Figure 3.6B), which increased to 6.17 % when the cells were infected with STM *ΔompA* (**Figure 3B**). The percentage of RAW264.7 cells producing [NO] after being infected with complement strains 3.94 % was comparable to the wild-type STM infected cells (Figure 3B). During the late stage of infection (16 h post-infection) in macrophages, the intracellular ROS and extracellular H_2_O_2_ levels were estimated using H_2_DCFDA (**Figure S3A and S3B**) and phenol red assay (**Figure S3C**), respectively.

**Figure 3.**
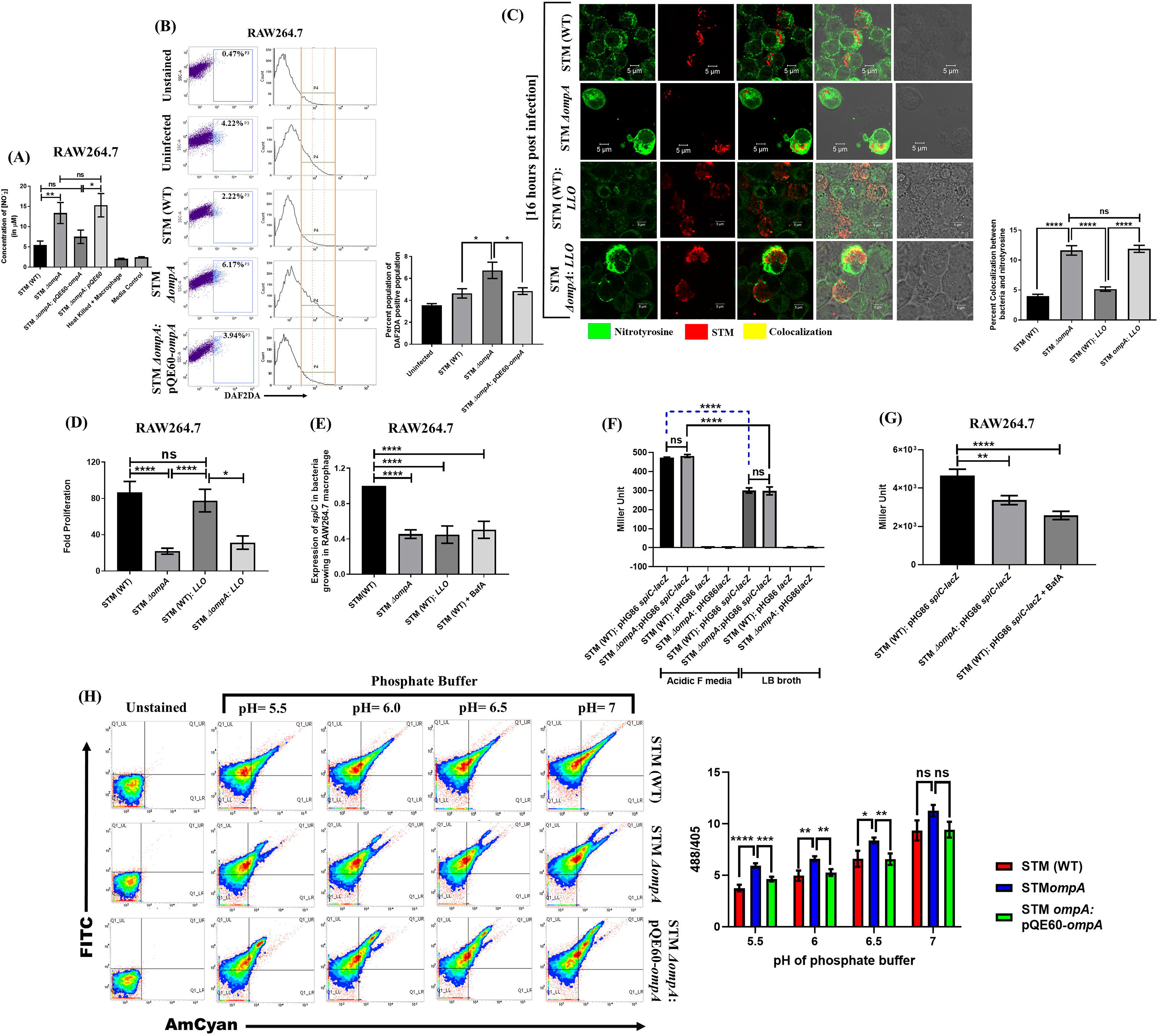
OmpA dependent protection of *Salmonella* against nitrosative stress inside RAW264.7 cells. (A) Estimation of extracellular nitrite from the culture supernatant of RAW264.7 cells infected with STM (WT), *ΔompA*, *ΔompA*: pQE60-*ompA*, *ΔompA*: pQE60, & heat-killed bacteria respectively at MOI 10. 16 hours post-infection, the culture supernatants were collected and subjected to Griess assay. Data are represented as mean ± SEM (n=3, N=5). (B) Estimation of intracellular nitric oxide level in RAW 264.7 cells infected with STM (WT), *ΔompA*, and *ΔompA*: pQE60-*ompA* at MOI 10, 16 hours post-infection using DAF-2 DA [5 µM] by flow cytometry. Unstained and uninfected RAW264.7 cells have also been used as controls. Both dot plots (SSC-A vs. DAF-2 DA) and histograms (Count vs. DAF-2 DA) have been represented. The percent population of DAF-2 DA positive RAW264.7 cells has been represented in the form of a graph. Data are represented as mean ± SEM (n≥3, N=4). (C) Immunofluorescence image of RAW264.7 cells infected with STM (WT): RFP, *ΔompA*: RFP, (WT): *LLO*, and *ΔompA*: *LLO* at MOI 20. 16 hours post-infection, the cells were fixed with PFA and labeled with anti-*Salmonella* antibody and anti-mouse nitrotyrosine antibody followed by visualization under confocal microscopy. Percent colocalization of STM (WT), *ΔompA*, (WT): *LLO,* and *ΔompA*: *LLO* with nitrotyrosine was determined by analyzing 50 different microscopic stacks from three independent experiments. Data are represented as mean ± SEM (n=50, N=3). Scale bar = 5μm. (D) RAW264.7 cells were infected with STM (WT), *ΔompA*, (WT): *LLO,* and *ΔompA*: *LLO* at MOI 10. 16- and 2-hours post-infection, the cells were lysed and plated. The CFU at 16 hours was normalized with the CFU at 2 hours to calculate the fold proliferation of bacteria. Data are represented as mean ± SEM (n≥3, N=2). (E) The transcript-level expression profile of *spiC* from RAW264.7 cells infected with STM (WT), *ΔompA* & (WT): *LLO* at MOI 50. STM (WT) infected RAW264.7 cells treated with bafilomycin A (50 nM) were used as a control. 12 hours post-infection, the cells were lysed, and total RNA was extracted. After the synthesis of cDNA, the expression of *spiC* was measured by RT PCR. Data are represented as mean ± SEM (n=3, N=3). (F) The measurement of the activity of *spiC* promoter in stationary phase culture of STM (WT) and *ΔompA* growing overnight in acidic F media and neutral LB media. Data are represented as mean ± SD (n=6). (G) The measurement of the activity of *spiC* promoter in STM (WT) and *ΔompA* proliferating intracellularly in RAW264.7 cells (MOI= 50) 12 hours post-infection. Macrophages infected with STM (WT) and further treated with bafilomycin A (50 nM) have been used as a negative control. Data are represented as mean ± SEM (n=5, N=2). (H) Estimation of intracellular acidification of STM (WT), *ΔompA*, and *ΔompA*: pQE60-*ompA* in phosphate buffer of pH 5.5, 6, 6.5, and 7, respectively using 20 µM of ratiometric pH indicator BCECF-AM by flow cytometry. The ratio of median fluorescence intensity of BCECF-AM labeled on STM- (WT), *ΔompA*, and *ΔompA*: pQE60-*ompA* at 488 and 405 nm, respectively. Data are represented as mean ± SEM (n=4, N=3). ***(P)* *< 0.05, *(P)* **< 0.005, *(P)* ***< 0.0005, *(P)* ****< 0.0001, ns= non-significant, (Student’s *t* test- unpaired).**

We did not find any considerable change in the population of cells producing ROS, infected with wild-type (3.06%; **Figure S3A**) (2.157 ± 0.1611) % (**Figure S3B**), knockout (4.09%; **Figure S3A**) (2.192 ± 0.2955) % (**Figure S3B**), and the complement strains (3.81%; **Figure S3A**) (2.61 ± 0.2244) % (**Figure S3B**) of *Salmonella* (**Figure S3A and S3B**). We further checked the accumulation of H2O2 in the culture supernatants of infected RAW264.7 cells to validate this observation. But there was hardly any difference in H_2_O_2_ production (**Figure S3C**). The [NO] produced from the cellular pool of L-arginine by the catalytic activity of inducible nitric oxide synthase (iNOS) is further oxidized into [NO]-adducts (NONOates, peroxynitrite, nitrite, etc.) with the help of the superoxide ions which have higher oxidation potential and bactericidal activity [27]. The damage caused by peroxynitrite (ONOO^-^) can be monitored by checking the recruitment of nitrotyrosine on the surface of intracellular bacteria using specific anti-nitrotyrosine antibodies by confocal microscopy (**Figure 3C**). During the late stage of infection in RAW264.7 cells, the STM *ΔompA* strain showed higher surface recruitment of nitrotyrosine residues and greater colocalization than the wild-type bacteria (**Figure 3C**), indicating massive damage caused by the bactericidal function of peroxynitrite. *Listeria monocytogenes* produce listeriolysin O (LLO), a pore-forming toxin to degrade the vacuolar membrane to escape lysosomal fusion [28]. Despite having intact OmpA on its surface, wild-type *Salmonella* residing in the cytoplasm due to the ectopic expression of listeriolysin O (LLO) showed poor colocalization with nitrotyrosine (**Figure 3C**). These results suggest the ability of OmpA to protect the cytosolic population of wild-type *Salmonella* against the RNI of murine macrophages. The greater recruitment of nitrotyrosine on STM *ΔompA*: *LLO,* which is comparable to STM *ΔompA,* suggests that LLO does not play any role in modulating the activity of iNOS. Consistent with this, the intracellular survival of STM (WT): *LLO* in macrophages was better than STM *ΔompA* and comparable to that of the wild-type bacteria (**Figure 3D**). Taken together, we show that OmpA protects intracellular wild-type *Salmonella* Typhimurium against the nitrosative stress of macrophages. While proliferating in macrophages, it utilizes SPI-2 encoded virulent factor SpiC to downregulate inducible nitric oxide synthase activity in SOCS-3 dependent manner [29, 30]. To verify whether the OmpA dependent downregulation of the action of iNOS happens in SpiC dependent manner, the transcript level of *spiC* was measured from wild-type and the mutant bacteria growing in macrophages (**Figure 3E**). Unlike the wild-type *Salmonella*, STM *ΔompA* was unable to produce *spiC* within macrophages. The reduced expression of *spiC* by STM (WT): *LLO* and bafilomycin A treated cells infected with wild-type bacteria not only shows the requirement of the acidic pH of SCV for the expression of *spiC* but also indirectly authenticates the cytosolic localization of STM *ΔompA*. To validate this observation, the promoter activity of *spiC* was measured in STM (WT) and *ΔompA* growing in acidic F media (**Figure 3F**), LB media (**Figure 3F**), and macrophages (**Figure 3G**) by beta-galactosidase assay. No significant change was observed in the activity of the *spiC* promoter between the wild-type and mutant bacteria growing in acidic F media and LB broth for 12 hours (**Figure 3F**). However, the enhanced activity of the *spiC* promoter in STM (WT) and *ΔompA* growing in F media (pH= 5.4) compared to the bacterial culture obtained from LB broth (which is less acidified compared to the F media) suggests that the expression of the *spiC* gene is regulated by the acidification of the environment around the bacteria. Inside the macrophages, where there exists a notable difference between the localization of wild-type and *ompA* deficient bacteria, a significant drop in the activity of the *spiC* promoter was observed in STM *ΔompA* (**Figure 3G**). The localization of STM *ΔompA* in the cytosol of macrophages where the pH is relatively higher than SCV can be held accountable for the abrogated expression of *spiC*. To determine the degree of acidification of the cytosol of STM (WT) and *ΔompA* upon alteration in the pH of the surrounding media, we have used a pH-sensitive dye BCECF-AM. We observed a higher 488 nm/ 405 nm ratio of STM *ΔompA* labeled with 20 µM concentration of BCECF-AM resuspended in phosphate buffer of acidic pH (range-5.5, 6, and 6.5) in comparison with STM (WT) and the complemented strain (*ΔompA*: pQE60-*ompA*) (**Figure 3H**). This result suggests reduced acidification of the cytosol of STM *ΔompA* compared to STM (WT) and STM *ΔompA*: pQE60-*ompA* even when they are present in the same (*in vitro*) acidic environment (**Figure 3H**). Surprisingly when all these strains were incubated separately in a phosphate buffer of pH= 7, we found a comparable 488 nm/ 405 nm ratio of BCECF-AM, which unveils an uncharacterized novel but controversial role of OmpA in the acidification of the cytosol of *S.* Typhimurium in response to extracellular acidic stress.

### Inhibition of the activity of iNOS improves the survival of STM *ΔompA* in the *in vitro* and *in vivo* infection models

To investigate further the pro-bacterial role of *Salmonella* OmpA against the nitrosative stress of macrophages, we treated the cells with 1400W dihydrochloride (10 µM), an irreversible inhibitor of inducible nitric oxide synthase (iNOS), the key enzyme in the production of [NO] (**Figure 4A, 4C, and 4D**), or mouse IFN-ɣ that upregulates the expression of iNOS (at 100 U/ mL concentration) (**Figure 4B, 4C, and 4E**). Inhibition of iNOS using 1400W completely restored the intracellular proliferation of STM *ΔompA* (43.61 ± 6.837) compared to the untreated reference (19.32 ± 3.913) (**Figure 4A**). Consistent with this finding, STM *ΔompA* showed poor colocalization with nitrotyrosine under 1400W treatment (**Figure 4C and 4D**), which can be attributed to the poor biogenesis of RNI due to the inhibition of iNOS. Augmenting the iNOS activity of macrophages using mouse IFN-ɣ hindered the intracellular proliferation of STM *ΔompA* (12.52 ± 1.334) compared to the IFN-ɣ untreated cells (19.64 ± 2.11) (**Figure 4B**). As demonstrated by the confocal image, the enhanced biogenesis and colocalization of nitrotyrosine with STM ΔompA under IFN-ɣ treatment can be considered for its higher attenuation in intracellular proliferation (**Figure 4C and 4E**). The reduced CFU burden of *ompA* deficient *Salmonella* in the liver, spleen, and MLN of C57BL/6 mice (compared to the wild-type bacteria) strongly supports the role of OmpA in bacterial pathogenesis (**Figure 4G**). In our *ex vivo* studies, we established the role of OmpA in protecting the wild-type *Salmonella* against the nitrosative stress of murine macrophages. Further, we used *iNOS^-/-^* C57BL/6 mice and treated the wild type C57BL/6 mice with a pharmacological inhibitor of iNOS called aminoguanidine hydrochloride at a dose of 10 mg/ kg of body weight from 0^th^ day (On the day of infecting the mice) to 5^th^day (the day of sacrificing and dissecting the mice) (**Figure 4G**) [31]. We found a comparable CFU burden of wild type and *ompA* deficient bacteria in the liver, spleen, and MLN of *iNOS^-/-^* and wild type C57BL/6 mice administered with the inhibitor. The higher bacterial burden of STM *ΔompA* from the *iNOS^-/-^* and aminoguanidine-treated mice compared to PBS treated mice (**Figure 4G**) concomitantly authenticates our results obtained from *ex vivo* experiments. Likewise, the use of the *gp91phox***^-/-^** mice unable to produce ROS upon bacterial infection showed comparable CFU burden for both the wild type and *ΔompA* STM in the liver, spleen, and MLN. A significantly higher bacterial load of *ompA* deficient STM in the liver of *gp91phox***^-/-^** vs. wild type C57BL/6 mice indicates the inability of *gp91phox***^-/-^** mice to generate peroxynitrite due to dampened production of superoxide ions (ROS) (Figure 3.8G). However, a comparable burden of STM *ΔompA* and the wild-type strain in the spleen and MLN of *gp91phox***^-/-^** mice and the wild-type C57BL/6 mice suggests that ROS alone cannot clear the *in vivo* infection by STM *ΔompA*. Earlier, we found that the infection of RAW264.7 cells with STM *ΔompA* does not induce the level of intracellular ROS. To show the role of *ompA* in the *in vivo* infection of *Salmonella* Typhimurium, we challenged 4-6 weeks old adult BALB/c and C57BL/6 mice (**Figure 4H**) with a lethal dose (10^8^ CFU of bacteria/ animal) of wild type and knockout strains by oral gavaging. Almost 80% of BALB/c mice infected with STM *ΔompA* survived compared to the group of mice infected with wild-type bacteria (**Figure 4H**). On the other side, the C57BL/6 mice infected with STM *ΔompA* showed better survival and retarded death than those infected with the wild-type strain, suggesting a critical role of ompA in the infection caused by *Salmonella*.

**Figure 4.**
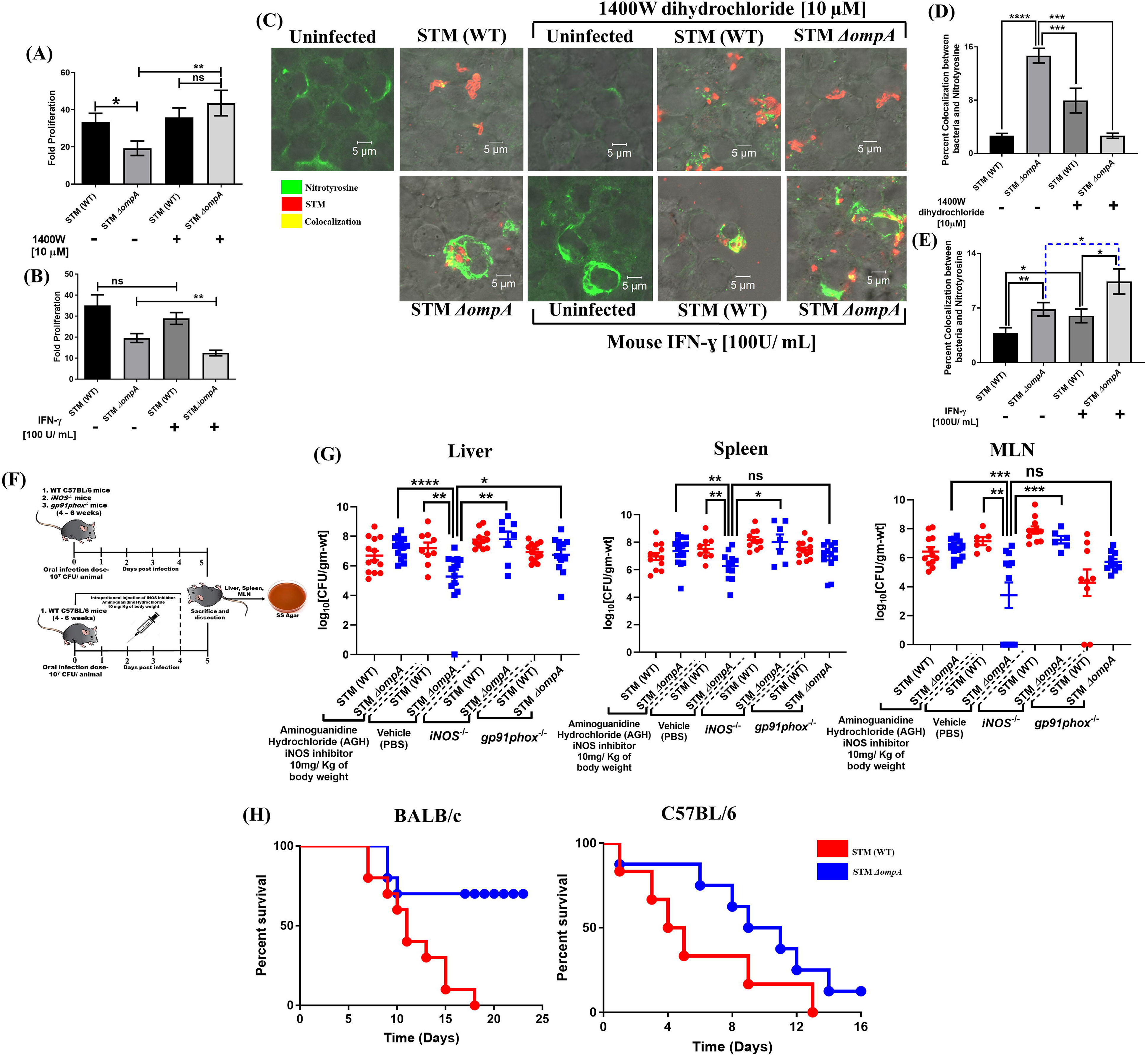
Alteration in the activity of iNOS by using a specific inhibitor or activator changes the fate of STM *ΔompA* in *in vitro* and *in vivo* infection models. Intracellular survival of STM (WT) and *ΔompA* (MOI= 10) in RAW264.7 cells (16 hours post-infection) in presence and absence of iNOS (A) inhibitor-1400W dihydrochloride [10 µM] and (B) activator-mouse IFN-ɣ [100U/ mL]. Fold proliferation of bacteria was calculated by normalizing the CFU at 16 hours to CFU at 2 hours. Data are represented as mean ± SEM (n=3, N=3). (C) Immunofluorescence image of RAW264.7 cells infected with STM- (WT) and *ΔompA* (MOI=20) in presence and absence of iNOS inhibitor- 1400W dihydrochloride [10 µM] and activator-mouse IFN-ɣ [100U/ mL]. 16 hours post-infection, the cells were fixed with PFA and probed with anti-*Salmonella* antibody and anti-mouse nitrotyrosine antibody. The quantification of the recruitment of nitrotyrosine on STM (WT) and *ΔompA* (MOI-20) in RAW264.7 cells (16 hours post-infection) in presence and absence of iNOS (D) inhibitor- 1400W dihydrochloride [10 µM] and (E) activator- mouse IFN-ɣ [100U/ mL]. Percent colocalization of bacteria with nitrotyrosine was determined after analyzing more than 50 different microscopic stacks from two independent experiments. Data are represented as mean ± SEM (n≥ 50, N=2). (F) The schematic representation of the strategy of animal experiments. (G) Four cohorts of 4 to 6 weeks old five C57BL/6 mice were orally gavaged with STM- (WT) and *ΔompA* at a sublethal dose of 10^7^ CFU/ animal. Two of these four cohorts were intraperitoneally injected with iNOS inhibitor aminoguanidine hydrochloride (10mg/ kg of body weight) regularly for 5 days post-infection. Two cohorts of five *iNOS^-/-^* and *gp91phox^-/-^* mice were orally gavaged with STM- (WT) and *ΔompA* at a sublethal dose of 10^7^ CFU/ animal separately. On the 5^th^-day post-infection, the mice were sacrificed, followed by isolation, homogenization of liver, spleen, and MLN. The organ lysates were plated on *Salmonella Shigella* agar. The colonies were counted 16 hours post-plating. The log_10_[CFU/ gm-wt.] for each CFU has been plotted. Data are represented as mean ± SEM (n=5, N=3). (H) Two cohorts- each consisting of 4 to 6 weeks old 20- BALB/c and C57BL/6 mice infected with a lethal dose (10^8^ CFU/ animal) of STM- (WT) and *ΔompA*. After they were infected by oral gavaging, the survival of the mice was monitored till the death of all the wild-type infected mice (n=10). *(P)* *< 0.05, *(P)* **< 0.005, *(P)* ***< 0.0005, *(P)* ****< 0.0001, ns= non-significant, (Student’s *t* test- unpaired, Mann-Whitney *U* test- for animal survival assay).

### OmpA dependent regulation of outer membrane permeability in *Salmonella* controls cytoplasmic redox homeostasis in response to *in vitro* nitrosative stress

In a mildly acidic environment (pH= 5 – 5.5), NaNO_2_ dissociates to form nitrous acid (HNO_2_), which undergoes dismutation reaction upon oxidation and eventually generates a wide range of reactive nitrogen intermediates (RNI) such as nitrogen dioxide (NO_2_), dinitrogen trioxide (N_2_O_3_), nitric oxide (NO), etc. [32]. After entering the bacterial cells, RNIs cause irreparable deleterious effects targeting multiple subcellular components such as nucleic acids (cause deamination of nucleotides), proteins (disruption of Fe-S clusters, heme group; oxidation of thiols and tyrosine residues), lipids (cause peroxidation), etc., and finally kill the pathogen [33]. To investigate the role of *Salmonella* OmpA against *in vitro* nitrosative stress, we decided to check *in vitro* sensitivity of the wild type and *ompA* knockout strains in the presence of a varying concentration (0-5 mM) of H_2_O_2_ (**Figure S3D**), NaNO_2_ (**Figure S3E**), and a combination of H_2_O_2_ and NaNO_2_ for 12 h (**Figure S3F**) by CFU and resazurin test. Compared with wild-type bacteria, STM *ΔompA* did not show any significant difference in viability when exposed to peroxide (**Figure S3D**). The knockout strain displayed a substantial reduction in viability compared to the wild-type bacteria at 800 µM concentration when incubated with NaNO_2_ alone (**Figure S3E**). The sensitivity of the knockout strain towards acidified nitrite further increased when H_2_O_2_ was added to NaNO_2_ (where the growth inhibition started at 600 µM concentration) (**Figure S3F**). To ensure OmpA dependent protection of wild-type *Salmonella* against the damage caused by *in vitro* nitrosative stress, we performed a death kinetics experiment with wild-type and knockout bacteria in the presence of 800 µM acidified nitrite (**Figure 5A**). Consistent with our previous observations, the knockout strain showed a notable hindered growth compared to the wild type at 12 h post-inoculation. Considering the impeded growth of the mutant strain in the presence of in vitro nitrosative stress, we hypothesized enhanced entry of nitrite into the *ΔompA* strain compared to the wild type. To verify this, we performed nitrite uptake assay with the wild type, knockout, complement, and empty vector strain in MOPS-NaOH buffer with 200 µM initial nitrite concentration. We noticed a greater time-dependent nitrite uptake by the *ΔompA* and empty vector strain than wild-type and complemented strains (**Figure 5B**).

**Figure 5.**
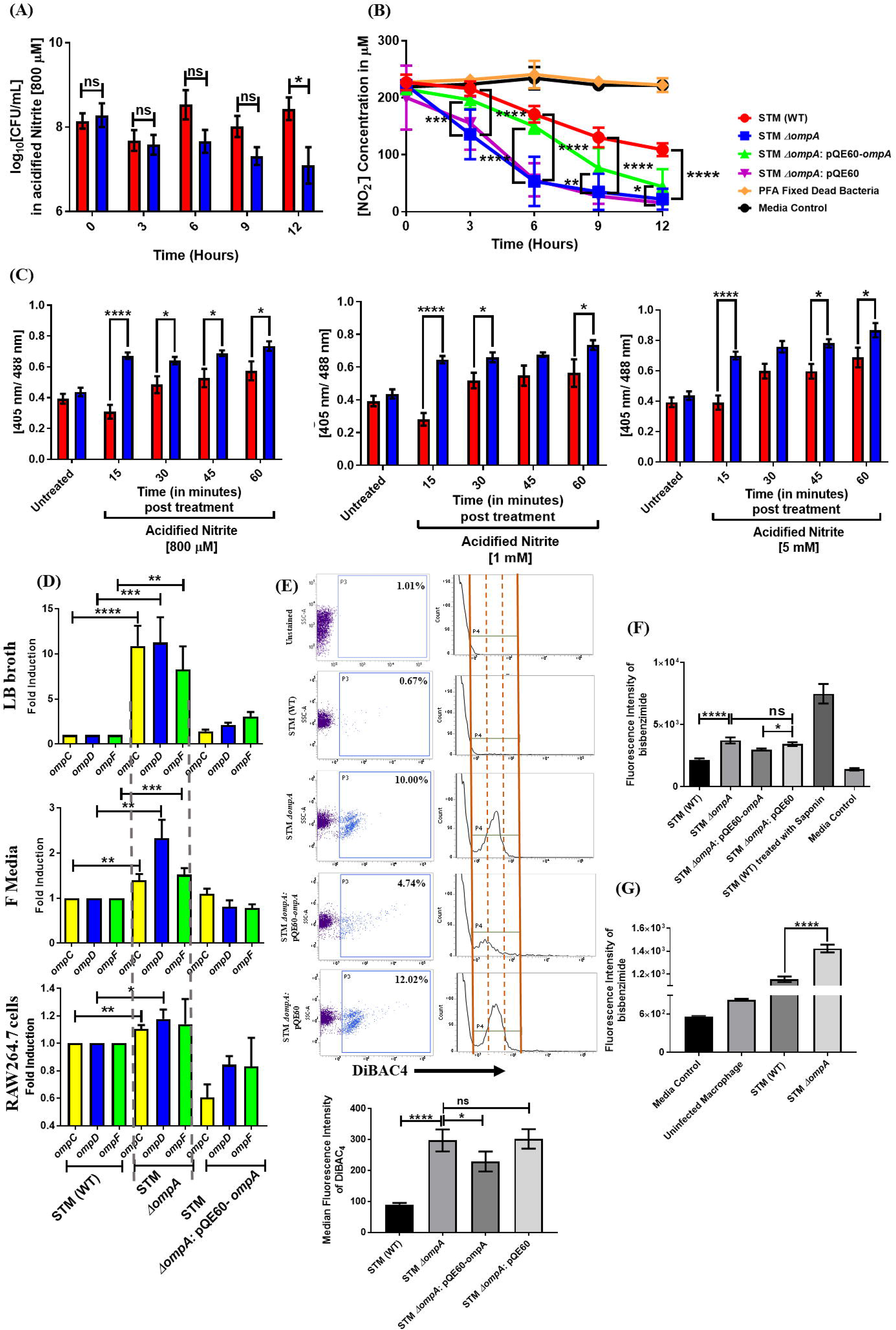
OmpA dependent regulation of outer membrane permeability in *Salmonella* controls cytoplasmic redox homeostasis in response to *in vitro* nitrosative stress. (A) Time-dependent *in vitro* death kinetics of STM (WT) and *ΔompA* in the presence of acidified nitrite (Nitrite concentration 800 µM in PBS of pH 5.4). 10^8^ CFU of overnight grown stationary phase culture of both the strains were inoculated in acidified nitrite. The survival of both the strains was determined by plating the supernatant with appropriate dilution in SS agar plates at 0, 3, 6, 9, 12 hours post-inoculation. Data are represented as mean ± SEM (N=5). (B) *In vitro* nitrite uptake assay of STM (WT), *ΔompA*, *ΔompA*: pQE60-*ompA*, *ΔompA*: pQE60, & PFA fixed dead bacteria. 10^8^ CFU of overnight grown stationary phase culture of all the strains were inoculated in MOPS- NaOH buffer (pH= 8.5) with an initial nitrite concentration of 200 µM. The remaining nitrite concentration of the media was determined by Griess assay at 0, 3, 6, 9, 12 hours post-inoculation. Data are represented as mean ± SEM (n=3, N=4). (C) Measurement of redox homeostasis of STM (WT) and *ΔompA* harboring pQE60-Grx1-roGFP2 in response to varying concentrations of acidified nitrite in a time-dependent manner. 4.5X 10^7^ CFU of overnight grown stationary phase culture of STM (WT) and *ΔompA* expressing pQE60- Grx1-roGFP2 were subjected to the treatment of acidified nitrite (concentration of 800µM, 1mM, 5mM) for 15, 30, 45, and 60 minutes. Median fluorescence intensities of Grx1-roGFP2 at 405nm and 488nm for the FITC positive population were used to obtain the 405/ 488 ratio in 800 µM, 1mM, 5 mM concentration of acidified nitrite, respectively. Data are represented as mean ± SEM (n=3, N=3). (D) The transcript level expression profiling of larger porin genes, namely- *OmpC*, *OmpD*, *OmpF* by real-time PCR in STM (WT), *ΔompA*, & *ΔompA*: pQE60- *ompA* in nutritionally enriched LB media, low magnesium acidic F media (pH= 5.4) mimicking the internal environment of SCV, and RAW264.7 murine macrophage cells (MOI 50) [12 hours post inoculation]. Data are represented as mean ± SEM (n=3, N=2). (E) Measurement of membrane porosity of STM- (WT), *ΔompA*, *ΔompA*: pQE60-*ompA*, & *ΔompA*: pQE60 in acidic F media (12 hours post-inoculation) using a slightly negatively charged dye named DiBAC_4_ (final concentration- 1 µg/ mL) by flow cytometry. Unstained bacterial cells were used as control. Both dot plots (SSC-A vs. DiBAC_4_) and histograms (Count vs. DiBAC_4_) have been represented. The median fluorescence intensity of DiBAC_4_ has been represented here. Data are represented as mean ± SEM (n=3, N=2). (F) Measurement of outer membrane porosity of STM- (WT), *ΔompA*, *ΔompA*: pQE60-*ompA* & *ΔompA*: pQE60 in acidic F media [12 hours post inoculation] using a nuclear binding fluorescent dye bisbenzimide [excitation- 346 nm and emission- 460 nm] (final concentration- 1 µg/ mL). (WT) treated with 0.1% saponin for 15 minutes has been used as a positive control. Data are represented as mean ± SEM (n=8, N=3). (G) Measurement of outer membrane porosity of STM (WT) and *ΔompA* isolated from infected RAW264.7 cells 12 hours post-infection by bisbenzimide (Sigma) (final concentration- 1 µg/ mL). Data are represented as mean ± SEM (n=6, N=3). *(P)* *< 0.05, *(P)* **< 0.005, *(P)* ***< 0.0005, *(P)* ****< 0.0001, ns= non-significant, (2way ANOVA, Student’s *t-*test- unpaired).

To verify OmpA dependent redox homeostasis of *Salmonella* in response to *in vitro* nitrosative stress, we exposed both the wild type and knockout strains harboring pQE60-Grx1-roGFP2 plasmid to 800 µM, 1 mM, and 5 mM (**Figure 5C**) concentration of acidified nitrite for 15, 30, 45 and 60 minutes. Glutathione (GSH), a low molecular weight thiol of Gram-negative bacteria, maintains the reduced state of the cytoplasm. In the presence of external ROS/ RNI stress cytosolic GSH pool is oxidized to form glutathione disulfide (GSSG) [34]. The redox- sensitive GFP2 (roGFP2) having two redox-sensitive cysteine residues at 147th and 204th positions (which form disulfide bond upon oxidation) can absorb at two wavelengths, 405 nm and 488 nm depending upon its oxidized and reduced state, respectively. It has a fixed emission at 510 nm [35]. The glutaredoxin protein (Grx1) fused to the redox-sensitive GFP2 can reversibly transfer electrons between the cellular (GSH/GSSG) pool and the thiol groups of roGFP2 at a much faster rate. The ratio of fluorescence intensity of Grx-roGFP2 measured at 405 nm and 488 nm demonstrates the redox status of the cytoplasm of bacteria [34]. In all the three concentrations of acidified nitrite {800 µM, 1 mM, 5 mM (**Figure 5C**)}, we found a time-dependent increase in the 405/ 488 ratio of Grx1-roGFP2 in STM *ΔompA* strain in comparison with STM (WT), suggesting a heightened redox burst in the cytoplasm of ompA knockout strain in response to RNI in vitro. Taken together, our data indicate the importance of OmpA in maintaining the cytosolic redox homeostasis of wild-type *Salmonella*. Earlier, we have noticed a significant downregulation in the transcript levels of outer membrane-bound larger porins (*ompC*, *ompD*, and *ompF*) compared to *ompA* in wild-type *Salmonella* growing in acidic F media and RAW264.7 cells. Our previous study depicted enhanced nitrite uptake by STM *ΔompA* strain compared to STM (WT), which further indicates increased permeability of the bacterial outer membrane. Hence, we decided to check the expression of larger porins in STM *ΔompA* strain growing in *in vitro* and *ex vivo* conditions. Surprisingly, in contrast to the wild-type bacteria, we have found elevated expression of *ompC*, *ompD*, and *ompF* in STM *ΔompA* strain growing in nutritionally enriched LB media, nutrient-depleted acidic F media (pH= 5.4), and RAW264.7 macrophage cells (**Figure 5D**). This increased expression of larger porins was revoked in the complemented strain. In this regard, the enhanced outer membrane depolarization of STM *ΔompA* growing in acidic F media was further tested (during stationary phase) using a membrane-permeant negatively charged dye called DiBAC_4_ (**Figure 5E**) [36]. Because of the enhanced outer membrane permeability, when the negative charge of the bacterial cytosol is diminished by the inflow of cations (depolarization), the cell allows the entry of DiBAC_4,_ which binds to the cell membrane proteins and starts fluorescing.

Contrary to the wild-type and the complemented strain, the higher DiBAC_4_ positive population and greater median fluorescence intensity of DiBAC_4_ corresponding to STM *ΔompA* and the empty vector strain (**Figure 5E**) ensures enhanced outer membrane permeability of *Salmonella* in the absence of OmpA. We used another porin-specific DNA binding fluorescent dye called bisbenzimide (Figure 5F) [41] to strengthen this observation. In line with our previous observation, we found that the fluorescence intensity of bisbenzimide taken up by STM *ΔompA* growing in acidic F media is more than the wild-type and complemented bacterial strain (**Figure 5F**). Compared to the wild-type bacteria, the greater fluorescence intensity of bisbenzimide corresponding to STM *ΔompA* grown intracellularly in murine macrophages for 12h (**Figure 5G**) firmly endorsed the result obtained from the in vitro experiment. To show explicitly that the increased expression of larger porins such as *ompC*, *ompD*, and *ompF* on bacterial outer membrane enhances the outer membrane porosity, we have expressed *ompC, ompD,* and *ompF* in wild-type *Salmonella* with pQE60 plasmid. We have observed that the increased expression of *ompD* and *ompF* enhanced the permeability of the outer membrane of wild-type *Salmonella* growing overnight in acidic F media (**Figure S4A and S4B**). We have further elevated the expression of *ompC, ompD,* and *ompF* in wild-type *Salmonella* by adding IPTG to the LB broth. Our data suggested that the over-expression of *ompF* in the wild-type *Salmonella* causes massive depolarization (61.62%) of the outer membrane compared to the other porins, namely *ompC* (2.73%) and *ompD* (8.06%) (**Figure S4C and S4D**). Hence, we can conclude that in the absence of *ompA*, the expression of larger porins such as *ompC, ompD,* and *ompF* increases on the outer membrane of *Salmonella.* However, the enhanced outer membrane porosity of the bacteria majorly depends upon the elevated expression of *ompF*.

### The maintenance of the integrity of the SCV membrane inside RAW264.7 macrophages solely depends upon OmpA, not on other larger porins such as OmpC, OmpD, and OmpF

To strengthen our previous observation, we decided to knockout *ompC*, *ompD*, and *ompF* individually in the kanamycin-resistant *ΔompA* background of *Salmonella*. We have generated STM *ΔompA ΔompC*, STM *ΔompA ΔompD,* and STM *ΔompA ΔompF* using chloramphenicol resistant gene cassette (data not shown). We further investigated the intracellular niche of these double knockout bacterial strains in RAW264.7 cells during the late phase of infection (16 h post-infection). In line with our previous finding, compared to the wild-type bacteria, STM *ΔompA* showed poor colocalization with SCV marker LAMP-1 (**Figure 6A and 6B**). The drastic loss of SCV membrane from the surroundings of STM *ΔompA ΔompC*, *ΔompA ΔompD*, and *ΔompA ΔompF* as demonstrated by their poor colocalization with LAMP-1 (**Figure 6B**), indicating the cytosolic localization of these double knockout strains in macrophages during the late phase of infection. The reduced colocalization of STM (WT) expressing LLO (which usually stays in the cytoplasm) with LAMP-1 (**Figure 6A and 6B**) authenticates the cytosolic phenotype of the double knockout strains. To rule out the possibility that the lack of OmpC, OmpD and OmpF also contributes to the cytosolic localization of STM *ΔompA ΔompC*, *ΔompA ΔompD*, and *ΔompA ΔompF* (**Figure 6A and 6B**), we generated single knockout strains of *Salmonella* lacking *ompC*, *ompD*, and *ompF* using the method demonstrated earlier. We observed that the STM *ΔompC*, *ΔompD*, and *ΔompF* strains colocalized with LAMP-1 similarly to the wild-type strain (**Figure 6C and 6D**). Hence, it can be concluded that the maintenance of the vacuolar life of wild-type *Salmonella* depends on OmpA and not on OmpC, OmpD, and OmpF. A decreased recruitment of nitrotyrosine on STM *ΔompC*, *ΔompD*, and *ΔompF* in comparison with STM *ΔompA* while growing intracellularly in RAW264.7 cells further supports the presence of intact SCV membrane around them (**Figure S5A**). The *in vitro* sensitivity of STM *ΔompC*, *ΔompD*, and *ΔompF* against the acidified nitrite was also checked (**Figure S5B**). It was observed that, unlike the *ompA* deficient strain of *Salmonella*, the ability of STM *ΔompC*, *ΔompD*, and *ΔompF* to withstand the bactericidal effect of acidified nitrite is comparable with the wild type bacteria (**Figure S5B**). To further support this observation, we checked nitrite consumption by STM *ΔompC*, *ΔompD*, and *ΔompF* (**Figure S5C**). It was found that in comparison to STM *ΔompA* strain, *ΔompC*, *ΔompD*, and *ΔompF* are more efficient in restricting the entry nitrite (**Figure S5C**), which can be attributed to their better survival in the presence of *in vitro* acidified nitrite (**Figure S5B**).

**Figure 6.**
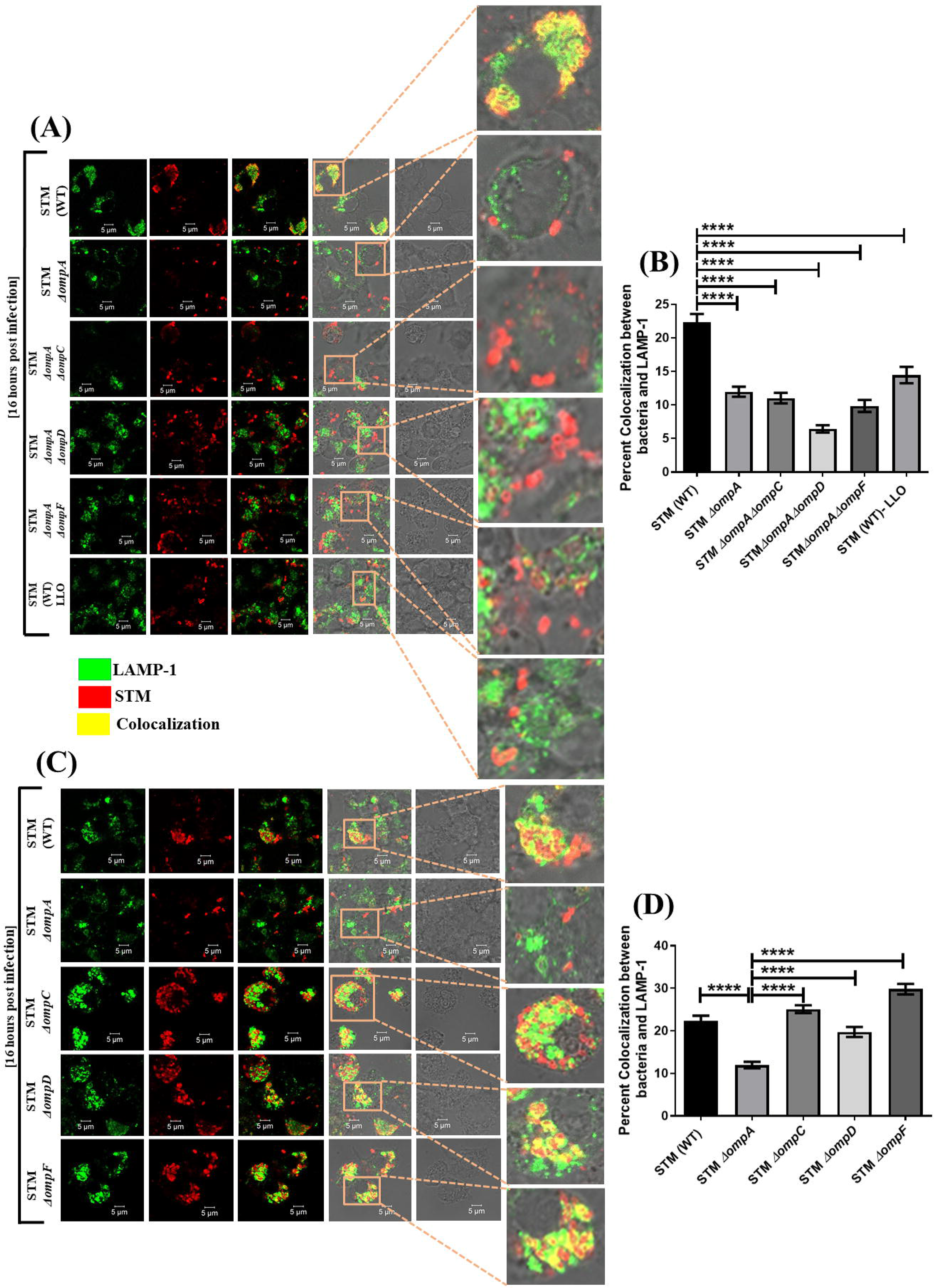
The maintenance of the integrity of the SCV membrane inside RAW264.7 macrophages solely depends upon OmpA, not on other larger porins such as OmpC, OmpD, and OmpF. (A) RAW264.7 cells were infected with STM (WT), *ΔompA*, *ΔompAΔompC*, *ΔompAΔompD*, *ΔompAΔompF*, & WT: *LLO* at MOI 20. Cells were fixed at 16 hours post-infection, followed by labeled with anti-*Salmonella* antibody and anti-mouse LAMP-1 antibody. (B) Quantification of LAMP-1 recruitment on STM (WT), *ΔompA*, *ΔompAΔompC*, *ΔompAΔompD*, *ΔompAΔompF*, (WT): *LLO*. Percent colocalization between bacteria and LAMP-1 was determined after analyzing more than 50 different microscopic stacks from two independent experiments. Data are represented as mean ± SEM (n≥50, N=2). Scale bar = 5μm. (C) RAW264.7 cells were infected with STM (WT), *ΔompA*, *ΔompC*, *ΔompD*, and *ΔompF* at MOI 20. Cells were fixed at 16 hours post-infection, followed by labeled with anti-*Salmonella* antibody and anti-mouse LAMP-1 antibody, respectively. (D) Quantification of LAMP-1 recruitment on STM (WT), *ΔompA*, *ΔompC*, *ΔompD*, and *ΔompF*. Percent colocalization of bacteria with LAMP-1 has been determined after analyzing more than 60 different microscopic stacks from two independent experiments. Data are represented as mean ± SEM (n≥60, N=2). Scale bar = 5μm. *(P)* ****< 0.0001, (Student’s *t* test- unpaired).

### In the absence of OmpA, porins OmpC and OmpF enhance the susceptibility of *Salmonella* against nitrosative stress of RAW264.7 cells

To dissect the role of each larger porin, namely OmpC, OmpD, and OmpF in the entry of nitrite into bacteria during the absence of OmpA, we performed nitrite uptake assay using the double knockout bacterial strains STM *ΔompA ΔompC*, *ΔompA ΔompD*, and *ΔompA ΔompF* (**Figure 7A**). In comparison with the wild-type bacteria, the rapid disappearance of nitrite, which corroborates substantial nitrite uptake from the media by STM *ΔompA* and *ΔompA ΔompD,* demonstrates the involvement of OmpC and OmpF (present in both STM *ΔompA* and *ΔompA ΔompD* strains) in this entry process (**Figure 7A**). This result was further validated by the measurement of the reduced *in vitro* percent viability of STM *ΔompA* and *ΔompA ΔompD* in comparison with wild type, *ΔompA ΔompC* and *ΔompA ΔompF* strains of *Salmonella* in the presence of acidified nitrite (800 µM) (**Figure 7B**). Using the *ex vivo* system of infection in murine macrophages, we have found enhanced recruitment of nitrotyrosine on STM *ΔompA* and *ΔompA ΔompD* as depicted by their higher colocalization in comparison with wild type, *ΔompA ΔompC*, and *ΔompA ΔompF* (**Figure 7C**) *Salmonella*. The intracellular nitrosative burst of macrophages infected with STM (WT), *ΔompA, ΔompA: ompA*, *ΔompA ΔompC, ΔompA ΔompD*, and *ΔompA ΔompF* was tested using DAF2DA (**Figure 7E**). Only 1.94% of macrophages infected with the wild-type bacteria produced [NO], which is comparable to STM *ΔompA: ompA* (1.66%), *ΔompA ΔompC* (1.15%) and *ΔompA ΔompF* (1.62%) infected macrophages population producing [NO]. The greater percent population of DAF2DA positive infected macrophages corresponding to STM *ΔompA* (3.78%) and *ΔompA ΔompD* (3.77%) (**Figure 7E**) strengthens the conclusion obtained from confocal data (**Figure 7C**). The macrophages infected with STM (WT): *LLO* showed a very low intracellular nitrosative burst (0.9%), suggesting that the cytosolic population of wild-type *Salmonella* expressing LLO was protected by the presence of OmpA (**Figure 7E**). This data was further verified by the attenuated intracellular proliferation of STM *ΔompA* and *ΔompA ΔompD* inside RAW264.7 cells (**Figure 7D**). It was found that unlike STM (WT), *ΔompA ΔompC*, and *ΔompA ΔompF*, which are showing very poor colocalization with intracellular nitrotyrosine, the intracellular proliferation of STM *ΔompA* and *ΔompA ΔompD* was severely compromised in murine macrophages (**Figure 7C and 7D**). Taken together, it can be concluded that in the absence of OmpA, the elevated expression of two major larger porins, namely OmpC and OmpF, on the bacterial outer membrane may help in the entry of nitrite into the bacterial cytoplasm and makes the bacteria highly susceptible to the intracellular nitrosative burst. To validate this observation, we have overexpressed *ompC, ompD,* and *ompF* in wild-type *Salmonella* and checked the susceptibility of these bacterial strains against *in vitro* nitrosative stress. We have incubated the wild-type *Salmonella,* overexpressing *ompA, ompC, ompD,* and *ompF* in the presence of acidified nitrite for 12 hours and checked the viability of the bacteria by flow cytometry (propidium iodide staining) (**Figure S6A and S6B**) and resazurin assay (**Figure S6C, S6D, and S6E**). Earlier, we have shown that over-expression of both *ompD* and *ompF* enhances the permeability of the bacterial outer membrane. In line with our previous observation, the flowcytometric data has shown that compared to *ompC,* the over-expression of *ompD* (14.97%) and *ompF* (16.17%) in wild-type *Salmonella* can induce bacterial death in the presence of acidified PBS (**Figure S6A**). However, over-expressing *ompF* in the wild-type *Salmonella* alone is responsible for increasing the susceptibility of the bacteria (22.8%) towards acidified nitrite (**Figure S6A-S6E**). This suggests that out of all three major larger porins, OmpF plays a pivotal role in increasing the susceptibility of *ompA* deficient *Salmonella* against *in vitro* and *in vivo* nitrosative stress.

**Figure 7.**
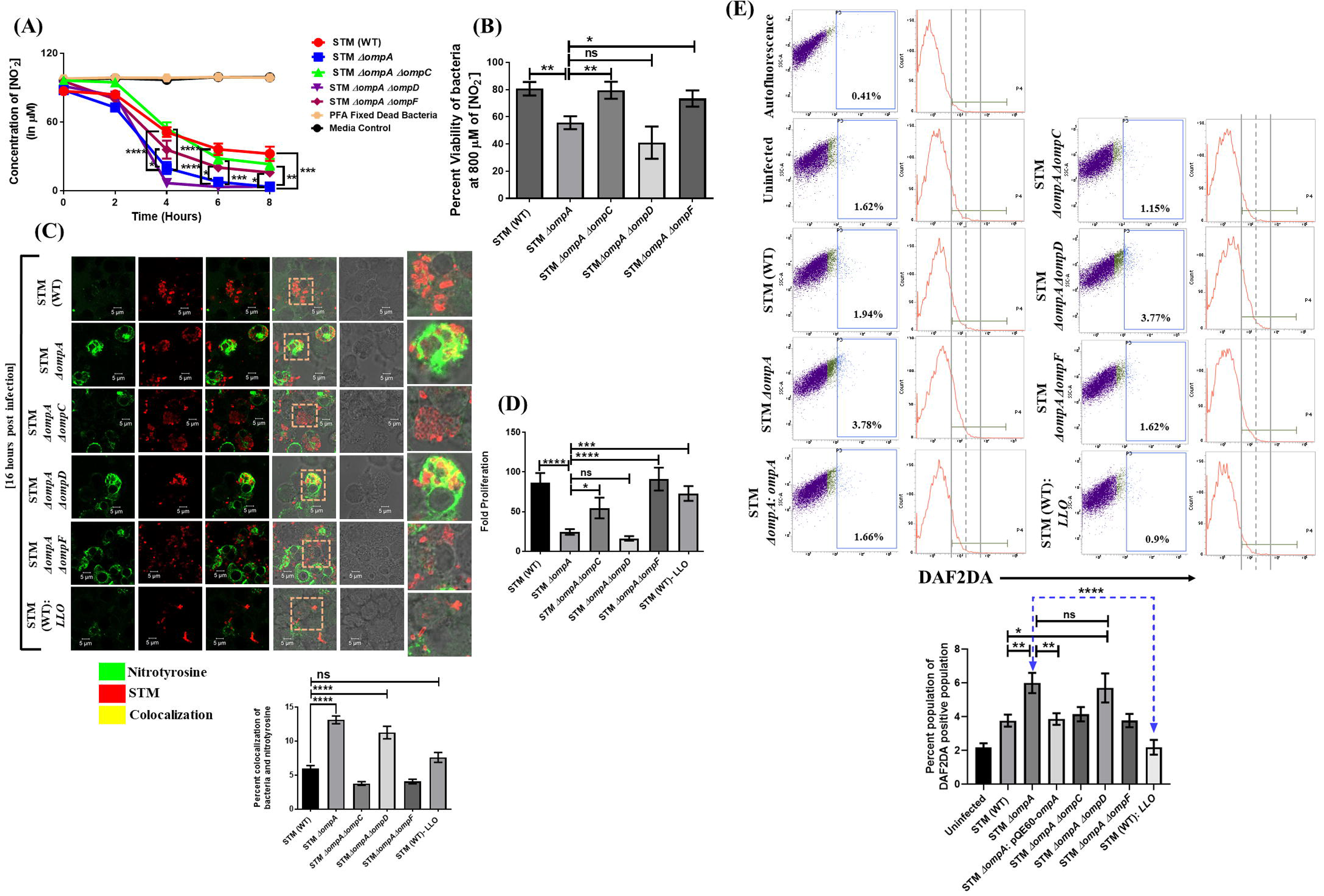
In the absence of OmpA, porins OmpC and OmpF enhance the susceptibility of *Salmonella* against the nitrosative stress of RAW264.7 cells. (A) *In vitro* nitrite uptake assay of STM (WT), *ΔompA*, *ΔompAΔompC*, *ΔompAΔompD*, *ΔompAΔompF*, & PFA fixed dead bacteria. 10^8^ CFU of overnight grown stationary phase culture of all the strains was inoculated in MOPS-NaOH buffer (pH= 8.5) with 100 µM initial nitrite concentration. The remaining nitrite in the media was estimated by Griess assay at 0, 2, 4, 6, 8 hours post-inoculation. Data are represented as mean ± SEM (n=3, N=6). (B) *In vitro* viability assay of STM (WT), *ΔompA*, *ΔompAΔompC*, *ΔompAΔompD*, & *ΔompAΔompF* in the presence of acidified nitrite [800 µM] 12 hours post-inoculation using resazurin solution. Data are represented as mean ± SEM (n=3, N=3). (C) Immunofluorescence image of RAW264.7 cells infected with STM (WT), *ΔompA*, *ΔompAΔompC*, *ΔompAΔompD*, *ΔompAΔompF*, & (WT): *LLO* at MOI 20. Cells were fixed at 16 hours post-infection, followed by labeled with anti-*Salmonella* antibody and anti-mouse nitrotyrosine antibody. Quantification of nitrotyrosine recruitment on STM (WT), *ΔompA*, *ΔompAΔompC*, *ΔompAΔompD*, *ΔompAΔompF*, (WT): *LLO* has been represented in the form of a graph. Percent colocalization between bacteria and nitrotyrosine was determined after analyzing more than 60 different microscopic fields from two independent experiments. Data are represented as mean ± SEM (n≥60, N=3). Scale bar = 5μm. (D) Intracellular survival of STM (WT), *ΔompA*, *ΔompAΔompC*, *ΔompAΔompD*, & *ΔompAΔompF*, & (WT): *LLO* respectively (MOI 10) in RAW264.7 cells. 16- and 2-hours post-infection, the cells were lysed and plated. The CFU at 16 hours was normalized with the CFU at 2 hours to calculate the fold proliferation of bacteria in macrophages. Data are represented as mean ± SEM (n=3, N=3). (E) Estimation of the intracellular nitrite level in RAW 264.7 cells infected with STM (WT), *ΔompA*, *ΔompA*:pQE60-*ompA*, *ΔompAΔompC*, *ΔompAΔompD*, *ΔompAΔompF*, and (WT): *LLO* respectively at MOI 10, 16 hours post-infection using DAF-2DA [5 µM] by flow cytometry. Unstained and uninfected macrophages have also been used as a control. Both dot plots (SSC- A vs. DAF-2 DA) and histograms (Count vs. DAF-2 DA) have been represented. The percent population of DAF-2DA positive macrophages has been represented in the form of a bar graph. Data are represented as mean ± SEM (n≥3, N=5). *(P)* *< 0.05, *(P)* **< 0.005, *(P)* ***< 0.0005, *(P)* ****< 0.0001, ns= non-significant, (Student’s *t* test- unpaired).

**Figure 8.**
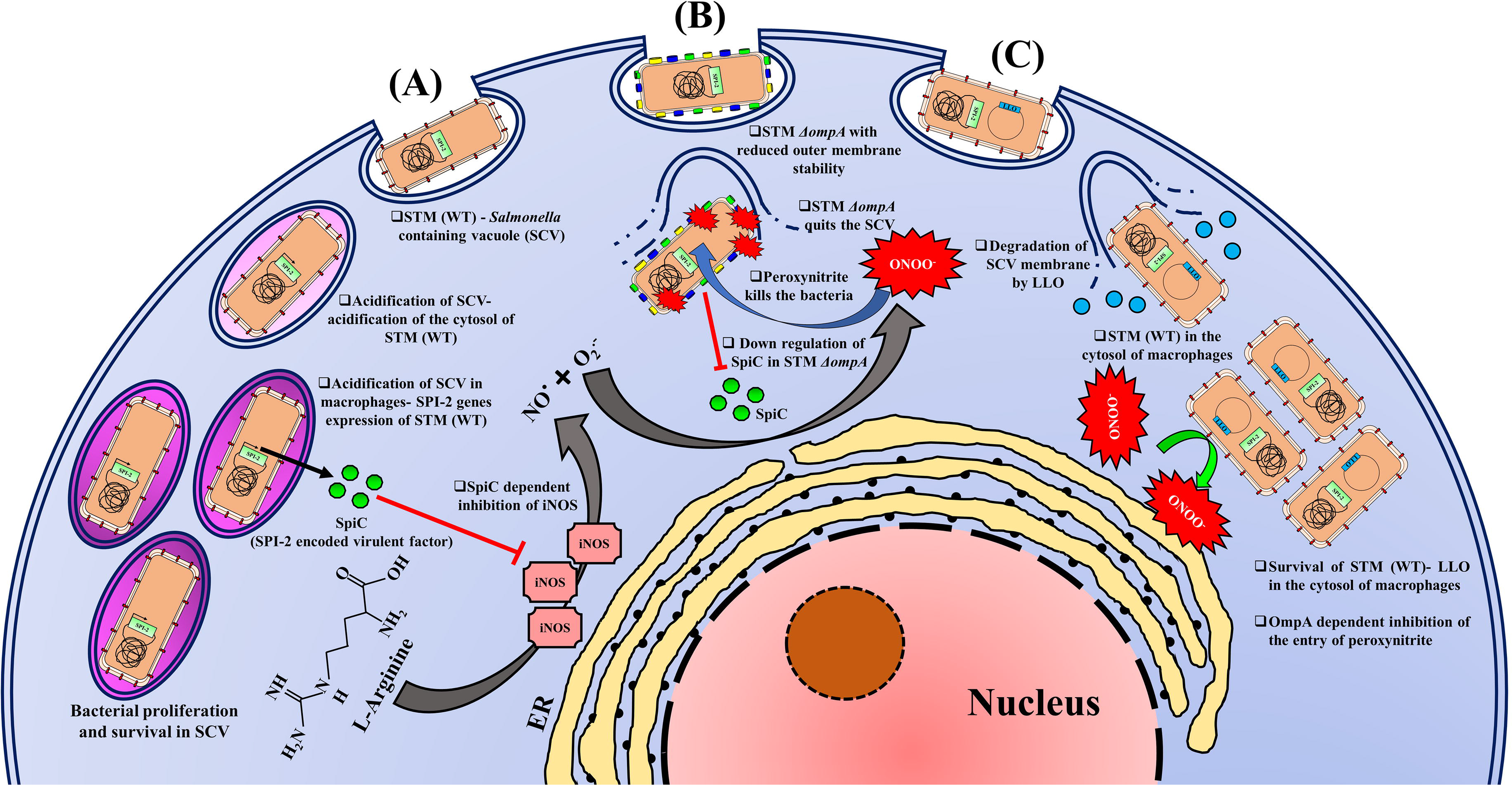
The hypothetical working model of intracellular survival of STM (WT), STM *ΔompA,* and STM (WT): *LLO*. The hypothetical model depicts the fate of (A) STM (WT), (B) STM *ΔompA*, and (C) STM (WT): *LLO* inside the murine macrophages. (A) STM (WT) staying inside the acidic SCV can proliferate efficiently by suppressing the activity of iNOS by SPI-2 encoded virulent factor SpiC. The acidification of the cytosol of wild-type bacteria due to the acidic pH of SCV triggers the expression of SPI-2 genes. (B) STM *ΔompA,* which comes into the cytosol of macrophages after quitting SCV, exhibits an attenuated expression of SpiC because of their stay in a less acidic environment. It is unable to suppress the activity of iNOS and is heavily bombarded with RNI in the cytosol of macrophages. The enhanced outer membrane permeability of the cytosolic population of STM *ΔompA* due to the upregulation of *ompC*, *ompD*, and *ompF* makes them vulnerable towards RNI. (C) STM (WT): *LLO* quits the SCV by expressing LLO. Unlike STM (WT), the cytosolic niche of STM (WT): *LLO* cannot produce SpiC. It can protect itself from RNI by reducing its outer membrane permeability by expressing *ompA* and survive efficiently in the cytosol of macrophages.

## Discussion

Bacterial pathogens (such as *Acinetobacter baumannii*, *E. coli*, *K. pneumoniae,* etc.) can restrict the entry of toxic molecules such as antibiotics and cationic antimicrobial peptides either by changing the outer membrane permeability [37, 38] or by augmenting options/ proteases that degrade the antimicrobial peptides [39]. The alteration in the outer membrane permeability of Gram-negative pathogen is strictly regulated by the core oligosaccharide composition of outer membrane lipopolysaccharide and differential expression of outer membrane porins [40–42]. OmpA, OmpC, OmpD, OmpF, and PhoE, the predominantly found porins on the outer membrane of *Salmonella* Typhimurium, are involved in a wide range of functions. OmpA tightly holds the outer membrane of the bacteria to the peptidoglycan layer of the cell wall, and hence, deletion of OmpA aggravates the biogenesis of outer membrane vesicles (OMV) [7, 43]. OmpD, the most abundant porin on *Salmonella* outer membrane, is involved in the uptake of H_2_O_2_ [44]. OmpC in conjunction with OmpF contributes to the acquisition of cations, whereas PhoE majorly helps in the uptake of negatively charged phosphate groups [8]. The correlation between outer membrane porins and *Salmonella* pathogenesis has been poorly understood due to the lack of extensive studies. Earlier, Heijden *et al.* showed that HpxF^-^ *Salmonella* altered outer membrane permeability by reciprocally regulating the expression of OmpA and OmpC when exposed to H_2_O_2_ [9]. OmpA deficient *Salmonella* was found to be incapable of reaching the mouse brain [45]. Alternative sigma factor of *Salmonella* Typhimurium regulates the expression of OmpA within macrophages to trhive in the presence of oxidative stress [46]. Deletion of *ompD* from the genome of *Salmonella* made it hyper-proliferative in RAW264.7 macrophages and BALB/c mouse models [8]. This study delineated the individual contributions of OmpA, OmpC, OmpD, and OmpF in *Salmonella* pathogenesis. Our study revealed the role of porins in the maintenance of the intravacuolar life of wild-type *Salmonella* Typhimurium and deciphered their role in the regulation of outer membrane permeability and bacterial resistance to the nitrosative stress of macrophages.

We have found a time-dependent steady decrease in the transcript levels of *ompC*, *ompD*, and *ompF* (**Figure S1B, S1C, and S1D**) in *Salmonella* growing in RAW264.7 cells. On the contrary, the consistently elevated expression of *ompA* during the late phase of infection in RAW264.7 cells (9 h and 12 h post-infection) (**Figure S1A**) suggested the importance of this porin in bacterial survival inside the macrophages. Our observations corroborate the findings of Eriksson *et al.* and Hautefort *et al.* [10, 11]. Unlike other significant porins, OmpA has a small pore size and a unique periplasmic domain (**Figure S1A, S1B, S1C, and S1D**), which can act as a gate to restrict the entry of many toxic molecules [9]. Hence it can be concluded that the wild-type bacteria growing in a nutrient-depleted stressed environment of SCV within the macrophages, where the bacterial growth is severely challenged by acidic pH, will prefer OmpA, a porin with a smaller pore size and periplasmic gating mechanism to be expressed on its outer membrane for the survival. OmpA, one of the most abundant porins of the outer membrane of *E. coli* K1, the causative agent of neonatal meningitis, is highly conserved in the family of *Enterobacteriaceae* [4]. Besides its structural role in the bacterial outer membrane, porins interact with host immune cells. OmpA of *E. coli* K1 and *Enterobacter sakazakii* has been reported to be involved in the invasion of hBMEC cells and INT407 cells by multiple studies [47–49]. Earlier, multiple groups reported that an eight-stranded β barrel outer membrane porin (*ompW*) of *Escherichia coli* helps the bacteria evade phagocytosis and confers resistance against alternative complement activation pathway mediated killing by the host [5, 6]. Another study carried out by March C *et al*. suggested that the *ompA* mutant strain of *Klebsiella pneumoniae* is severely attenuated in the pneumonia mouse model [2]. OmpA of *E. coli* K1 not only augments complement resistance by recruiting C4 binding protein (C4BP) on the surface of the bacteria [50] but also aggravates their intracellular survival in murine and human macrophages [51]. In our study, the OmpA deficient strain of *Salmonella* has shown an inclination towards greater phagocytosis and severe attenuation in intracellular proliferation in macrophages. The increased macrophage-mediated intake of complement coated STM *ΔompA* compared to complement coated wild-type bacteria proved that wild-type *Salmonella* Typhimurium impairs the complement activation in OmpA dependent manner. However, the detailed mechanism will be addressed in the future. Surprisingly, the deletion of *ompA* makes *Salmonella* invasion deficient and hyper-proliferative in the epithelial cells. The successful systemic infection of *Salmonella* in macrophages and epithelial cells depends upon its intravacuolar inhabitation. The acidification inside the SCV is a prerequisite for the synthesis and secretion of SPI-2 encoded virulent proteins required for *Salmonella*’s successful proliferation [17]. Mild tampering with the integrity of SCV may create unusual outcomes in the bacterial burden of the cells. Earlier it has been reported that the intracellular proliferation of *sifA* mutant of *Salmonella,* which comes into the cytosol after quitting the SCV, was severely abrogated in macrophages. On the contrary, the bacteria become hyperproliferative in the cytosol of epithelial cells [26]. The introduction of point mutations in the Rab5 or Rab7 proteins (markers of SCV) can also trigger the release of wild-type bacteria from SCV to the cytosol of epithelial cells [52]. Our immunofluorescence microscopy data and the result from chloroquine resistance assay showed the cytosolic localization of ompA deficient strain of *Salmonella* in macrophages and epithelial cells. The wild-type intracellular *Salmonella* recruits SPI-2 encoded translocon proteins SseC and SseD on the surface of SCV to form a functionally active T3SS needle complex. Eventually, wild-type bacteria secret these translocon proteins into the cytosol of the host cells to establish an actively proliferating niche by manipulating the host signaling cascade. The synthesis, surface accumulation, and secretion of SseC and SseD into the host cell cytosol depend upon the acidic pH and integrity of SCV [21]. In line with our expectation, STM *ΔompA* staying in the cytoplasm of macrophages (neutral pH) has exhibited poor colocalization with SseC and SseD. We further checked the expression of *sseC* and *sseD* from intracellularly growing bacteria. We have found that STM *ΔompA* is unable to produce *sseC* and *sseD* like wild-type bacteria, which is the reason behind the poor secretion of SseC and SseD into the cytosol of the macrophages. The cytosolic stay of STM *ΔompA* hampers the acidification of the cytosol of bacteria within macrophages, which further reduces the expression of several SPI-2 encoded virulent genes such as *ssaV* and *sifA.* Earlier, it has been proved that *sifA^-^ Salmonella* comes into the cytosol of the host cells after quitting the SCV and exhibits defective proliferation in the macrophages [53]. Taken together, the result obtained from the intracellular proliferation assay of STM *ΔompA* in macrophages and epithelial cells and the immunofluorescence microscopy data on the vacuolar/ cytosolic status of the bacteria are consistent with the available pieces of literature. Hence, to the best of our knowledge, this is the first report commenting on the role of *Salmonella* Typhimurium outer membrane protein A (OmpA) to maintain the stability of the SCV within macrophages. However, the effect exerted by OmpA in maintaining the integrity of SCV membrane is indirect and dependent on the reduced expression of *sifA*.

The SCV membrane functions as a protective barrier around wild-type bacteria. Once the intactness of the SCV membrane is lost, the bacteria will eventually be exposed to the threats present in the cytosol in the form of ROS and RNI [25]. Earlier, Bonocompain. G *et al.* reported that *Chlamydia trachomatis* infection in HeLa cells transiently induces ROS for the initial few hours of infection [54]. The epithelial cells (HeLa) cannot challenge wild-type *Salmonella* with ROS during infection as efficiently as the macrophages [25]. The generation of RNI indirectly depends upon the ROS burden of a cell. The epithelial cells, a poor producer of ROS, are assumed to be generating a lower level of RNI., which is insufficient to kill the cytosolic population STM *ΔompA* in epithelial cells. On the contrary, the RAW264.7 macrophages can produce both ROS and RNI upon bacterial infection, which might explain the attenuated proliferation of STM *ΔompA* in macrophages. Hence, the oxidative and nitrosative burst of macrophages infected with wild type and the *ompA* knockout strain of *Salmonella* was checked. Surprisingly, we found a remarkable rise in the level of intracellular and extracellular [NO] of macrophages infected with STM *ΔompA*, indicating the protective role of OmpA against the nitrosative stress of macrophages. In continuation with the previous observation, STM *ΔompA,* which has already quit the SCV and stayed in the cytosol of macrophages, showed greater colocalization with nitrotyrosine when compared with the wild-type bacteria protected inside the SCV. To further establish the role of *Salmonella* OmpA against the cytosolic nitrosative stress of macrophages, we have ectopically expressed listeriolysin O (LLO) in wild-type *Salmonella* Typhimurium. The intracellular population of *Listeria monocytogenes*, a causative agent of listeriosis, utilizes LLO to degrade the phagosomal membrane for escaping lysosomal fusion [28, 55]. Expressing LLO in wild-type *Salmonella* will force the bacteria to quit the SCV and be released into the cytosol with intact OmpA on their outer membrane. A decreased recruitment of nitrotyrosine on STM (WT): *LLO* staying in the cytosol of macrophages and their better survival compared to STM *ΔompA* and *ΔompA*: *LLO* proved the role of OmpA in defending the cytosolic population of wild-type *Salmonella* from the harmful effect of RNIs. On the contrary, the higher recruitment of the nitrotyrosine on STM *ΔompA* and STM *ΔompA*: *LLO* can be attributed to their attenuated intracellular proliferation compared to STM (WT) and STM (WT): *LLO.* The alteration in the proliferation of STM *ΔompA* in macrophages and the recruitment profile of nitrotyrosine upon manipulating iNOS activity using specific, irreversible inhibitor 1400W dihydrochloride and activator IFN-ɣ fell in line with the previous results. Wild-type *Salmonella*, with the help of its pathogenicity island (SPI)- 2 encoded virulent factor SpiC activates the suppressor of cytokine signaling 3 (SOCS-3) which inhibit IFN-γ signaling and thus eventually represses the activity of iNOS [29, 30]. As discussed earlier, the acidification of the SCV compartment acidifies the cytoplasm of wild-type *Salmonella*. This process of acidification is essential to synthesize and secret SPI-2 effector proteins into the host cell’s cytoplasm [17]. We have also found that STM *ΔompA,* which stays in the neutral pH of cytosol in macrophages cannot produce SpiC. The decreased expression of *spiC* activates the iNOS in the macrophages infected with STM *ΔompA*. We further checked the promoter activity of *spiC* in wild-type and mutant bacteria growing extracellularly and intracellularly by β galactosidase assay. Compared to the intracellularly growing STM *ΔompA*, the higher activity of *spiC* promoter in the wild-type *Salmonella* simultaneously supports the cytosolic inhabitation of STM *ΔompA* and answers the reason behind the nitrosative burst of macrophages. The downregulation of SPI-2 encoded gene *spiC* in STM *ΔompA* can also be attributed to its incompetence to acidify its cytosol in response to extracellular acidic response, which was determined by a higher 488 nm/ 405 nm ratio of BCECF-AM. But strikingly, STM *ΔompA* was unable to induce intracellular and extracellular ROS production while infecting macrophages. NADPH phagocytic oxidase is the key enzyme of macrophages involved in the synthesis of superoxide ions. It is a multimeric protein consisting of two membrane-bound subunits such as gp91*^phox^*, p22*^phox^,* and four cytosolic subunits such as p47*^phox^*, p40*^phox^*, p67*^phox,^* and RacGTP [56]. Continuing with our previous observations, we found poor colocalization of gp91*^phox^* with STM (WT) and *ΔompA* in macrophages during the late phase of infection (data not shown). Wild-type *Salmonella* can inhibit the recruitment of NADPH oxidase on the surface of the SCV membrane with the help of the SPI-2 encoded type 3 secretion system [57]. The restriction in the recruitment of NADPH oxidase on the surface of the *ompA* knockout bacteria lacking SCV membrane and staying in the cytosol is the probable reason behind the dampened oxidative stress inside the infected macrophages. On the contrary, the ability of iNOS to maintain its uninterrupted catalytic activity, despite being recruited on the cortical actin of macrophages [58], is the most probable reason behind elevated production of RNI in RAW264.7 cells infected with STM *ΔompA*. To investigate the role of OmpA in the establishment of *in vivo* infection by wild-type *Salmonella* Typhimurium we have used 4 to 6 weeks old *Nramp^-/-^* C57BL/6 and BALB/c mice. The better survival of the mice infected with *ompA* deficient strains of *Salmonella*, which was attributed to the reduced bacterial burden in the liver, spleen, and MLN, finally endorsed the essential role of OmpA in the *in vivo* infection of *Salmonella*. Administration of iNOS inhibitor aminoguanidine hydrochloride by intraperitoneal injection or direct oral infection of *iNOS^-/-^* mice diminished the *in vivo* attenuation of STM *ΔompA,* suggesting OmpA dependent protection of wild type bacteria against nitrosative stress *in vivo*. The comparable CFU burden of STM (WT) and *ΔompA* in *gp91phox^-/-^* C57BL/6 mice, which cannot produce ROS, can be accounted for the abrogated peroxynitrite response [59, 60]. Acidified nitrite generating a wide range of reactive nitrogen intermediates (RNI) such as nitrogen dioxide (NO_2_), dinitrogen trioxide (N_2_O_3_), nitric oxide (NO), etc., are extensively used for assessing *in vitro* viability of bacteria and fungi [32, 61]. After entering the bacterial cells, RNIs cause enormous irreparable damages to multiple subcellular components such as nucleic acids (cause deamination of nucleotides), proteins (disruption of Fe-S clusters, heme group; oxidation of thiols and tyrosine residues), lipids (cause peroxidation), etc., and eventually destroy the pathogen. The enhanced sensitivity of STM *ΔompA* against acidified nitrite and combination of acidified nitrite with peroxide, which generates peroxynitrite [62], suggested increased entry of nitrite in the knockout strain comparison with wild type bacteria. The faster depletion of nitrite from the media having STM *ΔompA* strongly supported our hypothesis. The time-dependent gradual increase in the 405/ 488 ratio of pQE60-Grx1-roGFP2 harbored by STM *ΔompA* under acidified nitrite’s tested concentrations proved the loss of redox homeostasis in STM *ΔompA*.

As discussed earlier, the outer membrane of Gram-negative bacteria acts as an impenetrable barrier to many toxic compounds (including antibiotics, bile salts, cationic antimicrobial peptides, reactive oxygen species, abnormal pH, osmotic stress, etc.) and protect the bacteria from environmental threats during its extracellular and intracellular life-cycle. The enhanced uptake of nitrite by *Salmonella* in the absence of OmpA and a time-dependent increase in the sensitivity of STM *ΔompA* towards acidified nitrite proves the occurrence of a permanent damage to the outer membrane of the bacteria. The increased uptake of DiBAC_4_ (measure the cell membrane depolarization) and bisbenzimide (binds to the DNA) by the *ompA* deficient strain of *Salmonella* provided further supports to our conclusion. To rationalize the enhanced uptake of fluorescent probes by STM *ΔompA,* we checked the expression of larger porins such as *ompC*, *ompD*, and *ompF* in wild-type, mutant, and complemented strains of *Salmonella* growing in LB broth, F media, and RAW264.7 macrophages. We found a remarkable increase in the expression of the larger porins in the knockout strain (lacking OmpA) compared to wild-type and complemented strains of *Salmonella*. Hence, we have concluded that OmpA regulates the outer membrane stability of *Salmonella* Typhimurium. When OmpA is deleted, the increased expression of larger porins reduces the stability of the outer membrane and makes it permeable to nitrites, which eventually kills the bacteria. To best of our knowledge, this is the first report where we have experimentally dissected the role of OmpA in maintaining outer membrane stability of *Salmonella* Typhimurium. To strengthen our conclusion, we have further overexpressed *ompC*, *ompD*, and *ompF* in the wild-type *Salmonella* with the help of a low copy number plasmid pQE60 that has a site for ‘lac operator.’ We over-expressed these larger porins in wild-type bacteria after incubating them with appropriate antibiotics and 500 µM of IPTG. Compared to other larger porins, the remarkably improved uptake of DiBAC_4_ by wild-type *Salmonella* expressing *ompF* pinpoints the contribution of OmpF in escalating the reduction of outer membrane stability in the absence of OmpA. This conclusion can be further extrapolated to understand the reason behind the enhanced recruitment of nitrotyrosine on STM *ΔompA* compared to STM (WT): *LLO* in macrophages during the late phase of infection. Despite having cytosolic niche STM (WT): *LLO* has intact OmpA in its outer membrane, unlike STM *ΔompA*, which maintains the integrity and stability of the outer membrane and prohibits the entry of peroxynitrite. This further proves that the outer membrane defect of STM *ΔompA* makes the bacteria accessible to RNI produced by the macrophages.

To ascertain the individual contribution of the larger porins such as *ompC*, *ompD*, and *ompF* in the entry of nitrite in *ompA* deficient bacteria, we constructed *ompC*, *ompD*, and *ompF* single and double knockout strains in wild type and *ompA*^-^ background of *Salmonella*, respectively. The cytosolic localization of double knockout strains, namely STM *ΔompA ΔompC*, *ΔompA ΔompD*, and *ΔompA ΔompF* and vacuolar imprisonment of the single knockout strains such as STM *ΔompC*, *ΔompD*, and *ΔompF* provided vital support to our previous conclusion on the dependence of SCV integrity and stability on OmpA. Under the *in vitro* challenge of acidified nitrite, the better survival of STM *ΔompC*, *ΔompD*, and *ΔompF* in comparison with *ΔompA* not only suggested the abrogated consumption of nitrite but also indisputably established the paramount importance of OmpA in the maintenance of the outer membrane permeability of wild-type *Salmonella.* STM *ΔompA ΔompD,* which possesses intact OmpC and OmpF on their outer membrane, showed enhanced nitrite consumption and higher sensitivity towards *in vitro* nitrosative stress (800 µM of acidified NaNO_2_). In comparison with STM *ΔompA ΔompC* and *ΔompA ΔompF,* greater recruitment of nitrotyrosine on the cytosolic population of STM *ΔompA ΔompD* (having OmpC and OmpF) due to the significant loss of outer membrane stability is considered as the sole reason behind their poor proliferation in murine macrophages. These results were further supported by the inability of STM *ΔompA ΔompC* and *ΔompA ΔompF* to induce heightened [NO] response while staying inside the macrophages, unlike STM *ΔompA ΔompD*. To strongly endorse this result, we have decided to verify the viability of wild-type *Salmonella* against the *in vitro* nitrosative stress upon expressing *ompC*, *ompD*, and *ompF*. In line with our previous observations, we have found that compared to *ompD* and *ompF,* the over-expression of *ompF* in wild-type *Salmonella* drastically increases the bacteria’s susceptibility towards acidified nitrites. These results collectively suggest the pivotal role of OmpF in the entry of nitrite in *ompA* deficient *Salmonella* Typhimurium by increasing the permeability and worsening the integrity of the bacterial outer membrane. In this context, we must mention that STM *ΔompA ΔompC* is also expected to express OmpD and OmpF on its outer membrane. Earlier, we have shown that compared to OmpC, OmpD and OmpF contribute more in the depolarization of the bacterial outer membrane. However, OmpF is solely responsible for the death of the bacteria in the presence of *in vitro* nitrosative stress. Despite having OmpF on their outer membrane, the better survival of STM *ΔompA ΔompC* compared to STM *ΔompA ΔompD* in response to *in vitro* and e*x vivo* nitrosative stress might be questioned by the conclusions of our study, which we will answer in the future.

To summarize, our study claims OmpA of *Salmonella* Typhimurium to be a versatile protein with a multitude of activities. The deletion of *ompA* from *Salmonella* interrupts the stability of SCV and imposes significant paradoxical consequences on the intracellular proliferation of bacteria in the macrophages and epithelial cells, respectively. We have experimentally proved that OmpA maintains the stability of the bacterial outer membrane. In the absence of OmpA, the porosity of the outer membrane increases, which makes the bacteria vulnerable to *in vitro* and *in vivo* nitrosative stress. We proposed an OmpA dependent mechanism that regulated the stability of the bacterial outer membrane and was employed cleverly by *Salmonella* to fight against the nitrosative stress of murine macrophages.

## Abbreviations

STM: *Salmonella* Typhimurium
OmpA: Outer membrane protein A
OmpC: Outer membrane protein C
OmpD: Outer membrane protein D
OmpF: Outer membrane protein F
LLO: Listeriolysin O
SCV: *Salmonella* containing vacuole
LAMP-1: Lysosome associated membrane protein-1
iNOS: Inducible nitric oxide synthase
RNI: Reactive nitrogen intermediates
ROS: Reactive oxygen species
RFP: Red fluorescent protein
roGFP2: Redox sensitive green fluorescent protein

## Materials and Methods

### Bacterial strains, media, and culture conditions

The wild type (WT) bacteria *Salmonella enterica* serovar Typhimurium [STM- (WT)] strain 14028S used in this study was a generous gift from Professor Michael Hensel, Max Von Pettenkofer-Institute for Hygiene und Medizinische Mikrobiologie, Germany. The bacterial strains were revived from glycerol stock (stored in −80°C) and plated either on LB agar (purchased from HiMedia) or LB agar along with appropriate antibiotics like-kanamycin (50 μg/mL), ampicillin (50 μg/mL), Chloramphenicol (25 μg/mL), kanamycin and ampicillin (both 50 μg/mL), Kanamycin and Chloramphenicol (Kanamycin= 50 μg/mL, Chloramphenicol= 25 μg/mL) for wild type, knockout (single and double), complement and mCherry expressing strains respectively. *Salmonella-Shigella* agar was used to plating cell lysates/ cell suspensions to calculate the bacterial burden in infected cell lines and several organs of infected mice. Bacterial LB broth cultures were grown in a shaker incubator at 180 rpm, either 37°C for typical wild type and knockout strains or 30°C for the strains having temperature-sensitive pKD46 plasmid or the strains harboring pQE60-Grx1-roGFP2 plasmid and undergoing IPTG (concentration= 500 µM) treatment. For growth curve experiments and *in vitro* RNA extraction studies, a single colony was inoculated in 5mL of LB broth and grown overnight with or without antibiotics at 37°C. Overnight-grown stationary phase bacteria were sub-cultured at a ratio of 1: 100 in freshly prepared LB or minimal F media (acidic) and kept in a 37°C shaker incubator. At different time intervals, aliquots were taken for RNA isolation, serial dilution, plating, and [OD]600nm measurement by TECAN 96 well microplate reader. The complete list of strains and plasmids has been listed below. (Description in Table- 1) Dead bacteria used in several experiments were produced from viable wild-type bacteria either heating at 650C for 20 minutes or treating the bacteria with 3.5% paraformaldehyde for 30 minutes.

**Table 1.** Strains and plasmids used in this study.

| Strains/ plasmids | Characteristics | Source/ references |
| --- | --- | --- |
| *Salmonella enterica* serovar Typhimurium ATCC strain 14028S | Wild type (WT) | Gifted by Prof. M. Hensel |
| *S. Typhimurium* $\Delta ompA$ | Kan<sup>R</sup> | This study |
| $\Delta ompA$: pQE60-*ompA* | Kan<sup>R</sup>, Amp<sup>R</sup> | This study |
| $\Delta ompA$: pQE60 | Kan<sup>R</sup>, Amp<sup>R</sup> | This study |
| *S. Typhimurium* $\Delta ompC$ | Chl<sup>R</sup> | This study |
| *S. Typhimurium* $\Delta ompD$ | Chl<sup>R</sup> | This study |
| *S. Typhimurium* $\Delta ompD$ | Kan<sup>R</sup> | This study |
| *S. Typhimurium* $\Delta ompF$ | Chl<sup>R</sup> | This study |
| *S. Typhimurium* $\Delta ompA$ $\Delta ompC$ | Kan<sup>R</sup>, Chl<sup>R</sup> | This study |
| *S. Typhimurium ΔompA*<br>*ΔompD* | Kan<sup>R</sup>, Chl<sup>R</sup> | This study |
| *S. Typhimurium ΔompA*<br>*ΔompF* | Kan<sup>R</sup>, Chl<sup>R</sup> | This study |
| pKD4 | Plasmid with FRT-flanked<br>Kanamycin resistance gene | KA Datsenko & BL<br>Wanner, PNAS, 2000 |
| pKD46 | Plasmid expressing λ red<br>recombinase, Amp<sup>R</sup> | KA Datsenko & BL<br>Wanner, PNAS, 2000 |
| pQE60 vector | Low copy number plasmid,<br>Amp<sup>R</sup> | Laboratory stock |
| pFV- mCherry (RFP) | Amp<sup>R</sup> | Laboratory stock |
| pFV: GFP | Amp<sup>R</sup> | Laboratory stock |
| STM (WT): *LLO* | Amp<sup>R</sup> | Laboratory stock |
| STM *ΔompA*: *LLO* | Amp<sup>R</sup> | This study |
| *S. Typhimurium* wild type:<br>pHG86 *spiC-LacZ* | Amp<sup>R</sup> | This study |
| *S. Typhimurium ΔompA*:<br>pHG86 *spiC-LacZ* | Amp<sup>R</sup> | This study |
| *S. Typhimurium* wild-type:<br>pQE60-*ompA* | Amp<sup>R</sup> | This study |
| *S. Typhimurium* wild-type:<br>pQE60-*ompC* | Amp<sup>R</sup> | This study |
| *S. Typhimurium* wild-type:<br>pQE60-*ompD* | Amp<sup>R</sup> | This study |
| *S. Typhimurium* wild-type:<br>pQE60-*ompF* | Amp<sup>R</sup> | This study |
| *S. Typhimurium* wild-type:<br>pQE60 | Amp<sup>R</sup> | This study |
| pHG86 *spiC-LacZ* | Amp<sup>R</sup> | Laboratory stock |
| pQE60-Grx1-roGFP2 | Amp<sup>R</sup> | Gifted by Dr. Amit Singh,<br>CIDR, IISc |

### Eukaryotic cell lines and growth conditions

The murine macrophage-like cell line RAW 264.7, human cervical adenocarcinoma cell line HeLa, human colorectal adenocarcinoma cell line Caco-2 were maintained in Dulbecco’s Modified Eagle’s Media (Sigma-Aldrich) supplemented with 10% FCS (Fetal calf serum, Gibco) at 37°C temperature in the presence of 5% CO_2_. Human monocyte cell line U937 cells were maintained in Roswell Park Memorial Institute 1640 media (Sigma-Aldrich) supplemented with 10% FCS (Fetal calf serum, Gibco). For polarizing the Caco-2 cells, DMEM media was further supplemented with 1% non-essential amino acid solution (Sigma-Aldrich). Phorbol Myristate Acetate (Sigma-Aldrich) (concentration- 20 ng/ mL) was used for the activation of U937 cells for 24 hours at 37°C temperature in the presence of 5% CO_2_, followed by the replacement of the media carrying PMA with normal RPMI supplemented with 10% FCS and further incubating the cells for 24 hours before starting the experiments.

### Construction of *ompA, ompC*, *ompD*, & *ompF* knockout strains of *Salmonella*

The knockout strains of *Salmonella enterica* serovar Typhimurium (strain 14028S) were made using one step chromosomal gene inactivation method demonstrated by Datsenko and Wanner [13]. Briefly, STM (WT) was transformed with pKD46 plasmid, which has a ‘lambda red recombinase system’ under arabinose inducible promoter. The transformed cells were grown in LB broth with ampicillin (50 μg/mL) and 50 mM arabinose at 30°C until the [OD]_600nm_ reached 0.35 to 0.4. Electrocompetent STM pKD46 cells were prepared after washing the bacterial cell pellet thrice with double autoclaved chilled Milli Q water and 10% (v/v) glycerol. Finally, the electrocompetent STM pKD46 cells were resuspended in 50 μL of 10% glycerol. Kanamycin resistant gene cassette (Kan^R^, 1.6kb- for knocking out *ompA, ompD*) and chloramphenicol resistant gene cassette (Chl^R^, 1.1 kb- for knocking out *ompC, ompD* & *ompF*) were amplified from pKD4 and pKD3 plasmids, respectively using knockout primers (Table- 3.2). The amplified Kan^R^ and Chl^R^ gene cassettes were subjected to phenol-chloroform extraction and electroporated into STM (WT) pKD46. The transformed cells were plated on LB agar with kanamycin (50 μg/mL) for selection of *ΔompA, ΔompD* strains and LB agar with chloramphenicol (25 μg/mL) for selection of *ΔompC, ΔompD,* and *ΔompF* strains. The plates were further incubated overnight at 37°C. The knockout colonies were confirmed by confirmatory and kanamycin/ chloramphenicol internal primers (Table- 2).

**Table 2.**
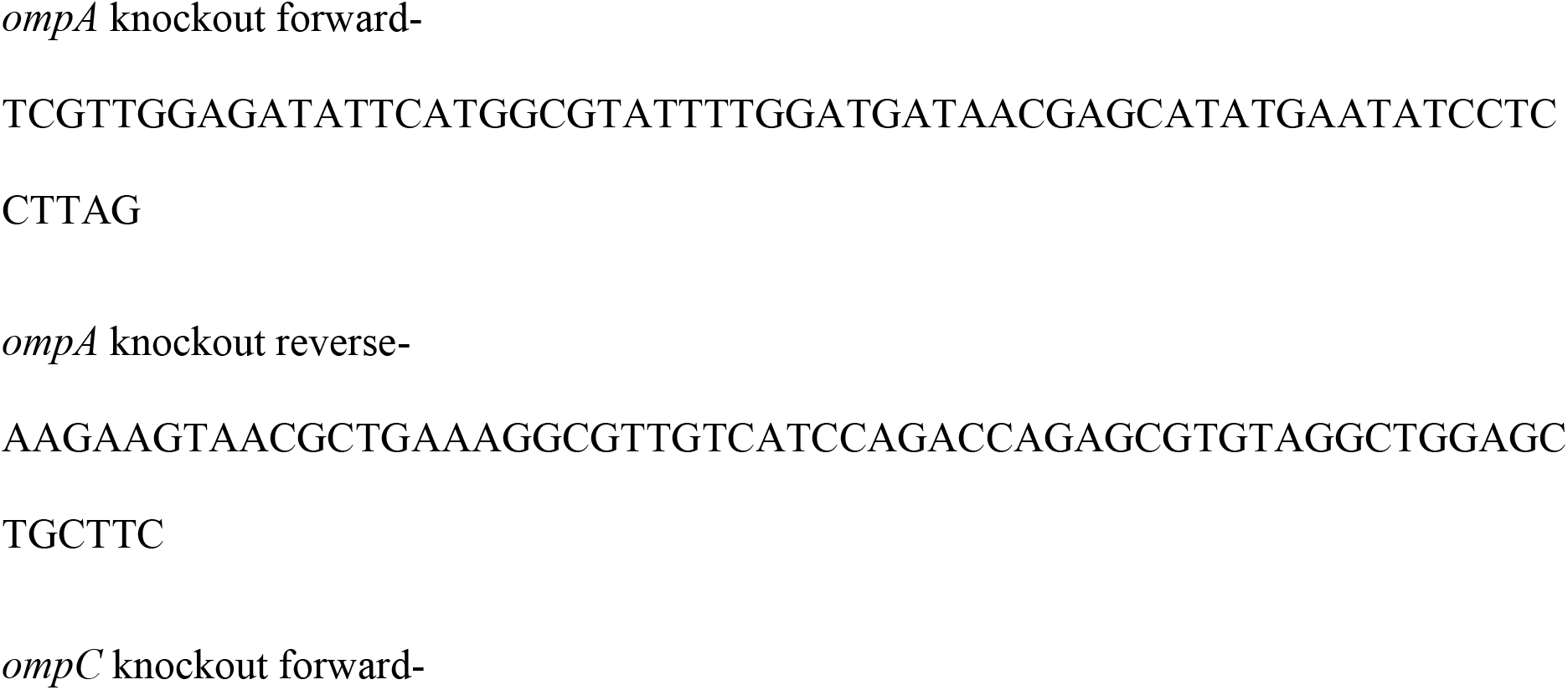
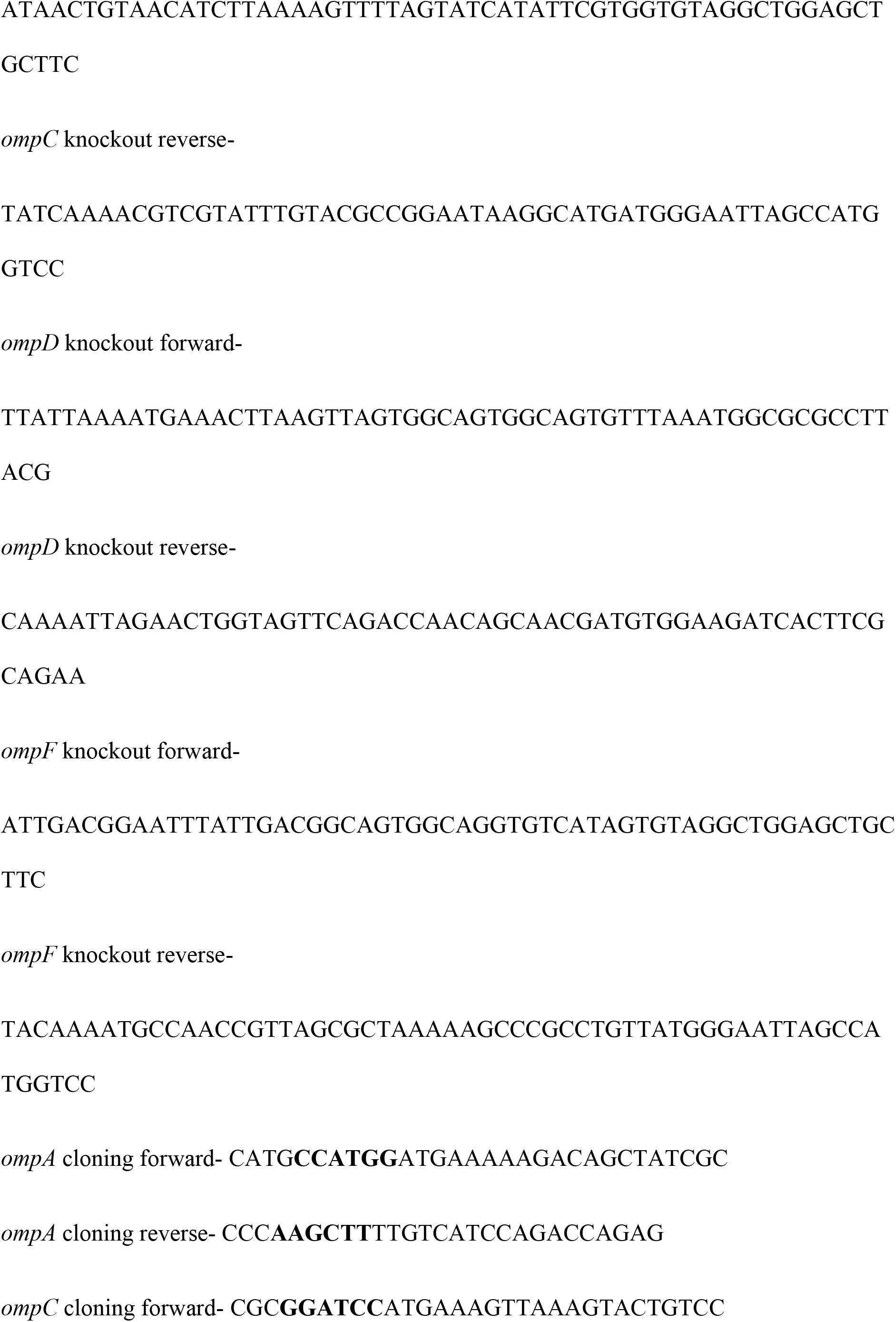
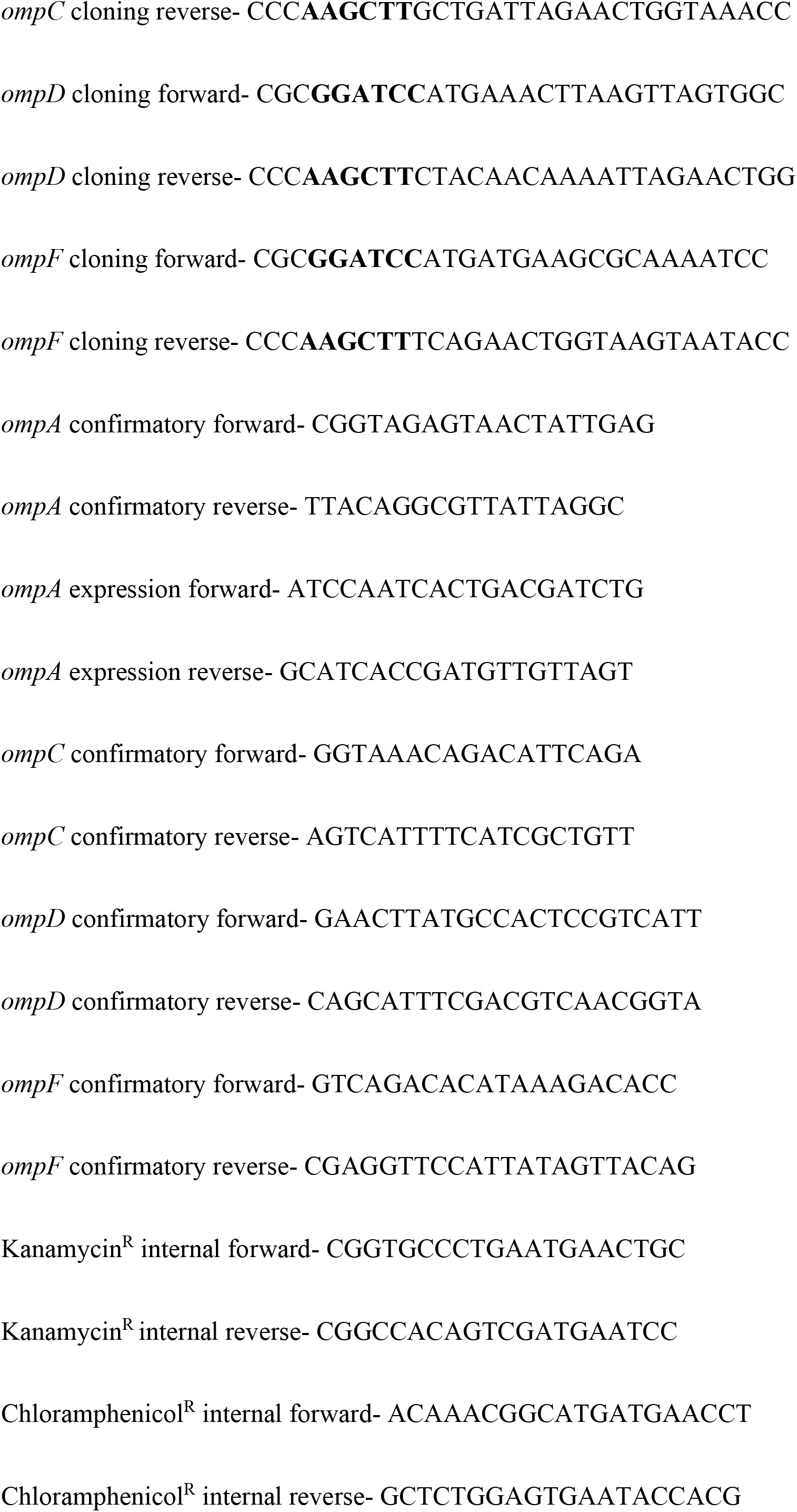
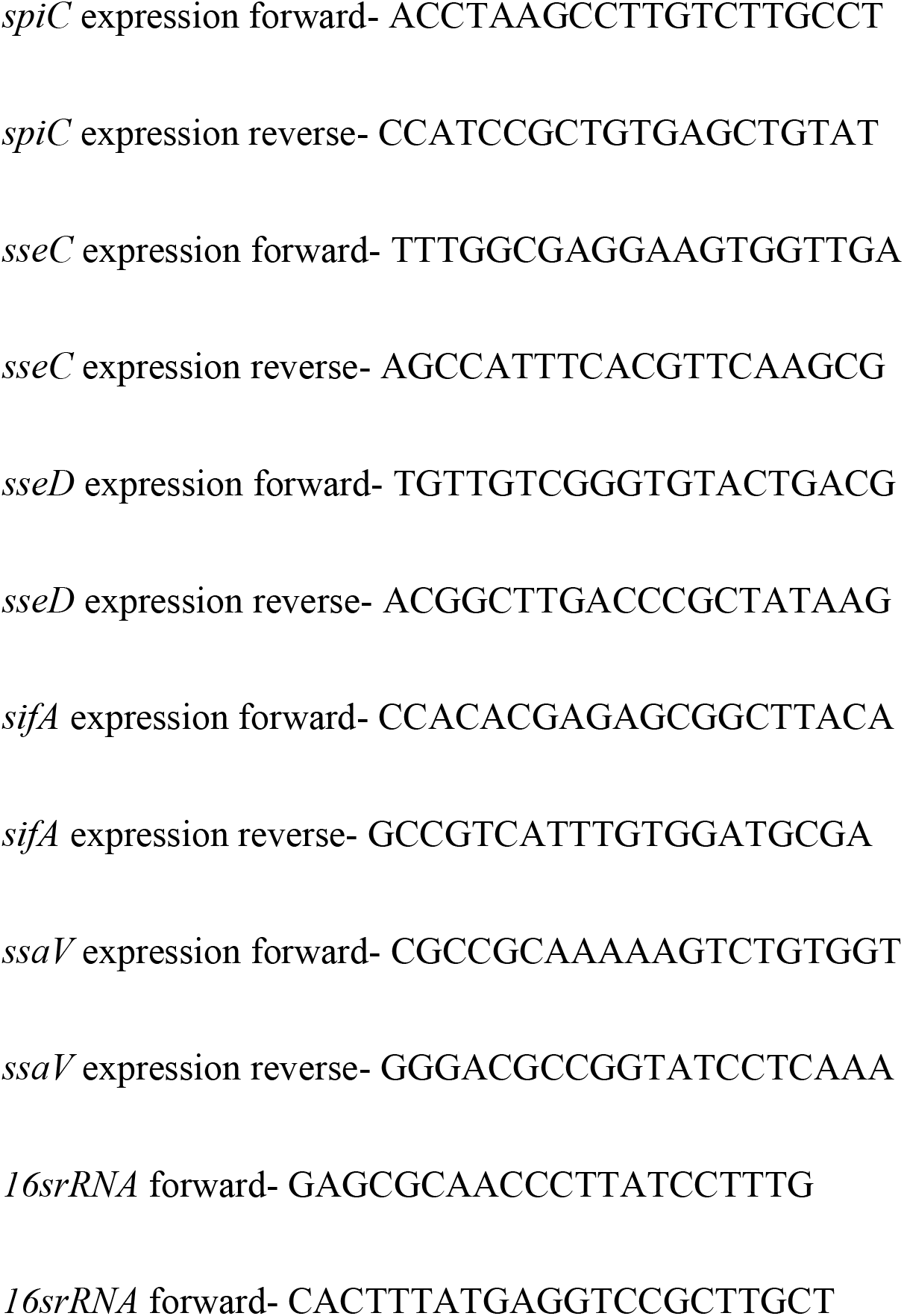
Primer sequences (5’ to 3’)

### Construction of *ompA, ompC, ompD, ompF* complemented strain of *Salmonella*

The *ompA, ompC, ompD,* and *ompF* genes were amplified with their respective cloning primers (Description in Table- 3.2) by colony PCR. The amplified PCR products, purified by phenol-chloroform extraction, was subjected to restriction digestion by specific restriction enzymes such as NcoI (NEB) and HindIII (NEB) for *ompA*, BamHI (NEB), and HindIII (NEB) for *ompC, ompD,* and *ompF* in the CutSmart buffer (NEB) at 37°C for 2-3 h along with the empty pQE60 vector backbone. Double digested insert and vector were subjected to ligation by a T_4_ DNA ligase in 10X ligation buffer (NEB) overnight at 16°C. The ligated products and the empty vector were transformed into respective bacterial strains separately to generate complemented, over-expression, and empty vector strains. Complementation and over- expression were initially confirmed by colony PCR with cloning and *ompA, ompC, ompD,* and *ompF* internal primers (data not shown) and finally by restriction digestion of recombinant plasmid. The expression level of *ompA* in the knockout, complemented, and empty vector strains were further confirmed by RT PCR using *ompA* specific RT primers (Description in Table- 2).

### Construction of *ΔompA ΔompC*, *ΔompA ΔompD*, & *ΔompA ΔompF* double knockout strains of *Salmonella*

The double knockout strains of *Salmonella enterica* serovar Typhimurium (strain 14028S) were made by slightly modifying the one-step chromosomal gene inactivation strategy demonstrated by Datsenko and Wanner [13]. Briefly, STM *ΔompA*, where the *ompA* gene has already been replaced with a kanamycin-resistant gene cassette, was transformed with a pKD46 plasmid. The transformed cells were grown in LB broth with ampicillin (50 μg/mL) and 50 mM arabinose at 30°C until [OD]_600nm_ reached 0.35 to 0.4. Electrocompetent STM *ΔompA* pKD46 cells were made using the protocol mentioned above. Chloramphenicol resistant gene cassette (Chl^R^, 1.1 kb) was amplified from pKD3 plasmid using knockout primers having a stretch of oligos at the 5’ end homologous to the flanking region of the target genes- *ompC*, *ompD*, *ompF* (Table- 3.2). The amplified Chl^R^ gene cassette was subjected to phenol-chloroform extraction and electroporated into STM *ΔompA* pKD46. The transformed cells were plated on LB agar with kanamycin (50 μg/mL) and chloramphenicol (25 μg/mL) both for selection of double knockout strains (*ΔompA ΔompC*, *ΔompA ΔompD*, and *ΔompA ΔompF*). The plates were further incubated overnight at 37°C. The knockout colonies were confirmed by confirmatory primers (Table- 2).

### RNA isolation and RT PCR

The bacterial cell pellets were lysed with TRIzol reagent (RNAiso Plus, Takara) and stored at −80°C overnight. The lysed supernatants were further subjected to chloroform extraction followed by precipitation of total RNA by adding an equal volume of isopropanol. The pellet was washed with 70% RNA-grade ethanol, air-dried, and suspended in 20 μL of DEPC treated water. RNA concentration was measured in nano-drop and analyzed on 1.5% agarose gel to assess RNA quality. To make cDNA, 3 μg of RNA sample was subjected to DNase treatment in the presence of DNase buffer (Thermo Fischer Scientific) at 37°C for 2h. The reaction was stopped by adding 5mM Na_2_EDTA (Thermo Fischer Scientific), followed by heating at 65°C for 10 min. The samples were incubated with random hexamer at 65°C for 10 min and then supplemented with 5X RT buffer, RT enzyme, dNTPs, and DEPC treated water at 42°C for 1h. Quantitative real-time PCR was done using SYBR/ TB Green RT PCR kit (Takara Bio) in Bio-Rad real-time PCR detection system. The expression level of target genes was measured using specific RT primers (Table- 2). 16S rRNA was used to normalize the expression levels of the target genes.

### Percent phagocytosis calculation/ invasion assay

1.5 to 2 × 10^5^ cells (RAW264.7, U937, Caco-2, and HeLa) were seeded into the wells of 24 well plates and incubated for 6-8 h at 37°C in the presence of 5% CO_2_. As demonstrated earlier, the protocol for calculating percent phagocytosis by macrophage cells has been modified a little [63]. Briefly, the phagocytic macrophage cells (RAW264.7 and activated U937 cells) were infected with 10- 12 h grown stationary phase cultures of STM (WT), *ΔompA, ΔompA*: pQE60-*ompA* and *ΔompA*: pQE60 at MOI 10. For assays with the complemented strains, the strains were incubated with 10% mouse complement sera for 2h before the experiment, and MOI 50 was used for the infection. Non-phagocytic epithelial cells (Caco-2 and HeLa cells) were infected with the cells from 3 to 4 h old log phase culture of all four bacterial strains at MOI 10. The attachment of bacteria to the cell surface was increased by centrifuging the cells at 800 rpm for 5 min, followed by incubating the infected cells at 37°C in the presence of 5% CO_2_ for 25 min. Next, the cells were washed thrice with PBS to remove unattached bacteria and subjected to 100 μg/ mL and 25 μg/ mL concentration of gentamycin treatment for 1h each. 2 h post-infection, the cells were lysed with 0.1% triton-X-100. The lysate was plated on *Salmonella- Shigella* agar, and the corresponding CFUs were enumerated. Percent phagocytosis (for phagocytic macrophage cells)/ percent invasion (for non-phagocytic epithelial cells) was determined using the following formula-

### Percent phagocytosis/ percent invasion= [CFU at 2 h]**/** [CFU of pre-inoculum] × 100 Adhesion assay

The protocol of adhesion assay was as described before [14]. Briefly, 1.5 to 2 × 10^5^ cells (RAW264.7 and HeLa) were seeded on the top of sterile glass coverslips. Phagocytic macrophage cells (RAW264.7) were infected with 10- 12 h old stationary phase culture of STM (WT), *ΔompA, ΔompA*: pQE60-*ompA,* and *ΔompA*:pQE60 at MOI 50. Non-phagocytic epithelial cells (HeLa) were infected with 3-4 hours old log phase culture of all four bacterial strains at the same MOI. After centrifuging the cells at 800 rpm for 5 min, the infected cells were incubated at 37°C temperature in the presence of 5% CO_2_ for 15 minutes (for RAW264.7 cells) and 25 minutes (for HeLa cells), respectively. After the incubation period, the cells were washed twice with sterile PBS and fixed with 3.5% PFA. To visualize the externally attached bacteria, the cells were primarily treated with anti-*Salmonella* antibody raised in rabbit (dilution 1: 100, duration 6 to 8 hours at 4°C temperature), which was followed by the treatment of the cells with secondary antibody conjugated to an appropriate fluorophore (dilution 1: 200, duration 1 hour at room temperature), dissolved in 2.5% BSA solution without saponin. Images were obtained by confocal laser scanning microscopy (Zeiss LSM 710) using a 63X oil immersion objective lens. The number of bacteria adhering per cell was calculated by dividing the total number of bacteria attached by the total number of host cells in a single microscopic field. The counting and analysis were done with the help of ZEN Black 2009 software provided by Zeiss.

### Intracellular proliferation assay

The protocol of intracellular proliferation assay has been followed, as demonstrated earlier [45]. Briefly, the seeded RAW264.7, U937, Caco-2, and HeLa cells (1.5 to 2 × 10^5^ cells per well) were infected with STM (WT), *ΔompA, ΔompA*: pQE60-*ompA* and *ΔompA*:pQE60 at MOI 10, as mentioned earlier in this study. After centrifuging the cells at 800 rpm for 5 minutes, the infected cells were incubated at 37°C temperature in the presence of 5% CO_2_ for 25 minutes. Next, the cells were washed thrice with PBS to remove all the unattached extracellular bacteria and subjected to 100 μg/ mL concentrations of gentamycin treatment for 1 hour. After that, the cells were washed thrice with sterile PBS and further incubated with25 μg/ mL concentrations of gentamycin till the lysis. The cells were lysed with 0.1% triton-X-100 at 2 hours and 16 hours post-infection. The lysates were plated on *Salmonella- Shigella* Agar, and the corresponding CFU at 2 hours and 16 hours were determined. The intracellular proliferation of bacteria (Fold proliferation) was determined using a simple formula-

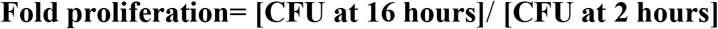

In some sets of experiments, the fold proliferation of STM (WT) and *ΔompA* in the macrophages (RAW 264.7) was measured in the presence of iNOS inhibitor 1400W dihydrochloride [10μM] and activator mouse IFN-γ [100U/ mL]. Both inhibitor (1400W) and activator (IFN-γ) were added to the cells infected with STM (WT) and *ΔompA* along with 25 μg/ mL of gentamycin solution. As usual, the cells were lysed with 0.1% triton-X-100 at 2 hours and 16 hours post-infection. The lysates were plated on *Salmonella- Shigella* Agar, and the corresponding CFU at 2 hours and 16 hours were calculated to determine Fold proliferation. In the intracellular survival assay, two consecutive dilutions were made from each technical replicates at 2 hours and 16 hours. After plating each dilution at 2 hours and 16 hours, the obtained CFU was used to calculate the fold proliferation.

### Chloroquine resistance assay

To estimate the number of intracellular bacteria localized in the cytoplasm of macrophages and epithelial cells, a chloroquine resistance assay was performed using a modified protocol as described previously [64, 65]. Briefly, the seeded RAW264.7 and Caco-2 cells (density- 1.5 to 2 × 10^5^ cells per well) were infected with STM- (WT), *ΔompA,* and *ΔompA*: pQE60-*ompA* at MOI 10, as mentioned earlier in this study. After centrifuging the cells at 800 rpm for 5 minutes, the infected cells were incubated at 37°C temperature in the presence of 5% CO_2_ for 25 minutes. Next, the cells were washed with PBS and subjected to 100 μg/ mL and 25 μg/ mL of gentamycin treatment, respectively. The old DMEM (having 25 μg/ mL gentamycin) was replaced from the wells with freshly prepared DMEM, supplemented with 25 μg/ mL gentamycin and 50 μg/ mL chloroquine two hours before lysis (14 hours post-infection). The cells were lysed with 0.1% triton-X-100 at 16 hours post-infection. The lysates were plated on *Salmonella- Shigella* agar, and the corresponding CFU at 16 hours was determined. Percent abundance of cytosolic and vacuolar bacteria was obtained after dividing the CFU from chloroquine treated set with chloroquine untreated set.

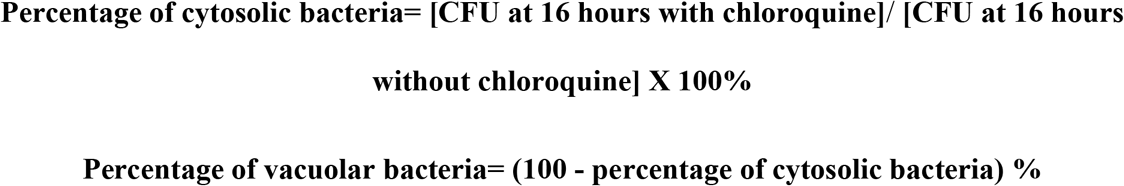

### Confocal microscopy

For immunofluorescence study, RAW 264.7 or Caco-2 cells seeded at a density of 1.5 to 2 × 10^5^ cells per sterile glass coverslip were infected with appropriate bacterial strains at MOI 20. The cells were washed thrice with PBS and fixed with 3.5% paraformaldehyde for 15minutes at indicated time points post-infection. The cells were first incubated with specific primary antibody raised against *Salmonella* SseC/ SseD proteins, mouse lysosome-associated membrane protein-1 (LAMP-1) (rat anti-mouse LAMP-1), mouse nitrotyrosine (mouse anti-mouse nitrotyrosine), and S. Typhimurium (anti- *Salmonella* O antigen) as per the requirements of experiments, diluted in 2.5% BSA and 0.01% saponin (dilution 1: 100, duration 6 to 8 hours at 4°C temperature). This was followed by incubating the cells with appropriate secondary antibodies conjugated with fluorophores (dilution 1: 200, duration 1 hours at room temperature).

The coverslips were mounted with anti-fade reagent and fixed on a glass slide with transparent nail paint. Samples were imaged by confocal laser scanning microscopy (Zeiss LSM 710) using a 63X oil immersion objective lens. The images were analyzed with ZEN Black 2009 software provided by Zeiss.

### Griess assay to measure extracellular nitrite concentration

Extracellular nitrite from infected macrophage cells was measured using a protocol described earlier [66]. 3.13 μM, 6.25 μM, 12.5 μM, 25 μM, 50 μM, 100 μM standard NaNO_2_ solutions were prepared from main stock [0.1 (M) NaNO_2_] by serial dilution in deionized distilled water. The [OD]_545nm_ of the standard solutions were measured after adding and incubating them with Griess reagent, and the standard curve was drawn. Culture supernatants were collected from RAW264.7 cells infected with STM- (WT), *ΔompA, ΔompA*: pQE60-*ompA*, *ΔompA*: pQE60, and heat-killed bacteria at 16 hours post-infection and subjected to nitrite estimation by adding Griess reagents. To 50 μL of culture supernatant, 50 μL of 1% sulphanilamide (made in 5% phosphoric acid), and 50 μL of 0.1% NED (N-1-naphthyl ethylene diamine dihydrochloride) were added and incubated for 10 minutes in darkness at room temperature. The [OD]_545nm_ was measured within 30 minutes of the appearance of a purple-colored product.

### Measurement of intracellular nitric oxide

The level of intracellular nitric oxide of infected macrophages was measured using cell membrane-permeable fluorescent nitric oxide probe 4, 5- diaminofluorescein diacetate (DAF2- DA) [67]. Briefly, RAW264.7 cells were infected with STM- (WT), *ΔompA, ΔompA*: pQE60- *ompA*, *ΔompA ΔompC*, *ΔompA ΔompD*, *ΔompA ΔompF* & LLO at MOI 10 as described before. 16 hours post-infection, the culture supernatants were replaced with DMEM media, supplemented with 5µM concentration of DAF2-DA, followed by further incubation of the infected cells at 370C temperature in the presence of 5% CO_2_ for 30 minutes. The cells were washed with sterile PBS and acquired immediately for analysis by flow cytometry (BD FACSVerse by BD Biosciences-US) using a 491 nm excitation channel and 513 nm emission channel.

### Measurement of the activity of the *spiC* promoter

The activity of *spiC* promoter in STM (WT) and *ΔompA* was measured by mild alteration of a protocol described earlier [66]. Briefly, 1.5 mL of overnight grown stationary phase culture of STM (WT) and *ΔompA* carrying pHG86 *spiC-lacZ* construct were centrifuged at 6000 rpm for 10 minutes, and the pellet was resuspended in 500 µL of Z-buffer (Na2HPO4, 60 mM; NaH2PO4, 40 mM; KCl, 10 mM; MgSO4.7H2O, 1mM). The OD of the Z-buffer was measured at 600 nm after resuspension. The cells were permeabilized by adding 5 µL of 0.1% SDS and 20 µL of chloroform and incubated at room temperature for 5 minutes. 100 µL of 4 mg/ mL of o-nitrophenyl *β*-D galactopyranoside was added in the dark and incubated till the color appeared. The reaction was stopped using 250 µL 1 M Na_2_CO_3_. The reaction mixture was centrifuged at 6000 rpm for 10 minutes, and the OD of the supernatant was measured at 420 nm and 550 nm on flat bottom transparent 96 well plates. STM (WT) and *ΔompA* harboring promoter-less empty pHG86 *LacZ* construct were used as control. The activity of the *spiC* promoter was measured in Miller Unit using the following formula

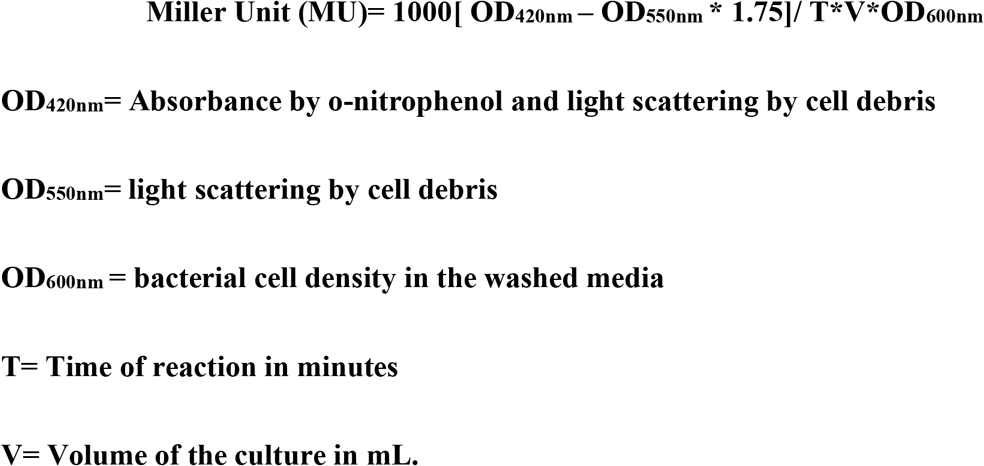

### Measurement of the activity of the *spiC* promoter from intracellular bacteria

RAW264.7 cells were infected with STM (WT): pHG86 *spiC-LacZ* and *ΔompA*: pHG86 *spiC- LacZ* at MOI 50. After centrifuging the cells at 800 rpm for 5 minutes, the infected cells were incubated at 37°C temperature in the presence of 5% CO_2_ for 25 minutes. Next, the cells were washed thrice with PBS to remove all the unattached extracellular bacteria and subjected to 100 μg/ mL and 25 μg/ mL concentrations of gentamycin treatment for 1 hour each. The cells were lysed with 0.1% triton-X-100 at 12 hours post-infection. The lysates were centrifuged at 14,000 rpm for 30 minutes, and the pellet was resuspended in 500 µL of Z-buffer. The OD of the Z-buffer was measured at 600 nm after resuspension. The cells were permeabilized by adding 5 µL of 0.1% SDS and 20 µL of chloroform and incubated at room temperature for 5 minutes. 100 µL of 4 mg/ mL of o-nitrophenyl *β*-D galactopyranoside was added in the dark and incubated till the color appeared. The reaction was stopped using 250 µL 1 M Na_2_CO_3_. The reaction mixture was centrifuged at 6000 rpm for 10 minutes, and the OD of the supernatant was measured at 420 nm and 550 nm on flat bottom transparent 96 well plates. The activity of the *spiC* promoter was measured in Miller Unit using the formula mentioned earlier.

### Measurement of the cytosolic acidification of the bacteria using BCECF-AM

The acidification of the cytosol of the bacteria in the presence of *in vitro* acidic stress was measured using a cell-permeable dual excitation ratiometric dye called 2’,7’-Bis-(2- Carboxyethyl)-5-(and-6)-Carboxyfluorescein, Acetoxymethyl Ester (BCECF-AM). 4.5 × 10^7^ CFU of STM (WT), *ΔompA,* and *ΔompA*: pQE60-*ompA* from 12 hours old overnight grown stationary phase culture was resuspended in phosphate buffer (of pH 5.5, 6, 6,5, and 7 respectively) separate microcentrifuge tubes and incubated for 2 hours in a shaker incubator at 37°C temperature. 30 minutes before the flow cytometry analysis, BCECF-AM (1mg/ mL) was added to each tube to make the final concentration 20 µM. At the end of the incubation period of 2 hours, the bacterial cells were analyzed in flow cytometry (BD FACSVerse by BD Biosciences-US) using 405 nm and 488 nm excitation and 535 nm emission channel. The median fluorescence intensity (MFI) of the bacterial population at 488 nm and 405 nm was obtained from BD FACSuite software. The 488/ 405 ratio was determined to estimate the level of acidification of the bacterial cytosol.

### Measurement of extracellular H_2_O_2_ by phenol red assay

H_2_O_2_ produced by RAW264.7 cells infected with STM- (WT), *ΔompA, ΔompA*: pQE60-*ompA* at MOI 10 was measured by modifying a protocol of phenol red assay as demonstrated before [68]. Briefly, two hours post-infection, infected RAW264.7 cells were supplemented with phenol red solution having potassium phosphate (0.01 M; pH 7.0), glucose (0.0055 M), NaCl (0.14 M), phenol red (0.2 g/ L), and HRPO (8.5 U/ mL; Sigma- Aldrich). 16 hours post-infection, the culture supernatant was collected and subjected to the [OD] measurement at 610 nm wavelength in TECAN 96 well microplate reader. In the presence of H_2_O_2_ produced by macrophages, horseradish peroxidase (HRPO) converts phenol red into a compound that has enhanced absorbance at 610 nm. The concentration of H_2_O_2_ produced by macrophages was measured using a standard curve of H_2_O_2_ in phenol red solution with known concentrations ranging from 0.5 to 5 µM.

### Measurement of intracellular ROS

The level of intracellular ROS in infected macrophages was measured using membrane-permeable redox-sensitive probe 2’,7’- dichlorodihydrofluorescein diacetate (H_2_DCFDA) [68]. Upon its oxidation by intracellular esterases, this non-fluorescent dye is converted into highly fluorescent 2’,7’- Dichlorofluorescein (H_2_DCF), which has emission at 492- 495 nm and excitation at 517 to 527 nm. Briefly, RAW264.7 cells were infected with STM- (WT), *ΔompA, ΔompA*: pQE60-*ompA* at MOI 10 as described before. 16 hours post-infection, the culture supernatants were replaced with DMEM media, supplemented with 10 µM concentration of H_2_DCFDA, followed by further incubation of the infected cells at 370C temperature in the presence of 5% CO_2_ for 30 minutes. The cells were washed with sterile PBS and acquired immediately for analysis by flow cytometry (BD FACSVerse by BD Biosciences-US) using a 492 nm excitation channel and 517 nm emission channel.

### Sensitivity assay of bacteria against *in vitro* nitrosative and oxidative stress

The sensitivity of STM (WT) and *ΔompA* was tested against *in vitro* nitrosative and oxidative stress. H_2_O_2_ dissolved in PBS of pH 5.4 was used for creating *in vitro* oxidative stress. Acidified nitrite (NaNO_2_ in PBS of pH 5.4) alone and a combination of acidified nitrite and H_2_O_2_ were used to generate *in vitro* nitrosative stress [32]. Sensitivity was checked in both concentration and time-dependent manner.

### Concentration-dependent sensitivity

10^8^ CFU of overnight grown stationary phase cultures of STM- (WT) and *ΔompA* were inoculated in varying concentrations of acidified nitrite and peroxide ranging from 200 µM to 5 mM and further incubated for 12 hours. At the end of the incubation period, supernatants from each concentration were collected, serially diluted, plated on *Salmonella- Shigella* agar and the log_10_[CFU/ mL] values were acquired to determine the inhibitory concentrations of nitrite, peroxide, and both together.

### Time-dependent sensitivity

10^8^ CFU of overnight grown stationary phase cultures of STM- (WT) and *ΔompA* were inoculated in 800µM concentration of acidified nitrite and further incubated for 12 hours. Aliquots were collected, serially diluted, and plated on *Salmonella- Shigella* agar at 0, 3, 6, 9, 12 hours post-inoculation to monitor the CFU.

### Bacterial cell viability assay by resazurin

Bacterial cell viability under the treatment of acidified nitrite and peroxide was measured using resazurin assay. Resazurin, a blue-colored non-fluorescent redox indicator, is reduced into resorufin, a pink-colored fluorescent compound (having excitation at 540 nm and emission at 590 nm) by aerobic respiration of metabolically active cells. Briefly, STM- (WT) and *ΔompA* were treated with varying concentrations of acidified nitrite and peroxide, as mentioned above. At the end of the incubation period, supernatants were collected and incubated with resazurin (1 µg/ mL) in a 37°C shaker incubator at 180 rpm for 2 hours in a 96 well plate. At the end of the incubation period, the fluorescence intensity was measured using TECAN 96 well microplate reader, and percent viability was calculated.

### Nitrite uptake assay

Nitrite uptake by different bacterial strains [STM- (WT), *ΔompA*, *ΔompA*: pQE60-*ompA*, *ΔompA*:pQE60, *ΔompAΔompC*, *ΔompAΔompD*, *ΔompAΔompF*, *ΔompC*, *ΔompD*, *ΔompF,* and PFA fixed dead bacteria] was determined by measuring the remaining concentration of nitrite in the uptake mixture using a protocol described earlier [66]. Briefly, 10^8^ CFU of overnight grown stationary phase bacterial cultures were inoculated in an uptake mixture consisting of 40 mM glucose, 80 mM MOPS-NaOH buffer (pH= 8.5), and nitrite (50/ 100/ 200 µM) in a final volume of 5 mL. The assay mixtures were kept in a 37°C shaker incubator after the inoculation was done. At indicated time points, 150 µL of suspension from each assay mixture was collected and subjected to Griess assay to determine the level of remaining nitrite, as mentioned earlier.

### Examination of *in vitro* redox homeostasis of STM (WT) and *ΔompA* in response to acidified nitrite

Stationary phase cultures of STM- (WT) and *ΔompA* harboring pQE60-Grx1-roGFP2 plasmid were sub-cultured in freshly prepared 5 mL LB broth at 1: 33 ratios in the presence of appropriate antibiotic in 37°C shaker incubator at 175 rpm. Once the [OD]_600 nm_ has reached 0.3 to 0.4, 500 µM of IPTG (Sigma-Aldrich) was added, and the cells were further grown at 30°C temperature at 175 rpm for 10 to 12 hours. At the end of the incubation period, 4.5 × 10^7^ CFU of bacteria were subjected to the treatment of acidified nitrite [varying concentrations (as mentioned in the figure legend) of NaNO_2_ in PBS of pH= 5.4] for 15, 30, 45, and 60 minutes. At the end of every indicated time point, the cells were analyzed in flow cytometry (BD FACSVerse by BD Biosciences-US) using 405 nm and 488 nm excitation and 510 nm emission channel. The mean fluorescence intensity at 405 nm and 488 nm was obtained from the FITC positive (GFP expressing) population, and the 405/ 488 ratio was determined.

### Determination of outer membrane porosity of intracellular and extracellular bacteria by bisbenzimide

The outer membrane porosity of STM- (WT), *ΔompA*, *ΔompA*: pQE60-*ompA*, and *ΔompA*:pQE60 grown in low magnesium acidic N s medium (pH= 5.4) was measured using bisbenzimide (Sigma-Aldrich) by modifying a protocol as specified previously [69]. The bacterial strains were grown in low magnesium acidic F medium for 12 hours. At the end of the incubation period, the culture supernatants were collected, and the [OD]_600 nm_ was adjusted to 0.1 with sterile PBS. 20 µL of bisbenzimide (10 µg/ mL) solution was added to 180 µL of culture supernatants (whose [OD]_600 nm_ has already been adjusted to 0.1) in 96 well a microplate and further incubated for 10 minutes in 37°C shaker incubator. Because of enhanced outer membrane porosity, when bisbenzimide is taken up by the bacterial cells, it binds to the bacterial DNA and starts fluorescing. The fluorescence intensity of DNA bound bisbenzimide was measured in TECAN 96 well microplate reader using 346 nm excitation and 460 nm emission filter.

To check the outer membrane porosity of intracellular STM- (WT) and *ΔompA*, infected RAW264.7 macrophage cells were lysed with 0.1% Triton X-100. The lysate was collected and centrifuged at 300g for 5 minutes to settle down eukaryotic cell debris. The sup was collected and further centrifuged at 5000 rpm for 20 minutes to settle down the bacteria. This was followed by decanting the sup and resuspending the bacterial pellets with PBS. Finally, the suspension was subjected to bisbenzimide treatment to measure the fluorescence intensity as mentioned above.

### Determination of bacterial membrane depolarization using DiBAC_4_

Outer membrane depolarization of STM (WT), *ΔompA*, *ΔompA*: pQE60-*ompA*, *ΔompA*:pQE60, STM (WT): pQE60, STM (WT): pQE60-*ompA*, STM (WT): pQE60-*ompC*, STM (WT): pQE60-*ompD*, and STM (WT): pQE60-*ompF* grown in low magnesium acidic F medium (pH= 5.4) for 12 hours was measured using a fluorescent membrane potential sensitive dye called bis-(1,3-dibutyl barbituric acid)-trimethylene oxonol (Invitrogen). Briefly, 4.5 × 10^7^ CFU of each bacterial strain was incubated with 1 µg/ml of DiBAC_4_ for 15 minutes in a 37°C shaker incubator. The DiBAC_4_ treated bacterial cells were further analyzed by flow cytometry (BD FACSVerse by BD Biosciences-US) to evaluate the change in membrane depolarization upon knocking out *ompA*.

Stationary phase cultures of STM (WT), STM (WT): pQE60, STM (WT): pQE60-*ompA*, STM (WT): pQE60-*ompC*, STM (WT): pQE60-*ompD*, and STM (WT): pQE60-*ompF* were sub- cultured in freshly prepared 5 mL LB broth at 1: 33 ratios in the presence of appropriate antibiotics in 37°C shaker incubator at 175 rpm. Once the [OD]_600 nm_ has reached 0.3 to 0.4, 500 µM of IPTG (Sigma-Aldrich) was added, and the cells were further grown at 30°C temperature at 175 rpm for 10 to 12 hours. At the end of the incubation period, 4.5 × 10^7^ CFU of bacteria were subjected to the treatment of 1 µg/mL of DiBAC_4_ for 15 minutes in a 37°C shaker incubator. At the end of the incubation period, the cells were analyzed in flow cytometry (BD FACSVerse by BD Biosciences-US) to measure the outer membrane depolarization.

### Expression profiling of *ompC*, *ompD*, *ompF* in STM- (WT), *ΔompA* and complement strains growing in LB broth, acidic F media, and macrophages

Overnight grown stationary phase cultures of STM- (WT), *ΔompA,* & *ΔompA*: pQE60-*ompA* were inoculated in freshly prepared LB broth, low magnesium acidic F media (pH= 5.4) at a ratio of 1: 100. The cells were further grown in a 37°C shaker incubator at 180 rpm for 12 hours. RAW264.7 cells were infected with above mentioned bacterial strains at MOI 50 and incubated further for 12 hours, as mentioned earlier. At the end of the specified incubation, period RNA was isolated, cDNA was synthesized, and the expression of *ompC, ompD, ompF* were checked, as mentioned earlier.

### Live dead assay by propidium iodide

10^8^ CFU of overnight grown stationary phase cultures of STM (WT): pQE60, STM (WT): pQE60-*ompA*, STM (WT): pQE60-*ompC*, STM (WT): pQE60-*ompD*, STM (WT): pQE60- *ompF* were inoculated in 1 mM concentration of acidified nitrite (total volume 10mL) and further incubated for 12 hours. 300 µL of aliquots (corresponding to 10^5^ to 10^6^ CFU of bacteria) were collected 12 hours post-inoculation and subjected to the treatment with propidium iodide (PI) (Sigma-Aldrich) (concentration- 1µg/ mL) for 30 minutes at 37°C temperature. After the incubation, the PI-treated bacterial samples were analyzed by flow cytometry (BD FACSVerse by BD Biosciences-US) to estimate the percent viability.

### Animal survival assay

4-6 weeks old BALB/c and C57BL/6 mice housed in the specific-pathogen-free condition of central animal facility of Indian Institute of Science, Bangalore was used for all the *in vivo* infection and survival studies. The Institutional Animal Ethics Committee approved all the animal experiments, and the National Animal Care Guidelines were strictly followed. Two cohorts of twenty 4-6 weeks old BALB/c and C57BL/6 mice were infected with 10-12 hours old overnight grown stationary phase cultures of STM (WT) and *ΔompA* by oral gavaging at lethal dose 10^8^ CFU/ animal respectively **(n= 10)**. The survival of infected mice was observed for the next few days until all the mice infected with STM (WT) died. The survival was recorded, and the data was represented as percent survival.

### Determination of bacterial burden in different organs

Four cohorts of five 4-6 weeks old C57BL/6 mice were infected with STM (WT) and *ΔompA* by oral gavaging at sub-lethal dose 10^7^ CFU/ animal, respectively **(n= 5)**. Two of these cohorts infected with STM- (WT) and *ΔompA* strains respectively were further intraperitoneally injected with iNOS inhibitor aminoguanidine hydrochloride (AGH- 10mg/ kg of body weight) regularly for five days post-infection. The other two cohorts were treated with a placebo. Two cohorts of *iNOS^-/-^* mice were orally infected with STM (WT) and *ΔompA* at 10^7^ CFU/ animal **(n=5)**. Two cohorts of five *gp91phox^-/-^* mice unable to generate ROS were orally gavaged with STM- (WT) and *ΔompA* at 10^7^ CFU/ animal **(n=5)**. On the 5^th^ day post-infection, all the mice were sacrificed, followed by isolation, weighing, and homogenization of specific organs like- liver, spleen, and MLN. The organ lysates were plated on *Salmonella Shigella* agar to determine the bacterial burden in different organs. The CFU corresponds to an organ was normalized with organ weight and the log_10_[CFU/ gm-wt.] value has been plotted.

### Statistical analysis

Each assay has been independently repeated 2 to 5 times [as mentioned in the figure legends]. The *in vitro* data and the results obtained from cell line experiments were analyzed by unpaired student’s *t*-test, and *p* values below 0.05 were considered significant. Results received from *in vitro* sensitivity assays were analyzed by 2way ANOVA. Data obtained from *in vivo* infection of mice were analyzed by Mann- Whitney *U* test from GraphPad Prism 8.4.3 (686) software. Flow cytometry data were analyzed and plotted using BD FACSuite (by BD Biosciences-US) and CytoFLEX (by Beckman Coulter Life Sciences) software. The results are expressed as mean ± SD or mean ± SEM. Differences between experimental groups were considered significant for *p*< 0.05.

## Acknowledgments

Departmental Confocal Facility, Departmental Real-Time PCR Facility, Central Bioimaging Facility, Central Flowcytometry Facility, Department of Biochemistry Flowcytometry Facility, and Central Animal Facility at IISc are duly acknowledged. Ms. Anusha and Ms. Navya are acknowledged for their help in image acquisition. Mrs. Ranjitha is acknowledged for her help in flow cytometry data acquisition. Dr. Amit Singh (Centre for Infectious Disease Research and Department of Microbiology and Cell Biology, Indian Institute of Science, Bangalore) is acknowledged for providing pQE60-Grx1-roGFP2 construct. Ms. Debapriya Mukherjee and Ms. Dipasree Hajra are recognized for their timely technical support.

## Availability of data and materials

All data generated and analyzed during this study, including the supplementary information files, have been incorporated in this article. The data is available from the corresponding author on reasonable request.

## Author Contributions

ARC and DC conceived the study and designed the experiments. ARC performed all the experiments, analyzed the data, and wrote the original draft of the manuscript. SS constructed pQE60-*ompA* recombinant plasmid under the supervision of UV. SS, UV, and DC reviewed and edited the manuscript. DC supervised the study. All the authors have read and approved the manuscript.

## Declarations

### Ethics statement

All the animal experiments were approved by the Institutional Animal Ethics Committee, and the Guidelines provided by National Animal Care were strictly followed. (Registration No: 48/1999/CPCSEA).

### Consent for publication

Not applicable.

## Supplementary Figures

**Figure S1.**
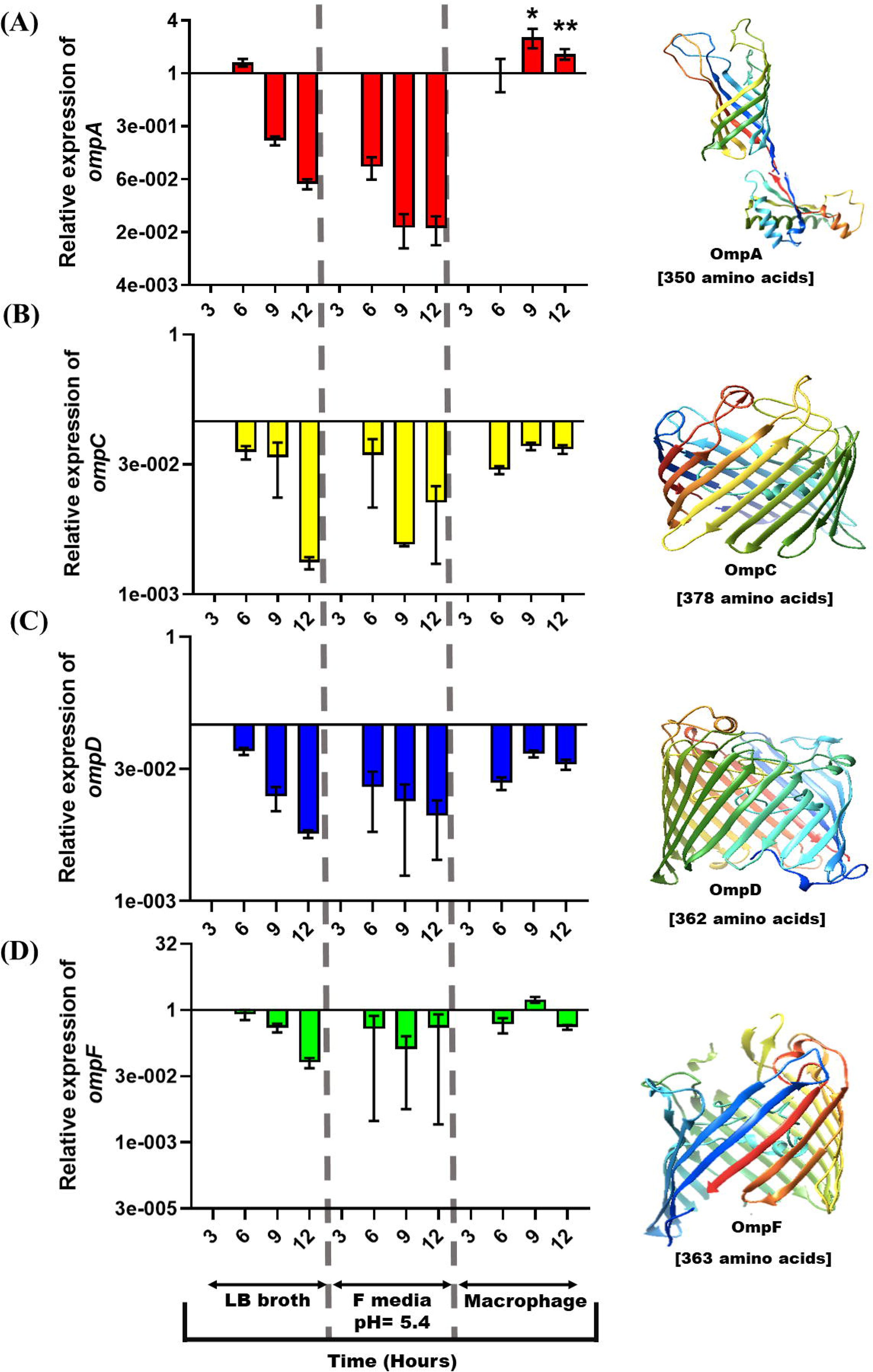
OmpA plays a crucial role in the survival of *Salmonella* Typhimurium in murine macrophages. The transcript level expression profile of (A) *ompA*, (B) *ompC*, (C) *ompD*, and (D) *ompF* in STM- (WT) at indicated time points (3, 6, 9, 12 hours) in LB broth (n=3, N=3), low magnesium acidic F media (pH=5.4) (n=3, N=3), and RAW264.7 murine macrophage cells (MOI= 50), (n=3, N=3). The time-dependent relative expression of *ompA, ompC, ompD, ompF* have been represented in the log2 scale. The predicted structures of porins (A) OmpA, (B) OmpC, (C) OmpD, and (D) OmpF using SWISS-MODEL protein structure homology-modeling server. ***(P)* *< 0.05, *(P)* **< 0.005, (Student’s *t* test- unpaired).**

**Figure S2.**
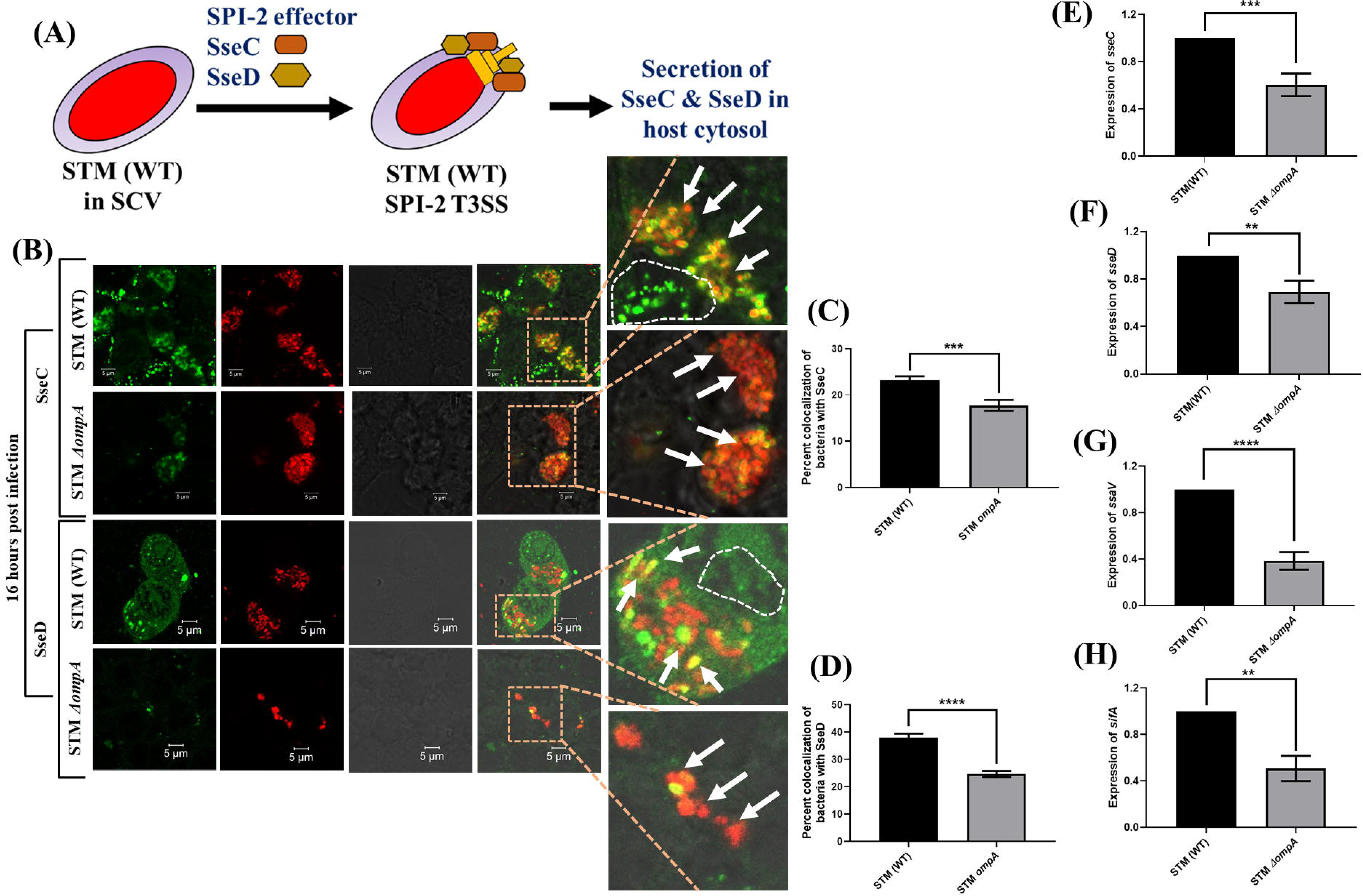
STM *ΔompA* comes out into the cytoplasm of the macrophages after quitting the vacuole. (A) The schematic representation of the SPI-2 encoded T3SS and the secreted effector proteins SseC and SseD around intracellular wild-type *Salmonella*. (B-D) RAW264.7 cells were infected with STM (WT)-: RFP, *ΔompA*: RFP, at MOI 20. Cells were fixed at 16 hours post- infection, *Salmonella* SPI-2 encoded translocon proteins SseC and SseD were labeled with anti- *Salmonella* SseC/ SseD antibody. (C), (D) The quantification of SseC and SseD recruitment on bacteria in RAW 264.7 cells, respectively. Percent colocalization between the bacteria and the effector was determined after analyzing 50 different microscopic stacks from three independent experiments. Scale bar = 5μm, (n=50, N=3). (E-H) The transcript level expression of (E) *sseC*, (F) *sseD*, (G) *ssaV*, and (H) *sifA* in STM (WT) and *ΔompA* growing intracellularly in RAW264.7 cells 12 hours post-infection, (n=3, N=3). ***(P)* **< 0.005, *(P)* ***< 0.0005, *(P)* ****< 0.0001, ns= non-significant, (Student’s t test- unpaired).**

**Figure S3.**
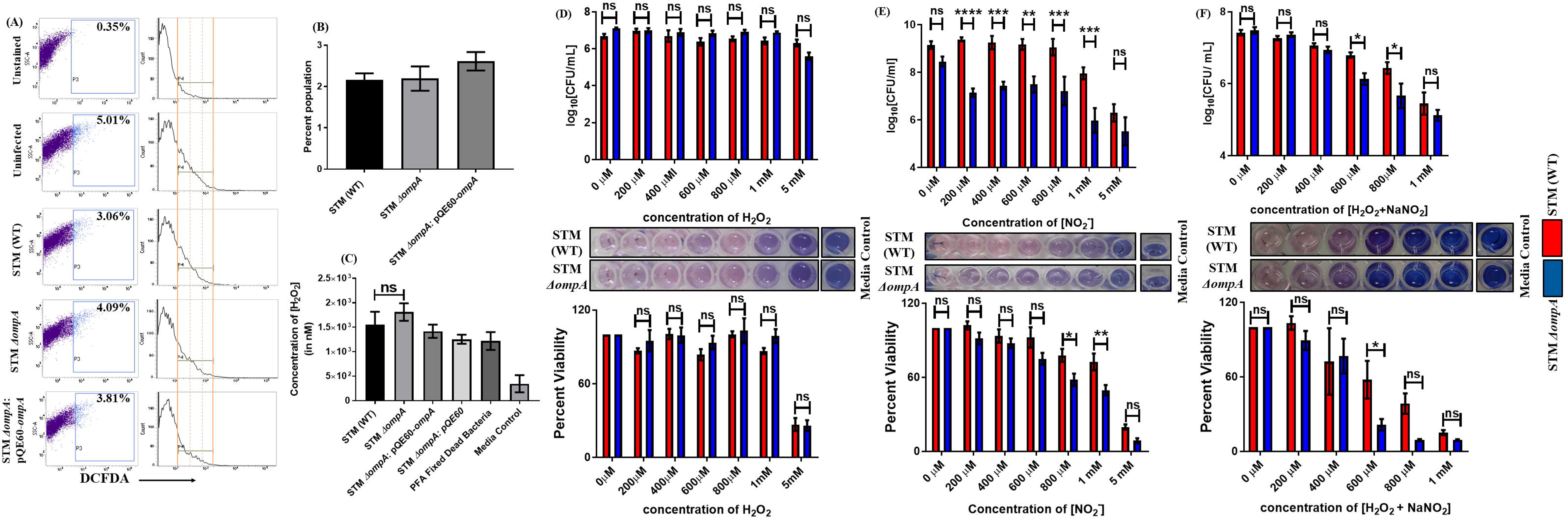
OmpA does not have a significant role in building up *in vitro* and *in vivo* protection of *Salmonella* against oxidative stress. (A) Estimation of the level of intracellular reactive oxygen species (ROS) in RAW 264.7 cells infected with STM- (WT), *ΔompA*, and *ΔompA*: pQE60-*ompA* at MOI 10, 16 hours post- infection using DCFDA [10 µM] by flow cytometry. Unstained and uninfected RAW264.7 cells have also been used as controls. Both dot plots (SSC-A vs. DCFDA) and histograms (Count vs. DCFDA) have been represented. (B) The percent population of DACFDA positive RAW264.7 cells, (n=4, N=3). (C) Estimation of extracellular ROS from the culture supernatant of RAW264.7 cells infected with STM- (WT), *ΔompA*, *ΔompA*: pQE60-*ompA*, *ΔompA*: pQE60, & PFA fixed dead bacteria respectively at MOI 10. 2 hours post-infection, the cells were supplemented with phenol red solution having horseradish peroxidase. 16 hours post-infection, the culture supernatants were collected, and the OD was measured at 610 nm (n=3, N=2). (D) Checking the *in vitro* sensitivity of STM- (WT) and *ΔompA* in the presence of H_2_O_2_ by serial dilution, plating, and CFU calculation and resazurin test. 10^8^ CFU of overnight grown stationary phase culture of STM (WT) and *ΔompA* were inoculated in PBS with varying concentrations of H_2_O_2_. 12 hours post-inoculation, the supernatants were collected for plating on SS agar to determine the log_10_[CFU/mL] (N=3) and resazurin test to determine percent viability. (n=3, N=3). (E) Checking the *in vitro* sensitivity of STM– (WT) and *ΔompA* in the presence of acidified nitrite by serial dilution, plating, and CFU calculation and resazurin test. 10^8^ CFU of overnight grown stationary phase culture of STM (WT) and *ΔompA* were inoculated in PBS (pH=5.4) with varying concentrations of NaNO_2_. 12 hours post-inoculation, the supernatants were collected for plating on SS agar to determine the log_10_[CFU/mL], (N=3) and resazurin test to determine percent viability. (n=3, N=3). (F) Checking the *in vitro* sensitivity of STM– (WT) and *ΔompA* in the presence of NaNO_2_ and H_2_O_2_ combined, by serial dilution, plating, and CFU calculation and resazurin test. 10^8^ CFU of overnight grown stationary phase culture of STM (WT) and *ΔompA* were inoculated in PBS (pH=5.4) with varying concentrations of NaNO_2_ and H_2_O_2_. 12 hours post-inoculation, the supernatants were collected for plating on SS agar to determine the log_10_[CFU/mL] (N=3) and resazurin test to determine percent viability (n=3, N=3). ***(P)* *< 0.05, *(P)* **< 0.005, *(P)* ***< 0.0005, *(P)* ****< 0.0001, ns= non-significant, (2way ANOVA), (Student’s *t* test- unpaired).**

**Figure S4.**
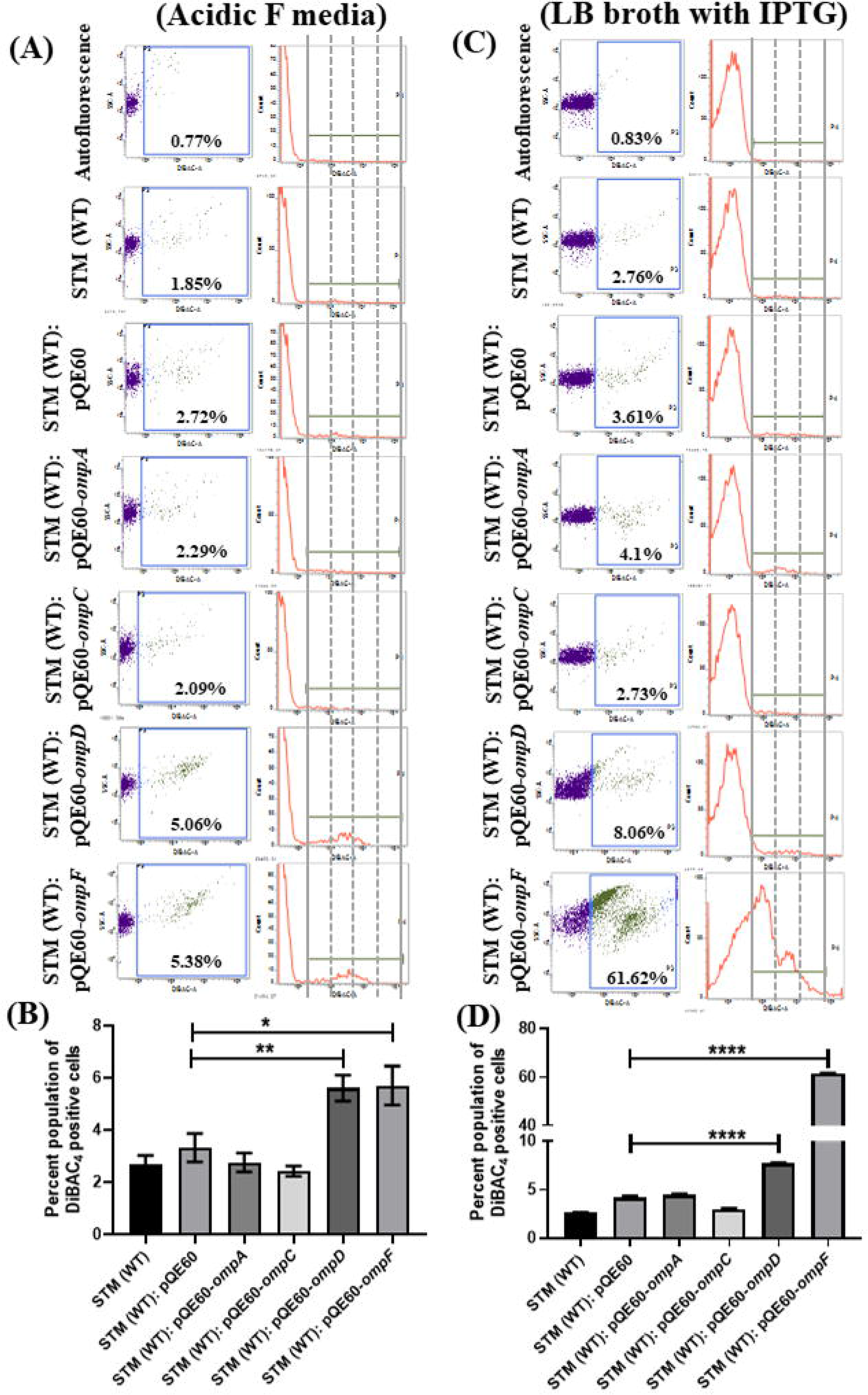
Over-expression of *ompF* in wild-type *Salmonella* enhances the outer membrane porosity of the bacteria. Measurement of outer membrane porosity of STM (WT), STM (WT): pQE60, STM (WT): pQE60-*ompA*, STM (WT): pQE60-*ompC*, STM (WT): pQE60-*ompD*, and STM (WT): pQE60- *ompF* in (A) acidic F media and (C) LB broth with 500 µM of IPTG [12 hours post inoculation] using DiBAC_4_ (final concentration- 1 µg/ mL) by flow cytometry. Unstained bacterial cells were used as control (A) and (C). Both dot plots (SSC-A vs. DiBAC_4_) and histograms (Count vs. DiBAC_4_) have been represented. Percent population of DiBAC_4_ positive cells in acidic F media (B) and LB broth culture (D) has been represented here in the form of a bar graph (n=6, N=3 for B and n=6 for D). ***(P)* *< 0.05, *(P)* **< 0.005, *(P)* ***< 0.0005, *(P)* ****< 0.0001, ns= non-significant, (Student’s *t* test- unpaired)**

**Figure S5.**
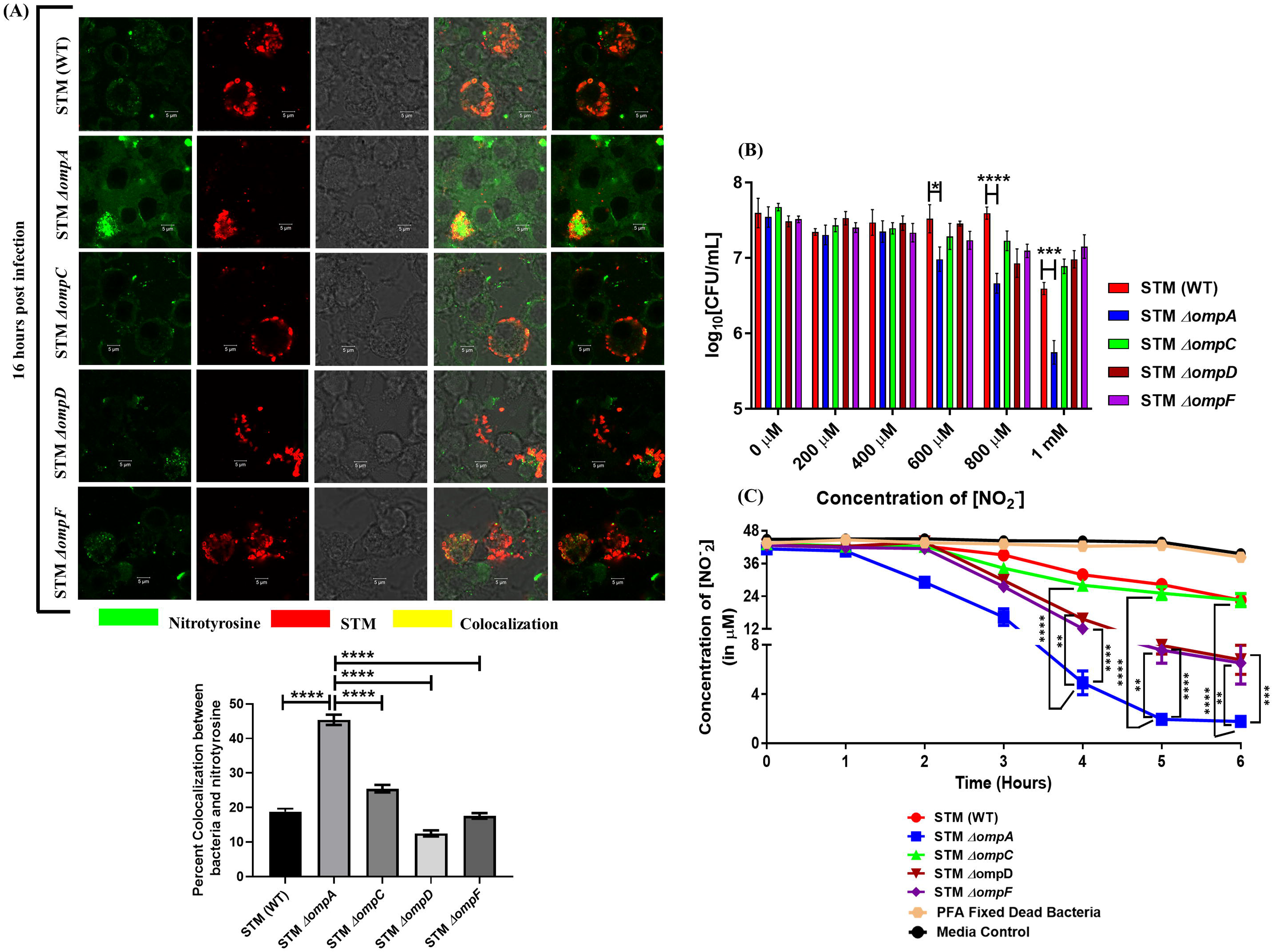
Unlike *ompA*, the deletion of *ompC, ompD,* and *ompF* do not hamper the viability of *Salmonella* against intracellular and extracellular nitrosative stress. (A) RAW264.7 cells were infected with STM- (WT), *ΔompA*, *ΔompC*, *ΔompD*, and *ΔompF* at MOI 20. Cells were fixed at 16 hours post-infection, followed by labeled with anti-*Salmonella* antibody and anti-mouse nitrotyrosine antibody, respectively. Quantification of nitrotyrosine recruitment on STM- (WT), *ΔompA*, *ΔompC*, *ΔompD*, and *ΔompF* has been represented in the form of a graph. Percent colocalization of bacteria with nitrotyrosine has been determined after analyzing more than 60 different microscopic fields from two independent experiments (n≥60, N=3). Scale bar = 5μm. (B) Checking the *in vitro* sensitivity of STM– (WT), *ΔompA*, *ΔompC*, *ΔompD*, and *ΔompF* in the presence of acidified nitrite by serial dilution, plating, and CFU calculation. 10^8^ CFU of overnight grown stationary phase culture of STM (WT), *ΔompA, ΔompC, ΔompD,* and *ΔompF* were inoculated in PBS (pH=5.4) with varying concentrations of NaNO_2_. 12 hours post-inoculation, the supernatants were collected for plating on SS agar to determine the log_10_[CFU/mL], (N=3). (C) *In vitro* nitrite uptake assay of STM- (WT), *ΔompA*, *ΔompC, ΔompD, ΔompF* & PFA fixed dead bacteria. 10^8^ CFU of overnight grown stationary phase culture of all the strains were inoculated in MOPS- NaOH buffer (pH= 8.5) with an initial nitrite concentration of 50 µM. The remaining nitrite concentration of the media was determined by Griess assay at 0, 1, 2, 3-, 4-, 5-, and 6-hours post-inoculation (n=3, N=3). *(P)* *< 0.05, *(P)* **< 0.005, *(P)* ***< 0.0005, *(P)* ****< 0.0001, ns= non-significant, (2way ANOVA), (Student’s *t* test).

**Figure S6.**
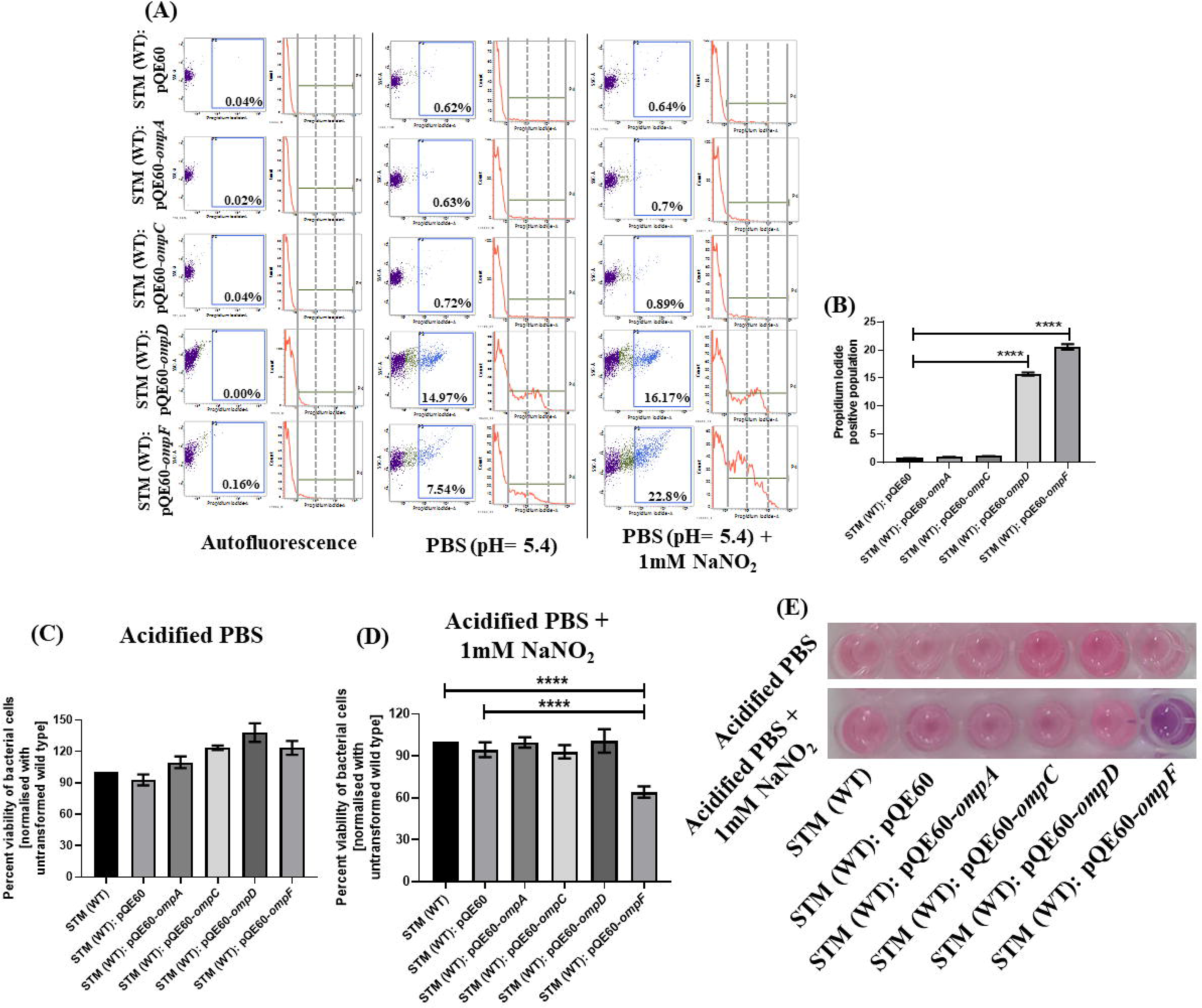
Overexpression of *ompF* enhances the susceptibility of wild-type *Salmonella* towards *in vitro* nitrosative stress. Measurement of *in vitro* viability of STM (WT), STM (WT): pQE60, STM (WT): pQE60- *ompA*, STM (WT): pQE60-*ompC*, STM (WT): pQE60-*ompD*, and STM (WT): pQE60-*ompF* in acidified nitrite (PBS of pH= 5.4 and 1 mM NaNO_2_ at 12 hours post inoculation using propidium iodide (final concentration- 1 µg/ mL) by flow cytometry (A). Unstained bacterial cells were used as control. Bacterial cells grown in acidified PBS were stained with propidium iodide to determine the bacterial death contributed by *in vitro* reactive nitrogen intermediates. Both dot plots (SSC-A vs. DiBAC_4_) and histograms (Count vs. DiBAC_4_) have been represented. Percent population of propidium iodide positive cells from acidified nitrite have been represented here in the form of bar graph (n=8, N=2). Measurement of *in vitro* viability of STM (WT), STM (WT): pQE60, STM (WT): pQE60-*ompA*, STM (WT): pQE60-*ompC*, STM (WT): pQE60-*ompD*, and STM (WT): pQE60-*ompF* in acidified PBS (C and E) and acidified nitrite (PBS of pH= 5.4 and 1 mM NaNO_2_) (D and E) at 12 hours post inoculation using resazurin (final concentration- 0.002 mg/ mL) (n=6 for C and n=8 for D). ***(P)* *< 0.05, *(P)* **< 0.005, *(P)* ***< 0.0005, *(P)* ****< 0.0001, ns= non-significant, (Student’s *t* test- unpaired).**

## References

1. Vergalli, J., et al., Porins and small-molecule translocation across the outer membrane of Gram-negative bacteria. Nat Rev Microbiol, 2020. 18(3): p. 164–176.

2. March, C., et al., Klebsiella pneumoniae outer membrane protein A is required to prevent the activation of airway epithelial cells. J Biol Chem, 2011. 286(12): p. 9956–67.

3. Hejair, H.M.A., et al., Functional role of ompF and ompC porins in pathogenesis of avian pathogenic Escherichia coli. Microb Pathog, 2017. 107: p. 29–37.

4. Krishnan, S. and N.V. Prasadarao, Outer membrane protein A and OprF: versatile roles in Gram- negative bacterial infections. FEBS J, 2012. 279(6): p. 919–31.

5. Wu, X.B., et al., Outer membrane protein OmpW of Escherichia coli is required for resistance to phagocytosis. Res Microbiol, 2013. 164(8): p. 848–55.

6. Li, W., et al., Contribution of the outer membrane protein OmpW in Escherichia coli to complement resistance from binding to factor H. Microb Pathog, 2016. 98: p. 57–62.

7. Park, J.S., et al., Mechanism of anchoring of OmpA protein to the cell wall peptidoglycan of the gram-negative bacterial outer membrane. FASEB J, 2012. 26(1): p. 219–28.

8. Ipinza, F., et al., Participation of the Salmonella OmpD porin in the infection of RAW264.7 macrophages and BALB/c mice. PLoS One, 2014. 9(10): p. e111062.

9. van der Heijden, J., et al., Salmonella Rapidly Regulates Membrane Permeability To Survive Oxidative Stress. mBio, 2016. 7(4).

10. Eriksson, S., et al., Unravelling the biology of macrophage infection by gene expression profiling of intracellular Salmonella enterica. Mol Microbiol, 2003. 47(1): p. 103–18.

11. Hautefort, I., et al., During infection of epithelial cells Salmonella enterica serovar Typhimurium undergoes a time-dependent transcriptional adaptation that results in simultaneous expression of three type 3 secretion systems. Cell Microbiol, 2008. 10(4): p. 958–84.

12. Deiwick, J., et al., Environmental regulation of Salmonella pathogenicity island 2 gene expression. Mol Microbiol, 1999. 31(6): p. 1759–73.

13. Datsenko, K.A. and B.L. Wanner, One-step inactivation of chromosomal genes in Escherichia coli K-12 using PCR products. Proc Natl Acad Sci U S A, 2000. 97(12): p. 6640–5.

14. Garai, P., et al., Peptide utilizing carbon starvation gene yjiY is required for flagella mediated infection caused by Salmonella. Microbiology (Reading), 2016. 162(1): p. 100–116.

15. Garai, P., D.P. Gnanadhas, and D. Chakravortty, Salmonella enterica serovars Typhimurium and Typhi as model organisms: revealing paradigm of host-pathogen interactions. Virulence, 2012. 3(4): p. 377–88.

16. Knuff, K. and B.B. Finlay, What the SIF Is Happening-The Role of Intracellular Salmonella- Induced Filaments. Front Cell Infect Microbiol, 2017. 7: p. 335.

17. Chakraborty, S., H. Mizusaki, and L.J. Kenney, A FRET-based DNA biosensor tracks OmpR- dependent acidification of Salmonella during macrophage infection. PLoS Biol, 2015. 13(4): p. e1002116.

18. Canton, J., et al., Contrasting phagosome pH regulation and maturation in human M1 and M2 macrophages. Mol Biol Cell, 2014. 25(21): p. 3330–41.

19. Martin-Orozco, N., et al., Visualization of vacuolar acidification-induced transcription of genes of pathogens inside macrophages. Mol Biol Cell, 2006. 17(1): p. 498–510.

20. Choi, J. and E.A. Groisman, Acidic pH sensing in the bacterial cytoplasm is required for Salmonella virulence. Mol Microbiol, 2016. 101(6): p. 1024–38.

21. Chakravortty, D., et al., Formation of a novel surface structure encoded by Salmonella Pathogenicity Island 2. EMBO J, 2005. 24(11): p. 2043–52.

22. Holzer, S.U. and M. Hensel, Functional dissection of translocon proteins of the Salmonella pathogenicity island 2-encoded type III secretion system. BMC Microbiol, 2010. 10: p. 104.

23. Foster, N., S.D. Hulme, and P.A. Barrow, Induction of antimicrobial pathways during early- phase immune response to Salmonella spp. in murine macrophages: gamma interferon (IFN- gamma) and upregulation of IFN-gamma receptor alpha expression are required for NADPH phagocytic oxidase gp91-stimulated oxidative burst and control of virulent Salmonella spp. Infect Immun, 2003. 71(8): p. 4733–41.

24. Umezawa, K., et al., Induction of nitric oxide synthesis and xanthine oxidase and their roles in the antimicrobial mechanism against Salmonella typhimurium infection in mice. Infect Immun, 1997. 65(7): p. 2932–40.

25. van der Heijden, J., et al., Direct measurement of oxidative and nitrosative stress dynamics in Salmonella inside macrophages. Proc Natl Acad Sci U S A, 2015. 112(2): p. 560–5.

26. Beuzon, C.R., S.P. Salcedo, and D.W. Holden, Growth and killing of a Salmonella enterica serovar Typhimurium sifA mutant strain in the cytosol of different host cell lines. Microbiology (Reading), 2002. 148(Pt 9): p. 2705–2715.

27. Chakravortty, D., I. Hansen-Wester, and M. Hensel, Salmonella pathogenicity island 2 mediates protection of intracellular Salmonella from reactive nitrogen intermediates. J Exp Med, 2002. 195(9): p. 1155–66.

28. Ruan, Y., et al., Listeriolysin O Membrane Damaging Activity Involves Arc Formation and Lineaction -- Implication for Listeria monocytogenes Escape from Phagocytic Vacuole. PLoS Pathog, 2016. 12(4): p. e1005597.

29. Uchiya, K.I. and T. Nikai, Salmonella virulence factor SpiC is involved in expression of flagellin protein and mediates activation of the signal transduction pathways in macrophages. Microbiology (Reading), 2008. 154(Pt 11): p. 3491–3502.

30. Uchiya, K. and T. Nikai, Salmonella pathogenicity island 2-dependent expression of suppressor of cytokine signaling 3 in macrophages. Infect Immun, 2005. 73(9): p. 5587–94.

31. Pearse, D.D., et al., Comparison of iNOS inhibition by antisense and pharmacological inhibitors after spinal cord injury. J Neuropathol Exp Neurol, 2003. 62(11): p. 1096–107.

32. Heaselgrave, W., P.W. Andrew, and S. Kilvington, Acidified nitrite enhances hydrogen peroxide disinfection of Acanthamoeba, bacteria and fungi. J Antimicrob Chemother, 2010. 65(6): p. 1207–14.

33. Fang, F.C., Perspectives series: host/pathogen interactions. Mechanisms of nitric oxide-related antimicrobial activity. J Clin Invest, 1997. 99(12): p. 2818–25.

34. Tung, Q.N., et al., Application of genetically encoded redox biosensors to measure dynamic changes in the glutathione, bacillithiol and mycothiol redox potentials in pathogenic bacteria. Free Radic Biol Med, 2018. 128: p. 84–96.

35. Reuter, W.H., et al., Utilizing redox-sensitive GFP fusions to detect in vivo redox changes in a genetically engineered prokaryote. Redox Biol, 2019. 26: p. 101280.

36. Marathe, S.A., S. Ray, and D. Chakravortty, Curcumin increases the pathogenicity of Salmonella enterica serovar Typhimurium in murine model. PLoS One, 2010. 5(7): p. e11511.

37. Smani, Y., et al., Role of OmpA in the multidrug resistance phenotype of Acinetobacter baumannii. Antimicrob Agents Chemother, 2014. 58(3): p. 1806–8.

38. Ghai, I. and S. Ghai, Understanding antibiotic resistance via outer membrane permeability. Infect Drug Resist, 2018. 11: p. 523–530.

39. Kukkonen, M. and T.K. Korhonen, The omptin family of enterobacterial surface proteases/adhesins: from housekeeping in Escherichia coli to systemic spread of Yersinia pestis. Int J Med Microbiol, 2004. 294(1): p. 7–14.

40. Ebbensgaard, A., et al., The Role of Outer Membrane Proteins and Lipopolysaccharides for the Sensitivity of Escherichia coli to Antimicrobial Peptides. Front Microbiol, 2018. 9: p. 2153.

41. Iyer, R., et al., Acinetobacter baumannii OmpA Is a Selective Antibiotic Permeant Porin. ACS Infect Dis, 2018. 4(3): p. 373–381.

42. Choi, U. and C.R. Lee, Distinct Roles of Outer Membrane Porins in Antibiotic Resistance and Membrane Integrity in Escherichia coli. Front Microbiol, 2019. 10: p. 953.

43. Nevermann, J., et al., Identification of Genes Involved in Biogenesis of Outer Membrane Vesicles (OMVs) in Salmonella enterica Serovar Typhi. Front Microbiol, 2019. 10: p. 104.

44. Calderon, I.L., et al., Response regulator ArcA of Salmonella enterica serovar Typhimurium downregulates expression of OmpD, a porin facilitating uptake of hydrogen peroxide. Res Microbiol, 2011. 162(2): p. 214–22.

45. Chaudhuri, D., et al., Salmonella Typhimurium Infection Leads to Colonization of the Mouse Brain and Is Not Completely Cured With Antibiotics. Front Microbiol, 2018. 9: p. 1632.

46. Li, J., et al., ChIP-Seq Analysis of the sigmaE Regulon of Salmonella enterica Serovar Typhimurium Reveals New Genes Implicated in Heat Shock and Oxidative Stress Response. PLoS One, 2015. 10(9): p. e0138466.

47. Mohan Nair, M.K. and K. Venkitanarayanan, Role of bacterial OmpA and host cytoskeleton in the invasion of human intestinal epithelial cells by Enterobacter sakazakii. Pediatr Res, 2007. 62(6): p. 664–9.

48. Mittal, R., et al., Deciphering the roles of outer membrane protein A extracellular loops in the pathogenesis of Escherichia coli K1 meningitis. J Biol Chem, 2011. 286(3): p. 2183–93.

49. Singamsetty, V.K., et al., Outer membrane protein A expression in Enterobacter sakazakii is required to induce microtubule condensation in human brain microvascular endothelial cells for invasion. Microb Pathog, 2008. 45(3): p. 181–91.

50. Abreu, A.G. and A.S. Barbosa, How Escherichia coli Circumvent Complement-Mediated Killing. Front Immunol, 2017. 8: p. 452.

51. Sukumaran, S.K., H. Shimada, and N.V. Prasadarao, Entry and intracellular replication of Escherichia coli K1 in macrophages require expression of outer membrane protein A. Infect Immun, 2003. 71(10): p. 5951–61.

52. Brumell, J.H., et al., Disruption of the Salmonella-containing vacuole leads to increased replication of Salmonella enterica serovar typhimurium in the cytosol of epithelial cells. Infect Immun, 2002. 70(6): p. 3264–70.

53. Beuzon, C.R., et al., Salmonella maintains the integrity of its intracellular vacuole through the action of SifA. EMBO J, 2000. 19(13): p. 3235–49.

54. Boncompain, G., et al., Production of reactive oxygen species is turned on and rapidly shut down in epithelial cells infected with Chlamydia trachomatis. Infect Immun, 2010. 78(1): p. 80–7.

55. Cheng, C., et al., Listeriolysin O Pore-Forming Activity Is Required for ERK1/2 Phosphorylation During Listeria monocytogenes Infection. Front Immunol, 2020. 11: p. 1146.

56. Panday, A., et al., NADPH oxidases: an overview from structure to innate immunity-associated pathologies. Cell Mol Immunol, 2015. 12(1): p. 5–23.

57. Gallois, A., et al., Salmonella pathogenicity island 2-encoded type III secretion system mediates exclusion of NADPH oxidase assembly from the phagosomal membrane. J Immunol, 2001. 166(9): p. 5741–8.

58. Webb, J.L., et al., Macrophage nitric oxide synthase associates with cortical actin but is not recruited to phagosomes. Infect Immun, 2001. 69(10): p. 6391–400.

59. Zielonka, J., et al., Mitigation of NADPH Oxidase 2 Activity as a Strategy to Inhibit Peroxynitrite Formation. J Biol Chem, 2016. 291(13): p. 7029–44.

60. Vlahos, R., et al., Inhibition of Nox2 oxidase activity ameliorates influenza A virus-induced lung inflammation. PLoS Pathog, 2011. 7(2): p. e1001271.

61. Weller, R., et al., Antimicrobial effect of acidified nitrite on dermatophyte fungi, Candida and bacterial skin pathogens. J Appl Microbiol, 2001. 90(4): p. 648–52.

62. Kono, Y., et al., Lactate-dependent killing of Escherichia coli by nitrite plus hydrogen peroxide: a possible role of nitrogen dioxide. Arch Biochem Biophys, 1994. 311(1): p. 153–9.

63. Balakrishnan, A., M. Schnare, and D. Chakravortty, Of Men Not Mice: Bactericidal/Permeability-Increasing Protein Expressed in Human Macrophages Acts as a Phagocytic Receptor and Modulates Entry and Replication of Gram-Negative Bacteria. Front Immunol, 2016. 7: p. 455.

64. Wrande, M., et al., Genetic Determinants of Salmonella enterica Serovar Typhimurium Proliferation in the Cytosol of Epithelial Cells. Infect Immun, 2016. 84(12): p. 3517–3526.

65. Knodler, L.A., V. Nair, and O. Steele-Mortimer, Quantitative assessment of cytosolic Salmonella in epithelial cells. PLoS One, 2014. 9(1): p. e84681.

66. Das, P., et al., Novel role of the nitrite transporter NirC in Salmonella pathogenesis: SPI2- dependent suppression of inducible nitric oxide synthase in activated macrophages. Microbiology (Reading), 2009. 155(Pt 8): p. 2476–2489.

67. Lahiri, A., P. Das, and D. Chakravortty, Arginase modulates Salmonella induced nitric oxide production in RAW264.7 macrophages and is required for Salmonella pathogenesis in mice model of infection. Microbes Infect, 2008. 10(10-11): p. 1166–74.

68. Lahiri, A., P. Das, and D. Chakravortty, The LysR-type transcriptional regulator Hrg counteracts phagocyte oxidative burst and imparts survival advantage to Salmonella enterica serovar Typhimurium. Microbiology (Reading), 2008. 154(Pt 9): p. 2837–2846.

69. Coldham, N.G., et al., A 96-well plate fluorescence assay for assessment of cellular permeability and active efflux in Salmonella enterica serovar Typhimurium and Escherichia coli. J Antimicrob Chemother, 2010. 65(8): p. 1655–63.

